# A multilayered network reveals the centrality of newly discovered *Nucleocytoviricota* in wastewater treatment plant communities

**DOI:** 10.64898/2026.05.21.726523

**Authors:** Dominik Lücking, Alejandro Manzano-Marín, Anouk Willemsen

## Abstract

Viruses of the phylum *Nucleocytoviricota* are paradigm-shifting entities due to their exceptionally large genomes and complex gene repertoires, which blur the lines between viral and cellular life. Previous research has leveraged computational approaches to map their extensive diversity, while experimental work has started to elucidate the intricate networks they form with hosts, bacterial and other symbionts, co-infecting virophages and other mobile genetic elements. Here, we analyzed deeply sequenced metagenomes sampled from wastewater treatment plants in Denmark, an environment with rapid abiotic changes and known to be a hotbed of dense microbial communities. We discovered 61 novel nucleocytoviruses, 15 virophages and 14 polinton-like viruses. By integrating them with microbial contigs into a multilayered interaction network, we explore the role of these entities on a mesocosm scale. We demonstrate the centrality of nucleocytoviruses, positioning them as important players shaping microbial community structure and evolution in wastewater treatment plants.

## Introduction

The phylum of *Nucleocytoviricota* comprises large double-stranded DNA viruses. Within this phylum, so-called “giant viruses” display astonishing characteristics, including species with large genomes (up to 2.5 Mb; Philippe et al., 2013) and giant virions of up to 2 µm (Legendre et al., 2014a; Abrahão et al., 2018). Since their first recorded successful isolation from coastal waters (Garza and Suttle, 1995), Nucleocytoviruses (hereafter referred to as NCVs) have been discovered in oceans, freshwaters, soils, permafrost, and wastewater treatment plants (Legendre et al., 2014b, 2015; Zhang et al., 2015; Schulz et al., 2018, 2022; Minch and Moniruzzaman, 2025), and classified as *Megaviricetes, Pokkesviricetes* and *Mriyaviricetes*. Not all NCVs display “giant” features, as neither virion nor genome gigantism is universal throughout the phylum. However, NCV genomes have been shown to be highly mosaic, with DNA fragments from different sources (eukaryotic, bacterial, and viral) incorporated into the genome (Filée et al., 2007; Moreira and Brochier-Armanet, 2008; Filée, 2013). NCVs infect eukaryotes ranging from metazoans to protists. The majority of cultivated NCVs have been isolated through co-cultivation with free-living amoebae, particularly *Acanthamoeba* and *Vermamoeba* species, which serve as model hosts for NCV biology (Colson et al., 2017) (Azza et al., 2009; Legendre et al., 2014b). At the same time, many unicellular eukaryotes, especially green algae and amoebae, host bacterial (endo-)symbionts, creating complex but intriguing tripartite eukaryote-virus-bacteria interactions (Arthofer et al., 2022).

Adding yet another layer of complexity, NCVs are targeted by hyperparasitic entities. Viruses of the phylum *Preplasmiviricota* are small genetic elements (10 - 35 kbp) that co-infect protists together with NCVs (Roux et al., 2023; Koonin et al., 2024) and rely on these giant counterparts for replication (La Scola et al., 2008; Fischer and Suttle, 2011; Stough et al., 2019; Roitman et al., 2023). In contrast to virophages (class *Virophaviricetes*), polinton-like viruses (PLVs, *Aquintoviricetes*) have not been shown to form viral particles. Nonetheless, their coding capacity suggests their ability to spread in a horizontal manner (Yutin et al., 2015; Bellas and Sommaruga, 2021) and both are regularly found to integrate into eukaryotic or NCV genomes (Blanc et al., 2015; Fischer and Hackl, 2016; Schulz et al., 2018; Bellas et al., 2023).

The convergence of NCVs, virophages, PLVs, and sympatric bacteria within the host cell creates an exceptional hub of potential interactions among these entities. In any given system in which four players interact, six *direct* interactions are already possible. Many of these interactions have been described for subsets of eukaryote-virus-bacteria-virophage systems, yet a holistic understanding remains unexplored. In addition to the extensive gene sharing, recombination, and integration events observed in these systems, diverse defense and counter-defense mechanisms have been described across multiple interaction levels. A prominent example of this interaction constitutes the virophage *Mavirus*, which readily integrates into the host *Cafeteria roenbergensis* and acts as an inducible defense system by suppressing subsequent infections of the NCV Cafeteria roenbergensis virus (CroV) (Fischer and Hackl, 2016).

Capturing these interactions in isolated systems is difficult. To remain experimentally tractable, such systems are required to be stable (which they might inherently not be) in culture; however, this stabilization inevitably leads to the loss of the complexity that naturally arises in environments shaped by fluctuating abiotic and biotic conditions. Therefore, studying these interactions through culture-independent methods has the potential to better capture the diversity and complexity of microbial communities.

In this study, we aimed to overcome these limitations by investigating (eukaryote)-virus-bacteria-virophage systems at the community scale directly in their natural environment. Using deeply sequenced metagenomic data from wastewater treatment plants, we reconstructed novel and highly curated genomes of NCVs, virophages, and PLVs. Leveraging the untargeted deep-sequencing nature of the datasets, we identified viral, bacterial, and hyperparasitic interaction partners. Based on high-quality viral genomes and microbial contigs, we demonstrate genome-level interactions between these entities by tracing integration and gene-sharing events and leveraging co-occurrence data to establish a multilayered network approach to disentangle the resulting complex interaction landscape.

## Methods

### Sampling, sequencing and initial genome assembly

This study is based on 23 metagenomes obtained from sequencing activated sludge samples collected from wastewater treatment plants in Denmark. Details on sampling, sequencing, and assembly are provided in Singleton et al. (Singleton et al., 2021). Raw short reads, long reads, and assembled contigs are publicly available under BioProject PRJNA629478. For this study, paired-end Illumina HiSeq X Ten and Nanopore PromethION reads were retrieved, along with the corresponding hybrid assemblies generated by Singleton et al., using CANU v1.8. Nanopore reads were additionally trimmed with Porechop v0.2.4 (https://github.com/rrwick/Porechop).

### Initial detection of potential Nucleocytoviruses

For an initial scan of the retrieved contigs, open reading frames were predicted using prodigal v2.6.3 (Hyatt et al., 2010) in *-meta* mode. Proteins were then scanned using hmmsearch, part of the hmmer3 suite v3.4 (Eddy, 2011), with a set of nine hmm models of NCV core genes (MCP, A32, PolB, Topo2, TF2B, VLT3, RNAPS, RNAPL and SF2) with custom hmm cutoffs, as described elsewhere (Moniruzzaman et al., 2020). Per gene cutoffs are listed in Supplementary Data 1, Tab. S3. Contigs with at least one hit were considered for downstream analysis: rRNA genes were predicted using rnammer v1.2 (Lagesen et al., 2007) with *-S bac -m lsu,ssu,tsu* parameters. Kraken2 v2.1.3 (Wood et al., 2019) was used to determine the percentage of proteins assigned to Bacteria, Archaea, and Eukaryotes. Additionally, a similar percentage was also calculated manually by searching the proteins against *nr* (downloaded 21.08.2024), using diamond blastp v2.1.11 (Buchfink et al., 2021) with *--max-target-seqs 1 --query-cover 50 --id 70 --fast* parameters. Lastly, a ViralRecall (Aylward and Moniruzzaman, 2021) score was computed using the parameters *-c -s 0 -w 15 -m 20 -e 1e-5 -g 3 -f*.

Only contigs with a Kraken score <= 60%, a Diamond blastp score <= 60% (less than 60% of the proteins were assigned to be of eukaryotic, archaeal or bacterial origin), a ViralRecall score >= 0, no rRNA genes detected and a sequence length above 30 kbp were selected for further downstream work.

### Reassembly and polishing

To ensure the best possible assembly of NCV genomes, all potential NCV genomes were reassembled in a multistep, iterative manner: First, Nanopore PromethION long reads were mapped to potential NCV contigs (see above) for each sample to avoid cross-contamination. Using minimap v2.30 (Li, 2018) in *Nanopore* mode *-ax map-ont*. A set of artificial reads for each contig was created as a backbone for re-assembly. For this, the contig was split into 20 kbp-long sequences, overlapping by 10 kbp. Then, reads mapped to this specific contig, as well as unmapped reads, were used as input for reassembly with flye v2.9.5 (Kolmogorov et al., 2019) in *–scaffold –meta* mode. Afterwards, circularity and length of each NCV genome were assessed. If a genome did not increase in size, compared to the input contig, or was circularized in the process, no further re-assembly steps were taken. However, if the genome grew by more than ∼1000 bp, reads were re-mapped to the new genome, and unmapped reads mapping to this contig, as well as artificial reads, were assembled again as described before. This was repeated up to five times until all genomes were closed or did not increase in size again. For all final genomes, the assembly graph was visualized using bandage v0.8.1 (Wick et al., 2015).

NCV genomes were refined by polishing with Illumina HiSeqXTen PE short reads. These were mapped to the unpolished contigs using bwa v0.7.18-r1243-dirty (Li and Durbin, 2009), filtered and polished with polypolish v0.6.0 (Wick and Holt, 2022) with default parameters.

### Gene prediction and completeness assessment of nucleocytoviruses genomes

Genes of the refined NCV genomes were predicted using prodigal v2.6.3 in metagenomic mode *(-p meta)*. NCV genomes were classified as being either “complete” or “draft”. A genome was considered “complete” if it was found to be circular, or b) had terminal inverted repeats and 3 or more marker genes were detected, or c) was over 250 kbp long with at least 5 marker genes detected, or d) it was one of the seven identified *Yaraviridae*.

*Nucleocytoviruses dereplication, terminal repeats, taxonomic assignment, and annotation*.

Genomes were dereplicated using dRep v3.5.0 (Olm et al., 2017) with 95% ANI cutoff and a minimum length of 10 kbp (Supplementary Figures 1, Fig. 1). Secondary structure of the ends of linear genomes was calculated using the DNAfold webservice https://www.unafold.org/mfold/applications/dna-folding-form.php, (SantaLucia, 1998). GVClass v1.0.8 (Pitot et al., 2024) and TIGTOG (Ha and Aylward, 2024) were used for initial taxonomic assignment of the NCV genomes. Additionally, 10 kbp segments were manually searched against the nr-database using the tblastx web interface. Furthermore, the position of each genome in each tree based on seven core genes (MCP, A32, PolB, Topo2, TF2B, VLT3 and SF2) was taken into account, as was the collapsed phylogenetic tree described below. All these lines of evidence were taken together to present a final taxonomic assignment. An overview of the taxonomic assignment of each genome is given in Supplementary Data 1, Tab. S1.

**Figure 1:**
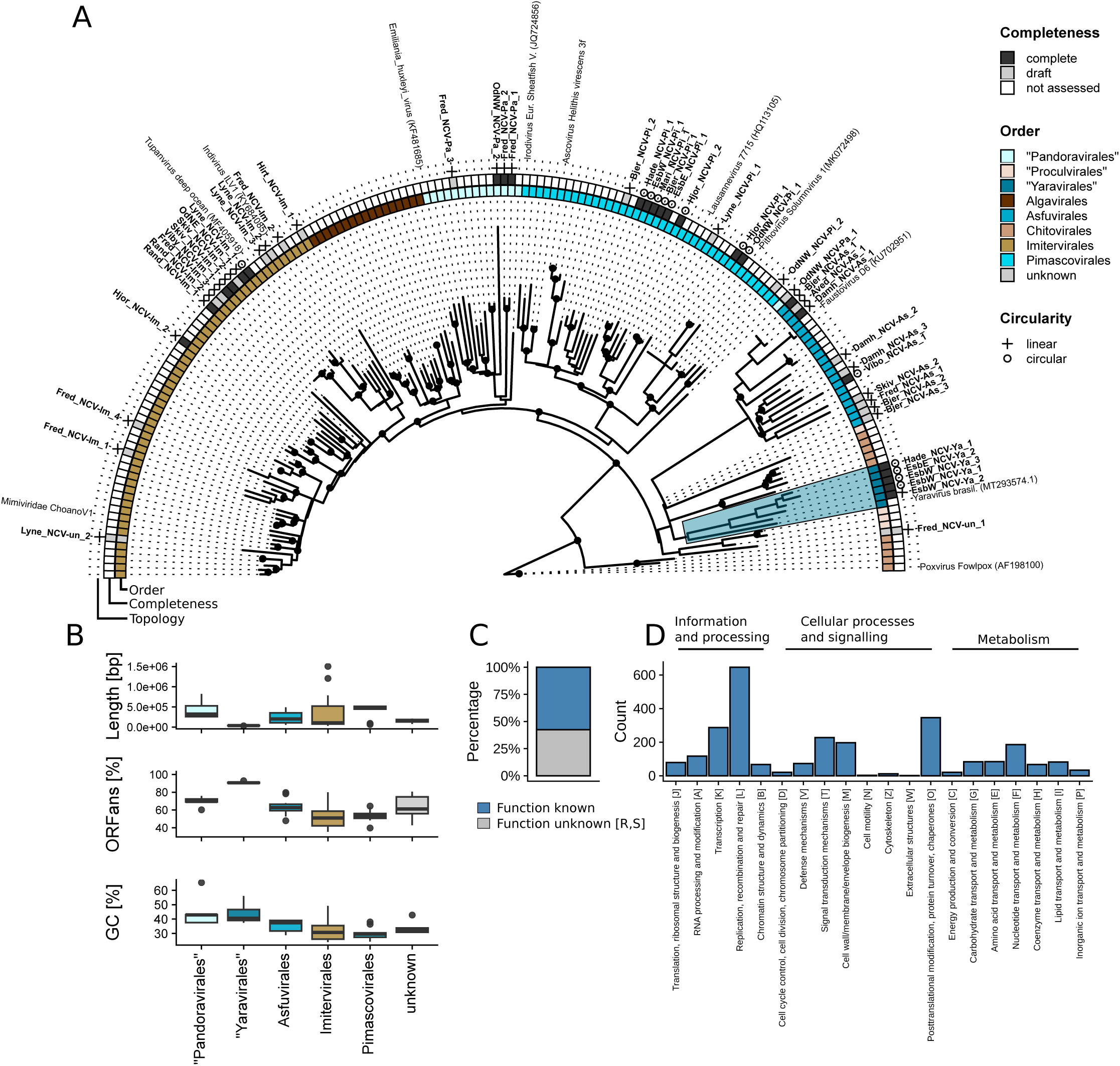
Phylogenetic tree of novel NCVs and references. Panel A depicts the collapsed phylogenetic tree of identified NCVs and references, with Poxvirus fowlpox (AF198100) as the outgroup. For that, a maximum-likelihood tree was calculated for the nine core genes of NCVs (MCP, SFII, RNAPS, RNAPL, PolB, TF2B, Topo2, A32 and VLTF3). Details are given in the methods. Black circles on the branches indicate branch support >75%. Highlighted in blue is the “Yaravirales” clade, with five genomes from this study and one reference genome included. The innermost ring indicates the order, either assigned to sequences recovered in this study based on the individual core genes or the assigned order of references. The next ring indicates the assessed completeness of the recovered NCVs, and the outermost ring depicts their circularity. Panel B shows the length and GC content of the recovered NCVs by assigned order. Panel C shows the percentage of NCV genes for which a COG category other than S (function unknown) and R (general function only) could be assigned. Panel D then gives the number of genes assigned to each COG category, grouped by general function.

Terminal repeats were predicted using TRFfinder (https://tandem.bu.edu/trf/trf.html) (Benson, 1999) and a custom approach, using blastn, part of the ncbiblastplus 2.16.0 suite (Camacho et al., 2009), with parameter -word_size 7 and searching 500 bp ends of each genome against each other. tRNA genes and introns were identified with ARAGORN v1.2.41 (Laslett and Canback, 2004). The funciotnal annotation of the predicted ORFs of the recovered *Nucleocytoviricota* was done with the PFAM (Mistry et al., 2021), GVOG (Aylward et al., 2021), and VOG databases (Grazziotin et al., 2017), and using the AnnoMazing pipeline (https://github.com/BenMinch/AnnoMazing). Defense systems were detected using padloc v2.0.0, (Payne et al., 2021) with default parameters. For proteins associated with PD-T4-6, namely Serine/Threonine or Tyrosine protein kinase, structures were predicted using the AlphaFold3 webserver (https://alphafoldserver.com/, accessed May 2026; Abramson et al., 2024) and compared to structures of PD-T4-7 (WP_036366335) and Poxvirus Kinase (AF-A0A6C0JZI9-F1) using ChimeraX v1.9 (Pettersen et al., 2021). COG categories were determined using eggNOG-mapper v2.1.13-f071154 with eggNOG-DB v5.0.2 (Huerta-Cepas et al., 2019; Buchfink et al., 2021; Cantalapiedra et al., 2021), with default parameters.

### Nucleocytoviricota phylogenetic tree construction

The resulting 108 reference genomes and all genomes identified in this study, were then searched for marker genes as described above. For each NCV marker gene (MCP, A32, PolB, Topo2, TF2B, VLT3, RNAPS, RNAPL and SF2), a phylogenetic tree was calculated using NuPhylo (https://github.com/BenMinch/NuPhylo, accessed August 2025), including a set of reference NCV genomes. Individual trees were visualized using iTol v6 (Letunic and Bork, 2024). Additionally, the individual trees were collapsed into a single tree using astral, part of the aster package v1.23.4.6 (Zhang et al., 2025) with default parameters. The resulting tree was visualized in R, using ggtree and ggtreeExtra (R Core Team, 2024).

### Mriyaviricetes major capsid structural prediction and phylogenetic tree construction

Protein structures of major capsid proteins of *Mriyaviricetes* identified in this study, *Yaravirus brasiliense* (NC_076895.1) and *Pleurochrysis sp. endemic virus 2* (AUD57312.1) were predicted using the AlphaFold3 webserver (https://alphafoldserver.com/, accessed March 2026). Structures were aligned, and alignment scores were calculated in ChimeraX v1.9. The identified MCP proteins were aligned with reference *Mriyaviricetes* MCP proteins from elsewhere (Yutin et al., 2024), using mafft v7.526. The resulting alignment was trimmed using trimAl v1.5.rev1 (Capella-Gutiérrez et al., 2009) and visualized in iTol v6.

### Virophage and polinton-like virus detection, classification, phylogenetic tree construction, genome maps

Open reading frames were predicted for all previously assembled contigs (Singleton 2021), using prodigal v2.6.3 in metagenomic mode *(-p meta)*. Virophages were detected by searching protein sequences of the assembled contigs with hmm models of four core genes (ATPase, MCP, Penton / mCP, PRO), as described elsewhere (Roux et al., 2023), using hmmsearch 3.4 with custom cutoffs *-T 50 –incE 0.01,* part of the hmmer suite (Eddy 2011). Contigs with at least three of the four virophage core genes were selected for downstream work. Similarly, polinton-like viruses were detected by searching assembled contigs with the hmm model of established MCP alignments of PLVs (Bellas et al., 2023), using hmmsearch (v3.4) with custom cutoffs *-T 50 –incE 0.01*. Contigs with a hit above these thresholds were retrieved and considered PLV-like sequences.

Reference proteins were retrieved from elsewhere (Roux et al., 2023), however only genomes marked as “complete”, coming from NCBI, IMG_VR v3 and Hackl et al. (Hackl et al., 2021) were included. Proteins were aligned using mafft in *-auto* mode v7.526, (Rozewicki et al., 2019), manually curated, and a phylogenetic tree was constructed using iqtree2 (Minh et al., 2020) with the *-B 1000* flag. The resulting tree was visualized in R, using ggtree and ggtreeExtra.

### Annotation of virophage and polinton-like viruses, terminal repeats

Terminal repeats of both virophages and PLVs were detected by blasting 1000 bp of each end of the sequence against each other, using blastn, part of the ncbiblastplus suite v2.16.0, with the following parameters: *-word_size 7, -evalue 1e-3, -strand minus*. Only hits longer than 100 bp were considered terminal repeats. Annotation was done using InterProScan5 v5.74-105.0 (Jones et al., 2014) with default parameters and an e-value cutoff of 10e-5.

### Selection and taxonomic classification of potential interaction partners (MC contigs)

In order to include potential microbial interaction partners in the downstream network analysis, contigs from the original assembly, longer than 100,000 bp were selected. As described above, NCV genomes were iteratively re-assembled, hence it was necessary to remove any potential microbial interaction partners whose sequence was now part of the re-assembled NCV genomes. Therefore, all contigs were compared to the final NCV genomes using blastn, part of the ncbiblastplus v2.16.0+ suite with the following parameters: *-task megablast -perc_identity 95 -qcov_hsp_perc 80.* A custom script was developed to taxonomically classify the selected microbial contigs (scripts are provided in the zenodo repository, *scripts/diamond_assign_classification.sh*). Protein-coding sequences were predicted from contigs using prodigal v2.6.3 in metagenomic mode *(-p meta)*. Predicted proteins were searched against a locally formatted NCBI NR database using DIAMOND v2.1.8 BLASTP (*--max-target-seqs 1, --evalue 1e-5, --faster, --threads 16*). For each contig, taxonomic assignment was inferred by extracting the organism name of the top-scoring hit per protein and identifying the majority taxon, along with the percentage of proteins supporting it. This resulted in 51,703 potential microbial interaction partners. Lastly, the contigs were clustered using a genome dissimilarity threshold of 0.05 (>95% similarity), calculated with mash v2.3 (Ondov et al., 2016) using the *-s 10000* parameter. This resulted in 23,519 microbial clusters.

### Network analysis: node and edge definitions

To construct the multilayered network, the following contigs were considered possible nodes: 14 polinton-like viruses, 15 virophages, 61 NCVs, 51,703 microbial contigs (>100,000 kbp). In order to reduce complexity while preserving the biological signal, microbial contigs were subsequently collapsed into microbial clusters (as described above). Edge lists were therefore updated to reflect cluster-level connections: edges between individual microbial contigs within the same cluster were removed, and edges from multiple microbial contigs to the same target were deduplicated.

In total, three interaction layers were constructed (see below): Gene sharing, co-occurrence and integration. Each layer resulted in (un)directed, non-weighted edges. Details on how the interaction layers were constructed are given in the respective paragraphs.

### Network analysis: gene sharing layer

For all contigs of interest (NCVs, virophages, PLVs and microbial contigs), proteins were predicted using prodigal in *-meta* mode. Then, an all-against-all blastp search was performed, using diamond with the following parameters: *–min-score 50, –sensitive.* Results were filtered and only hits above 80% AAI and 80% query coverage (*qcovhsp*) were kept. Contigs were connected if >5% of the total nucleotides were shared. The weight of an edge in this layer was set to 1.

### Network analysis: co-occurrence layer

This layer was technically split into four distinct layers: Positive and negative co-occurrence based on short-read mapping, and positive and negative co-occurrence based on long-read mapping. Short reads were first dereplicated using cd-hit-dup, part of the CD-HIT suite v4.8.1 (Huang et al., 2010), with *-e 0* and otherwise default parameters.

Short reads were mapped to all (microbial, NCV, PLV and virophage contigs with bowtie2 v2.5.4, (Langmead et al., 2009), using the following parameters: *--sensitive-local --no-mixed --no-unal -I 0 -X 1000.* Samtools v1.22 (Li et al., 2009; Danecek et al., 2021) was used to filter and retain only mapped reads and calculate per-contig coverage values.

Long reads were mapped using minimap2 v2.30-r1287, in *-ax map-ont* mapping mode. Samtools was used to retain mapped-only reads and calculate per-contig coverage values. Microbial contigs had to be present in at least 50% (12/23) of the samples to be considered. Only edges detected in both long and short-read co-occurrence correlations were considered. Correlation was calculated using Spearman’s rank correlation (ρ) on centered log-ratio (CLR) transformed coverage values. Co-occurrence edges were weighted according to correlation strength: 5 for very strong (|ρ| > 0.80), 3 for strong (|ρ| > 0.70), and 1 for moderate (|ρ| > 0.60) correlations.

### Network analysis: integration layer

In order to identify possible integration events, the following steps were performed: First, long reads were mapped to NCV, virophage and PLV sequences in *-ax map-ont* mode of minimap2. Samtools was used to retain only successfully mapped reads in the resulting bam/sam files. Then, reads were extracted from the mapping files, using a custom Python script (*scripts/pysam_filter_and_extract_overhangs.py*). Here, each read was first subjected to stringent length filtering, retaining only reads longer than 1000 bp, with at least 500 bp mapped to the reference contig. Then, CIGAR strings of each mapped read were analyzed, identifying soft-clipped (overhanging) reads, where the overhanging segment was longer than 100 bp. Reads overhanging in such a manner were then classified as “boundary overhangs”, if the reads alignment start or end position was within 50 bp of the start or end of the reference contig, or as “internal/middle overhang” if the alignment was not within 50 bp of the ends of the reference. The overhanging segments of the reads were then saved and blasted against all (microbial, NCVs, PLV and virophage) contigs, using blastn, part of the ncbiblastplus suite, with default parameters. Only matches with> 80% nucleotide identity and >80% coverage of the read segment were retained, thereby establishing “boundary” or “middle” integration interaction events between contigs. The weight of an edge in this layer was set to 100.

### Network analysis: measures of centrality, detection of clusters, visualization

The following R-libraries were used for network visualization and analysis: ggiraph, tidygraph, ggraph, igraphdat, igraph. The undirected network graph was constructed using the *graph_from_frame()* function from the igraph library, with *directed = FALSE*. Louvain groups, used for single subcluster visualization, were calculated using the *group_graph()* (tidygraph) function with resolution set to 10. *Degree* and *betweenness* for the network were calculated using the corresponding functions in the igraph library. *Betweenness* was calculated with the parameters *directed = FALSE* and *normalized = FALSE*. In visualizations, biologically directed edges (integration) are represented with arrows to indicate the biological direction of the interaction, while undirected edges (gene sharing, co-occurrence, integration) are shown as lines. Network clusters were visualized using the ggiraph library, and interactive figures were created with the *girafe()* function of ggiraph. Scale-free properties of the network were assessed by calculating degree distribution using the poweRlaw library (Gillespie, 2015). Minimum degree threshold (xmin) and scaling exponent (alpha) were estimated using maximum likelihood, and the two models were compared using a likelihood ratio test to determine which model better fit the observed degree distribution.

## Results

### Recovery of novel Nucleocytoviricota

We devised a multi-step, iterative reassembly approach to recover complete or near-complete NCVs from 23 wastewater treatment plants in Denmark, which had been previously sequenced and assembled elsewhere (Singleton et al., 2021). Briefly, assembled contigs were screened using HMM profiles of established NCV marker genes (NCLDV major capsid protein - MCP; Packaging ATPase - A32, DNA polymerase family B - PolB, DNA topoisomerase II - Topo2, Transcription initiation factor IIB - TF2B, Poxvirus Late Transcription Factor - VLTF3, DNA-directed RNA polymerase beta subunit - RNAPS, DNA-directed RNA polymerase alpha subunit - RNAPL and DEAD/SNF2-like helicase - SF2; Aylward et al., 2021). Potential NCV contigs were subsequently filtered by contig size and the number of marker-gene hits and iteratively reassembled until the resulting contig no longer grew or was identified as circular. Finally, genome completeness was carefully assessed using stringent criteria (number of marker-gene hits, genome size and topology, terminal inverted repeats; see Methods).

This iterative assembly and curation yielded 61 high-quality NCV genomes: 28 complete and 33 draft genomes. All genomes were double-stranded DNA, with inferred linear (n = 45) or circular (n = 16) topology. For all nine marker genes of the recovered NCV genomes and 108 reference genomes, individual alignments and phylogenetic trees were generated (Supplementary Figures 1, Fig. 2-10) and subsequently collapsed into a single consensus tree (Fig. 1a). Of the 28 complete genomes, four were classified as belonging to the order of *Asfuvirales*, six as *Imitervirales*, three as *Pandoravirales*, eight as *Pimascovirales* and seven as *Yaravirales*. Beyond phylogenetic placement, the genomes (complete and draft) displayed distinct features: genome sizes ranged from 30 kbp (*Yaravirales* EsbE_NCV-Ya_1) to 1.5 mbp (*Imitervirales* OdnW_NCV-Im_1), with 25 to 1394 ORFs, and GC content ranging from 24% to 65% (Fig. 1b). The predicted coding capacity varied substantially across orders, with ORFans accounting for ∼40% (*Imitervirales*) to ∼90% (*Yaravirales*) of all ORFs. Assigning NCV genes to COG categories revealed that nearly all categories are present, with only a few cell function-related ones occurring at lower frequency (Fig 1c): Cell motility (N), cytoskeleton (Z), energy production and conversion (C), and cell cycle control, cell division, chromosome partitioning (D) (Fig 1d). On the other hand, most genes were assigned to categories associated with replication (replication, recombination and repair [L], transcription [K] and posttranslational modification, protein turnover, chaperons [O]). The number of marker genes detected varied substantially, and only four genomes contained more than eight of the nine marker genes (Supplementary Data 1, Tab. S1), broadly consistent with phylogenetic placement. Several genomes encoded putative tRNAs (Tab. 1). Most strikingly, Lyne_NCV-un_3, an unclassified draft genome, encoded 27 (21 distinct) tRNA genes. The unclassified genome Lyne_NCV-un_3 displays an interesting phylogenetic placement, forming an outgroup to a clade containing the putative new *Nucleocytoviricota* class “*Proculviricetes*” (Fig. 1a), a lineage previously known only from MAGs detected in polar marine environments of the Arctic and Southern Oceans, suggesting potential biogeographic diversification within this emerging taxonomic group (Delmont et al., 2022). Nine NCV genomes encoded a Thr/Ser kinase. Similar kinases are involved in viral protein phosphorylation in e.g., *Poxviridae* (Punjabi and Traktman, 2005), however the detected proteins showed more structural similarity to the phage defense system PD-T4-6 kinase, originally described in bacteria (Supplementary Data 1, Tab. S2; Vassallo et al., 2022).

**Figure 2:**
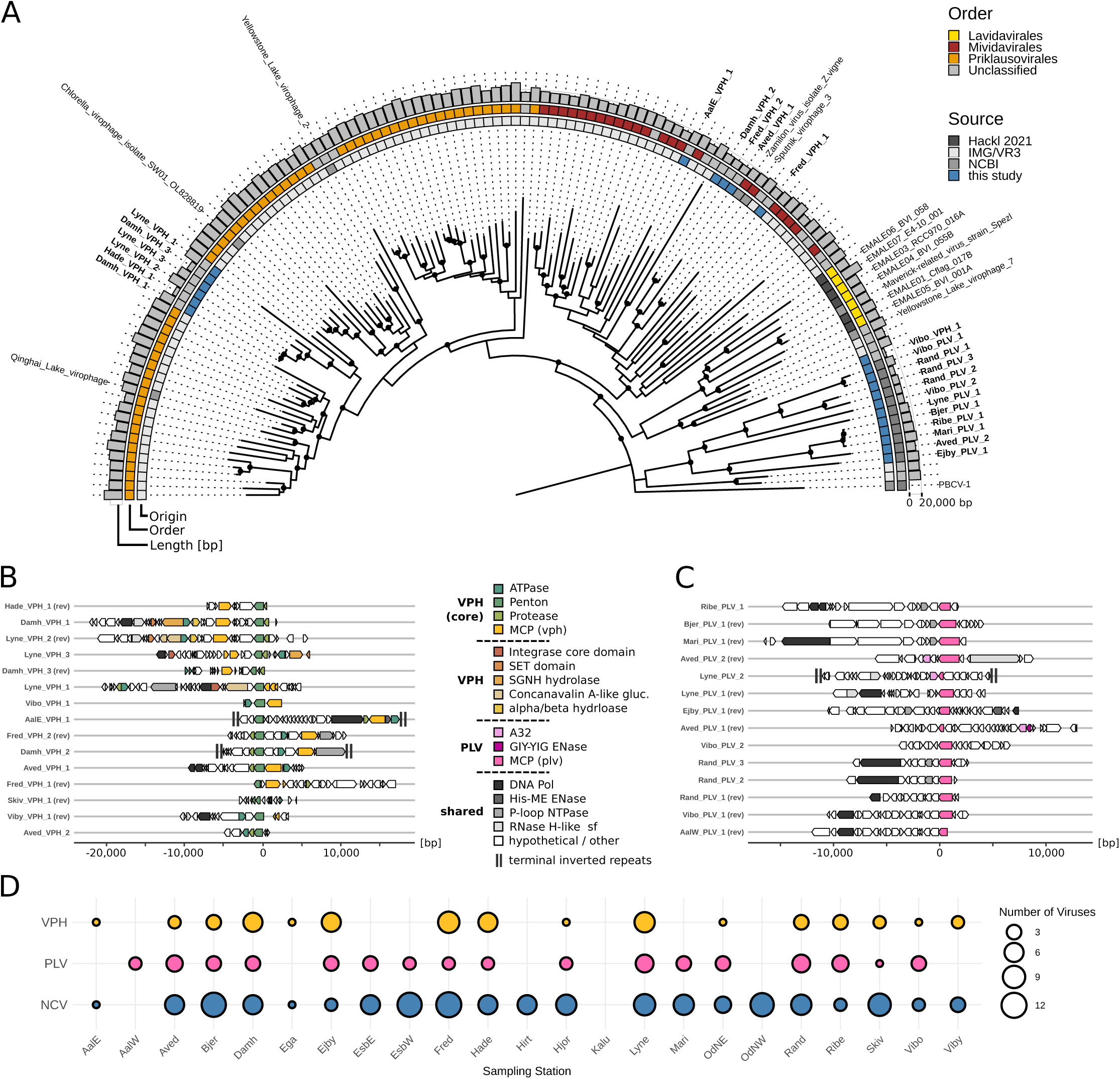
Phylogenetic tree and genome maps of identified virophages, polinton-like viruses and references. Panel A depicts the MCP phylogeny of virophages and polinton-like viruses from four sources (Hackl 2021, IMG/VR3, NCBI and this study). Black dots indicate branch support >90%. The inner ring indicates the source, the middle ring indicates the assigned order of reference genomes, and the outermost ring indicates the length of each genome. Panel B shows the genome maps for 15 recovered virophages, and panel C the genome map of 14 polinton-like viruses. Panel D indicates the number of NCV/ virophage/PLV contigs recovered from each sampling station. MCP_1 - virophage major capsid protein; ATPase_1 - virophage ATPase; DNA Pol - DNA polymerase; Penton_1 - virophage minor capsid protein (Penton protein); PRO_1 - virophage protease; A32 - Poxvirus A32 protein; GIY-YIG ENase - GIY-YIG Endonuclease; P-loop NTPase - P-loop containing nucleoside triphosphate hydrolases; His-Me ENase - His-Me finger endonucleases; RNaseH-like sf - Ribonuclease H-like; hypothetical - hypothetical protein.

**Table 1:**
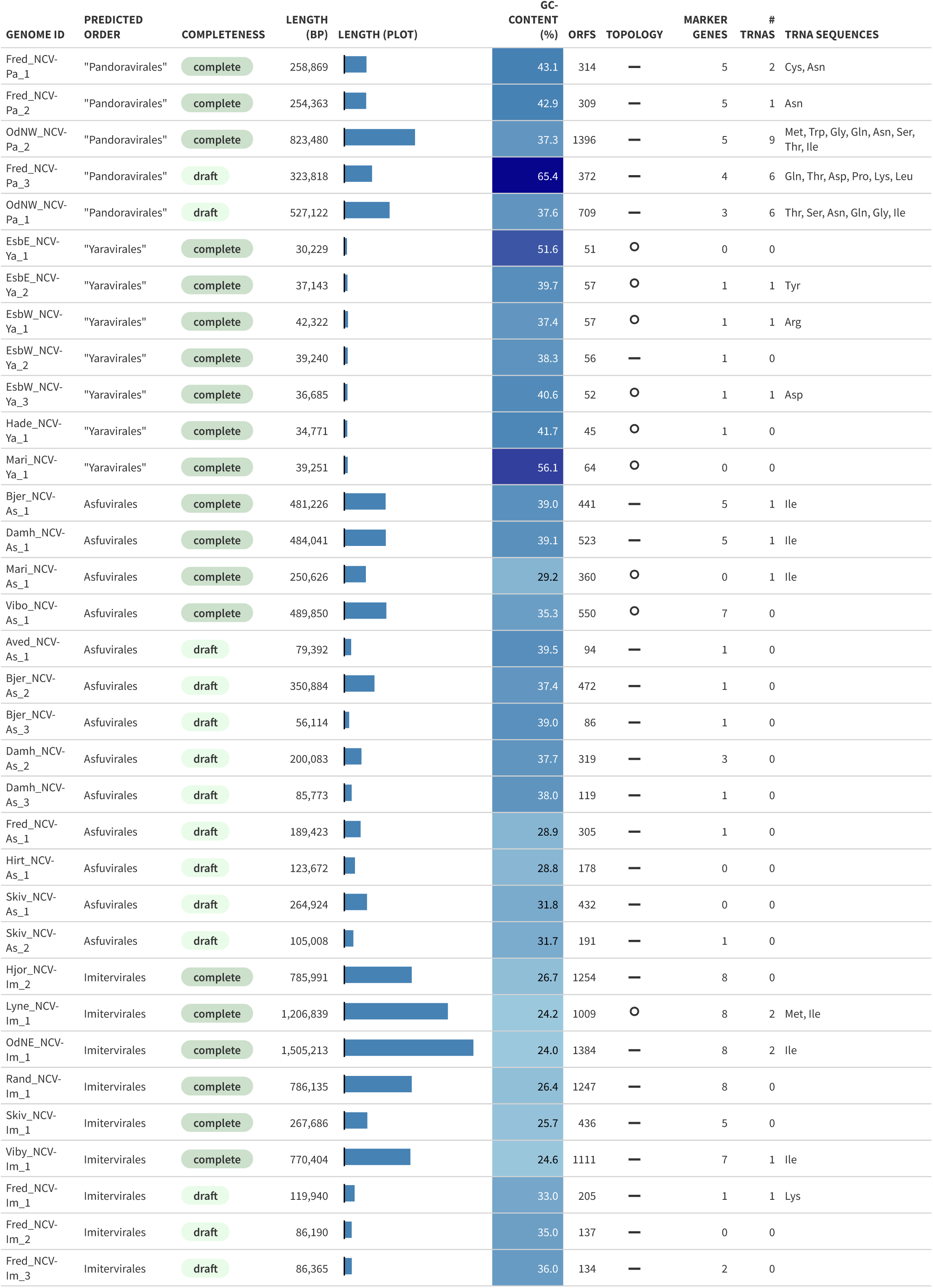
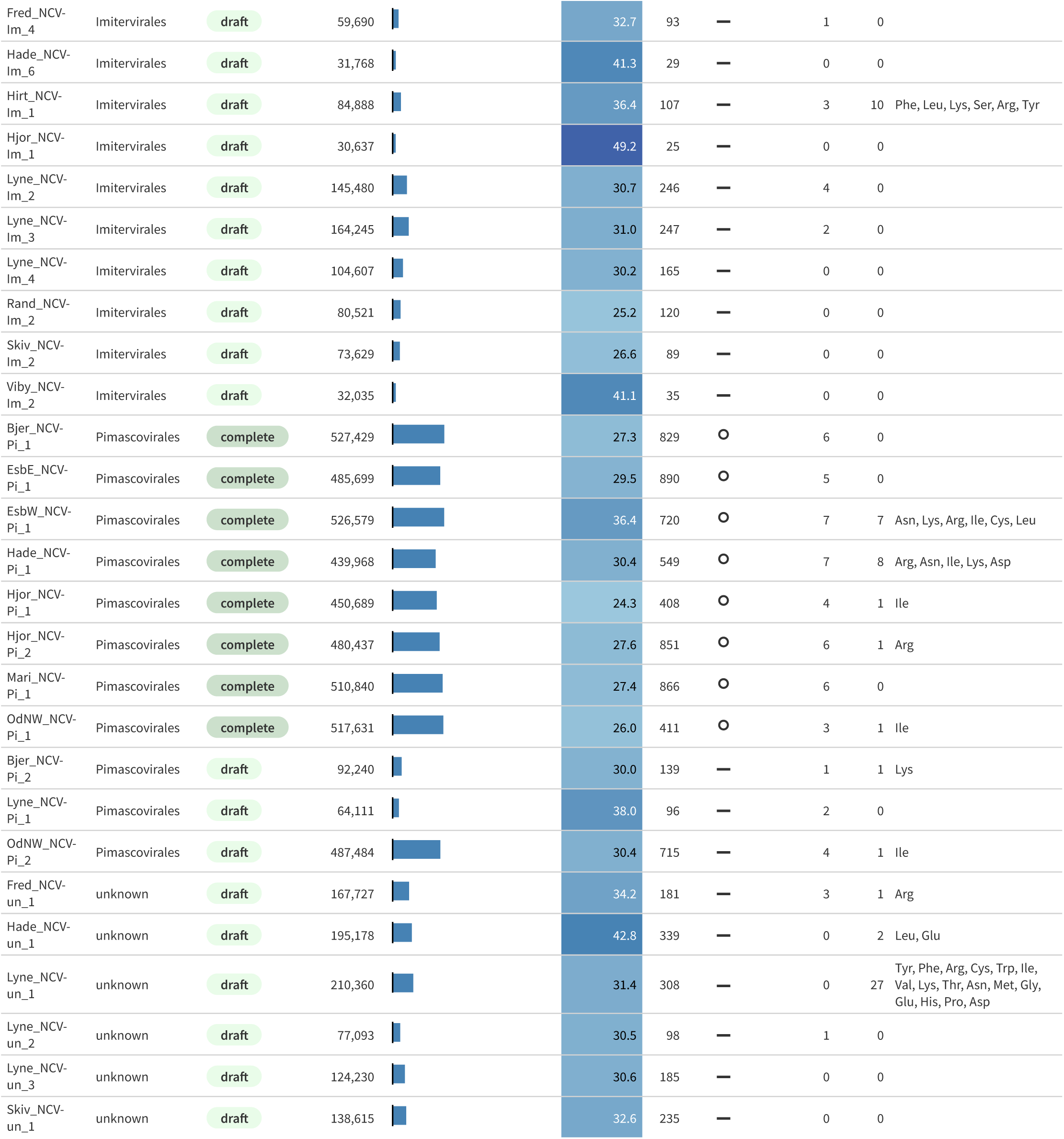
Overview of key genomic features of identified Nucleocytoviricota.

| Genome ID | Predicted Order | Completeness | Length (bp) | Length (Plot) | GC-Content (%) | ORFs | Topology | Marker Genes | # tRNAs | tRNA Sequences |
| --- | --- | --- | --- | --- | --- | --- | --- | --- | --- | --- |
| Fred_NCV-Pa_1 | "Pandoravirales" | complete | 258,869 |  | 43.1 | 314 | — | 5 | 2 | Cys, Asn |
| Fred_NCV-Pa_2 | "Pandoravirales" | complete | 254,363 |  | 42.9 | 309 | — | 5 | 1 | Asn |
| OdNW_NCV-Pa_2 | "Pandoravirales" | complete | 823,480 |  | 37.3 | 1396 | — | 5 | 9 | Met, Trp, Gly, Gln, Asn, Ser, Thr, Ile |
| Fred_NCV-Pa_3 | "Pandoravirales" | draft | 323,818 |  | 65.4 | 372 | — | 4 | 6 | Gln, Thr, Asp, Pro, Lys, Leu |
| OdNW_NCV-Pa_1 | "Pandoravirales" | draft | 527,122 |  | 37.6 | 709 | — | 3 | 6 | Thr, Ser, Asn, Gln, Gly, Ile |
| EsbE_NCV-Ya_1 | "Yaravirales" | complete | 30,229 |  | 51.6 | 51 | ○ | 0 | 0 |  |
| EsbE_NCV-Ya_2 | "Yaravirales" | complete | 37,143 |  | 39.7 | 57 | ○ | 1 | 1 | Tyr |
| EsbW_NCV-Ya_1 | "Yaravirales" | complete | 42,322 |  | 37.4 | 57 | ○ | 1 | 1 | Arg |
| EsbW_NCV-Ya_2 | "Yaravirales" | complete | 39,240 |  | 38.3 | 56 | — | 1 | 0 |  |
| EsbW_NCV-Ya_3 | "Yaravirales" | complete | 36,685 |  | 40.6 | 52 | ○ | 1 | 1 | Asp |
| Hade_NCV-Ya_1 | "Yaravirales" | complete | 34,771 |  | 41.7 | 45 | ○ | 1 | 0 |  |
| Mari_NCV-Ya_1 | "Yaravirales" | complete | 39,251 |  | 56.1 | 64 | ○ | 0 | 0 |  |
| Bjer_NCV-As_1 | Asfuvirales | complete | 481,226 |  | 39.0 | 441 | — | 5 | 1 | Ile |
| Damh_NCV-As_1 | Asfuvirales | complete | 484,041 |  | 39.1 | 523 | — | 5 | 1 | Ile |
| Mari_NCV-As_1 | Asfuvirales | complete | 250,626 |  | 29.2 | 360 | ○ | 0 | 1 | Ile |
| Vibo_NCV-As_1 | Asfuvirales | complete | 489,850 |  | 35.3 | 550 | ○ | 7 | 0 |  |
| Aved_NCV-As_1 | Asfuvirales | draft | 79,392 |  | 39.5 | 94 | — | 1 | 0 |  |
| Bjer_NCV-As_2 | Asfuvirales | draft | 350,884 |  | 37.4 | 472 | — | 1 | 0 |  |
| Bjer_NCV-As_3 | Asfuvirales | draft | 56,114 |  | 39.0 | 86 | — | 1 | 0 |  |
| Damh_NCV-As_2 | Asfuvirales | draft | 200,083 |  | 37.7 | 319 | — | 3 | 0 |  |
| Damh_NCV-As_3 | Asfuvirales | draft | 85,773 |  | 38.0 | 119 | — | 1 | 0 |  |
| Fred_NCV-As_1 | Asfuvirales | draft | 189,423 |  | 28.9 | 305 | — | 1 | 0 |  |
| Hirt_NCV-As_1 | Asfuvirales | draft | 123,672 |  | 28.8 | 178 | — | 0 | 0 |  |
| Skiv_NCV-As_1 | Asfuvirales | draft | 264,924 |  | 31.8 | 432 | — | 0 | 0 |  |
| Skiv_NCV-As_2 | Asfuvirales | draft | 105,008 |  | 31.7 | 191 | — | 1 | 0 |  |
| Hjor_NCV-Im_2 | Imitervirales | complete | 785,991 |  | 26.7 | 1254 | — | 8 | 0 |  |
| Lyne_NCV-Im_1 | Imitervirales | complete | 1,206,839 |  | 24.2 | 1009 | ○ | 8 | 2 | Met, Ile |
| OdNE_NCV-Im_1 | Imitervirales | complete | 1,505,213 |  | 24.0 | 1384 | — | 8 | 2 | Ile |
| Rand_NCV-Im_1 | Imitervirales | complete | 786,135 |  | 26.4 | 1247 | — | 8 | 0 |  |
| Skiv_NCV-Im_1 | Imitervirales | complete | 267,686 |  | 25.7 | 436 | — | 5 | 0 |  |
| Viby_NCV-Im_1 | Imitervirales | complete | 770,404 |  | 24.6 | 1111 | — | 7 | 1 | Ile |
| Fred_NCV-Im_1 | Imitervirales | draft | 119,940 |  | 33.0 | 205 | — | 1 | 1 | Lys |
| Fred_NCV-Im_2 | Imitervirales | draft | 86,190 |  | 35.0 | 137 | — | 0 | 0 |  |
| Fred_NCV-Im_3 | Imitervirales | draft | 86,365 |  | 36.0 | 134 | — | 2 | 0 |  |
| Fred_NCV-Im_4 | Imitervirales | draft | 59,690 |  | 32.7 | 93 | — | 1 | 0 |  |
| Hade_NCV-Im_6 | Imitervirales | draft | 31,768 |  | 41.3 | 29 | — | 0 | 0 |  |
| Hirt_NCV-Im_1 | Imitervirales | draft | 84,888 |  | 36.4 | 107 | — | 3 | 10 | Phe, Leu, Lys, Ser, Arg, Tyr |
| Hjor_NCV-Im_1 | Imitervirales | draft | 30,637 |  | 49.2 | 25 | — | 0 | 0 |  |
| Lyne_NCV-Im_2 | Imitervirales | draft | 145,480 |  | 30.7 | 246 | — | 4 | 0 |  |
| Lyne_NCV-Im_3 | Imitervirales | draft | 164,245 |  | 31.0 | 247 | — | 2 | 0 |  |
| Lyne_NCV-Im_4 | Imitervirales | draft | 104,607 |  | 30.2 | 165 | — | 0 | 0 |  |
| Rand_NCV-Im_2 | Imitervirales | draft | 80,521 |  | 25.2 | 120 | — | 0 | 0 |  |
| Skiv_NCV-Im_2 | Imitervirales | draft | 73,629 |  | 26.6 | 89 | — | 0 | 0 |  |
| Viby_NCV-Im_2 | Imitervirales | draft | 32,035 |  | 41.1 | 35 | — | 0 | 0 |  |
| Bjer_NCV-Pi_1 | Pimascovirales | complete | 527,429 |  | 27.3 | 829 | ○ | 6 | 0 |  |
| EsbE_NCV-Pi_1 | Pimascovirales | complete | 485,699 |  | 29.5 | 890 | ○ | 5 | 0 |  |
| EsbW_NCV-Pi_1 | Pimascovirales | complete | 526,579 |  | 36.4 | 720 | ○ | 7 | 7 | Asn, Lys, Arg, Ile, Cys, Leu |
| Hade_NCV-Pi_1 | Pimascovirales | complete | 439,968 |  | 30.4 | 549 | ○ | 7 | 8 | Arg, Asn, Ile, Lys, Asp |
| Hjor_NCV-Pi_1 | Pimascovirales | complete | 450,689 |  | 24.3 | 408 | ○ | 4 | 1 | Ile |
| Hjor_NCV-Pi_2 | Pimascovirales | complete | 480,437 |  | 27.6 | 851 | ○ | 6 | 1 | Arg |
| Mari_NCV-Pi_1 | Pimascovirales | complete | 510,840 |  | 27.4 | 866 | ○ | 6 | 0 |  |
| OdNW_NCV-Pi_1 | Pimascovirales | complete | 517,631 |  | 26.0 | 411 | ○ | 3 | 1 | Ile |
| Bjer_NCV-Pi_2 | Pimascovirales | draft | 92,240 |  | 30.0 | 139 | — | 1 | 1 | Lys |
| Lyne_NCV-Pi_1 | Pimascovirales | draft | 64,111 |  | 38.0 | 96 | — | 2 | 0 |  |
| OdNW_NCV-Pi_2 | Pimascovirales | draft | 487,484 |  | 30.4 | 715 | — | 4 | 1 | Ile |
| Fred_NCV-un_1 | unknown | draft | 167,727 |  | 34.2 | 181 | — | 3 | 1 | Arg |
| Hade_NCV-un_1 | unknown | draft | 195,178 |  | 42.8 | 339 | — | 0 | 2 | Leu, Glu |
| Lyne_NCV-un_1 | unknown | draft | 210,360 |  | 31.4 | 308 | — | 0 | 27 | Tyr, Phe, Arg, Cys, Trp, Ile, Val, Lys, Thr, Asn, Met, Gly, Glu, His, Pro, Asp |
| Lyne_NCV-un_2 | unknown | draft | 77,093 |  | 30.5 | 98 | — | 1 | 0 |  |
| Lyne_NCV-un_3 | unknown | draft | 124,230 |  | 30.6 | 185 | — | 0 | 0 |  |
| Skiv_NCV-un_1 | unknown | draft | 138,615 |  | 32.6 | 235 | — | 0 | 0 |  |

**Table 2:** Overview of interaction layers considered in network construction.

| Layer | Description | Weight | Directionality |
| --- | --- | --- | --- |
| Gene Sharing | all against all blastp, >80% pident, >80% qcov, at least 5% of genes shared | 1 | Undirected, all connections possible. |
| Co-occurrence | Split into positive/negative co-occurrence. Only the overlap between short- and long-read co-occurrence signals was considered. | 1,3,5 | Undirected, all connections possible. |
| Integration | Split into two separate layers: “boundary” and “middle” integration. | 100 (middle), 100 (boundary) | Directed, only (NCV, PLV, virophage) <-> ALL connections are possible. |

### Identification of seven novel Mriyaviricetes genomes

Among the identified NCV genomes, seven displayed an additional set of interesting characteristics: Only a single marker gene (A32-like ATPase) was detected above initial cutoffs (Supplementary Data 1, Tab. S3) in five of these genomes, while in two, we detected none. Furthermore, they showed a higher GC content (38% - 56%; p = 0.001) and shorter genome length (30 kbp - 42 kbp; p = 0.0001) compared to the other NCVs orders and a high percentage of genes with no homolog in any of the tested databases (∼90% ORFans, Fig. 1b), yet six of the seven genomes were inferred circular, indicating a complete genome. Initial phylogenetic analysis clustered them with *Yaravirus brasiliense* (class *Mriyaviricetes*, MT293574.1; Boratto et al., 2020), forming a deeply branching, distinct clade (Fig. 1a, highlighted in blue). Predicting and comparing protein structures to the major capsid proteins of the *Yaraviridae* representative *Yaravirus brasiliense* (NC_076895.1) and the “*Gamadviridae”* (*sensu* Yutin et al., 2024) representative *Pleurochrysis sp. endemic virus 2* (AUD57312.1) revealed candidate MCP proteins on all seven genomes (Supplementary Data 1, Tab. S4). After curating the candidate sequences, we constructed a maximum-likelihood phylogenetic tree of the recovered MCP candidates, together with representatives of the class *Mriyaviricetes*. This placed all seven genomes in a well-supported clade within *Yaraviridae*, leading us to classify these as novel viruses within this family (Supplementary Figures 1, Fig. 11).

### Identification of 15 novel virophages and 14 polinton-like viruses

We identified 15 virophages and 14 PLVs in the dataset (Fig. 2a), ranging from 4.3 to 24.8 kbp and 8.4 to 19 kbp, respectively (Fig. 2a, Supplementary Data 1, Tab. S5 and Tab. S6). The virophages detected in this study were all found in an excised form, with only 2 out of 15 (AalE_VPH_1 and Damh_VPH_2) being flanked by terminal inverted repeats (TIRs) (Fig. 2b). Recently, four marker genes of virophages were established (Roux et al., 2023): A major capsid protein (MCP, also “hexon”), a minor capsid protein (mCP, also “penton”), a FtsK-HerA family DNA-packaging ATPase (ATPase) and a maturation cysteine protease (PRO). For 8 out of 15 virophages, all four marker genes were detected, while for the remaining 7, one was missing.

The GC content of the virophages was similar to that of known virophages, with low GC values being the norm (median = 31.3%, sd = 9.22%). Interestingly, Vibo_VPH_1 had a high GC content of 61.3%, substantially higher than previously reported virophages (Supplementary Data 1, Tab. S7). However, due to its short genome size (4,311 bp), lack of TIRs and absence of a protease, this sequence likely represents an incomplete virophage. Beyond the genes involved in morphogenesis, we detected genes involved in nucleic acid processing (e.g., His-Me finger endonucleases, H-like ribonucleases, and P-loop containing nucleoside triphosphate hydrolases) and carbohydrate metabolism (a concanavalin A-like glucanase), as well as other diverse hydrolase superfamilies (SGNH- and alpha/beta-hydrolases). Lyne_VPH_2 and Lyne_VPH_3 carry an integrase domain, suggesting the ability to integrate into hosts or NCV genomes.

Similarly diverse PLVs were detected (Fig. 2a). All PLV genomes were present in an excised, linear form, with one (Lyne_PLV_2) being flanked by TIRs (Fig. 2c). Their genomes spanned a wide GC range (24 - 57.5%), but with a clear bimodal distribution: one group of low-GC PLVs clustered tightly around ∼24% (5 genomes between 24 - 24.8%), while high-GC PLVs around ∼40% (5 genomes between 39 - 44.7%). Functional annotation revealed genes involved in energy production (P-loop containing nucleoside triphosphate hydrolases), DNA repair, replication and recombination (DNA/RNA polymerases, GIY-YIG endonuclease, His-Me finger endonucleases, Ribonuclease H-like) and virion packaging (A32 packaging ATPase). While the GIY-YIG endonuclease might be involved in integration processes, as is the case in many NCV-associated inteins and introns (Filée, 2018; Gallot-Lavallée et al., 2023), we notably did not detect integrases or recombinases on any PLV.

Virophages and PLVs were detected in 21 out of 23 wastewater treatment plants (Fig. 2d). This pattern largely coincides with the presence or absence of NCVs, with two exceptions: At station *Hirt*, NCVs were present, but no virophages or PLVs were detected, while at station *AalW,* PLVs occurred in the absence of NCVs.

### Interaction between microbes, nucleocytoviruses, polinton-like viruses and virophages

A multi-layered interaction network was constructed to connect NCVs, virophages, PLVs, and collapsed microbial contigs. The latter were collapsed into clusters with a mash distance of 0.05 to reduce redundant contigs into a single microbial cluster. The network’s connections (edges) were defined based on three distinct interaction layers: (i) center-log-ratio (CLR) transformed positive and negative pairwise co-occurrence, when two entities tend to occur together (positive) or exclude each other (negative) across samples (edge weight = 1 to 5, see methods), (ii) integration events, e.g., a virophage integrating into an nucleocytoviral or microbial contig (edge weight = 100), and (iii) gene sharing events, when two copies of the same gene were detected on two separate entities (edge weight = 1). The resulting network comprised 23,611 nodes, including 61 NCVs, 15 virophages and 14 PLVs, connected by 977,398 edges. The network exhibits a heavy-tailed degree distribution with hubs, though formal power-law fitting did not yield conclusive evidence of scale-free topology (likelihood ratio test: p = 0.83; Supplementary Figures 1, Fig. 12). To illustrate general patterns of connectivity, we compared two measures of centrality across the different contig types: *degree* and *betweenness*, the former being a measure for *how many other nodes node X is directly connected to* (Fig. 3a), while the latter represents a measure for *how often node X occurs on all shortest paths between all nodes* (Fig. 3b). Therefore, a high *degree* of centrality can be interpreted as an entity directly interacting with many other entities, e.g., a host sharing genes with an NCV and a virophage. A node with a high level of *betweenness* facilitates interactions between other nodes, e.g., an NCV sharing genes with different hosts, which then themselves interact with different entities each. Nodes representing microbial contigs were further subdivided into being directly connected to NCVs on *any* layer (MC NCV-connected, n = 265) and regular microbial contigs (MC, n = 22572).

**Figure 3:**
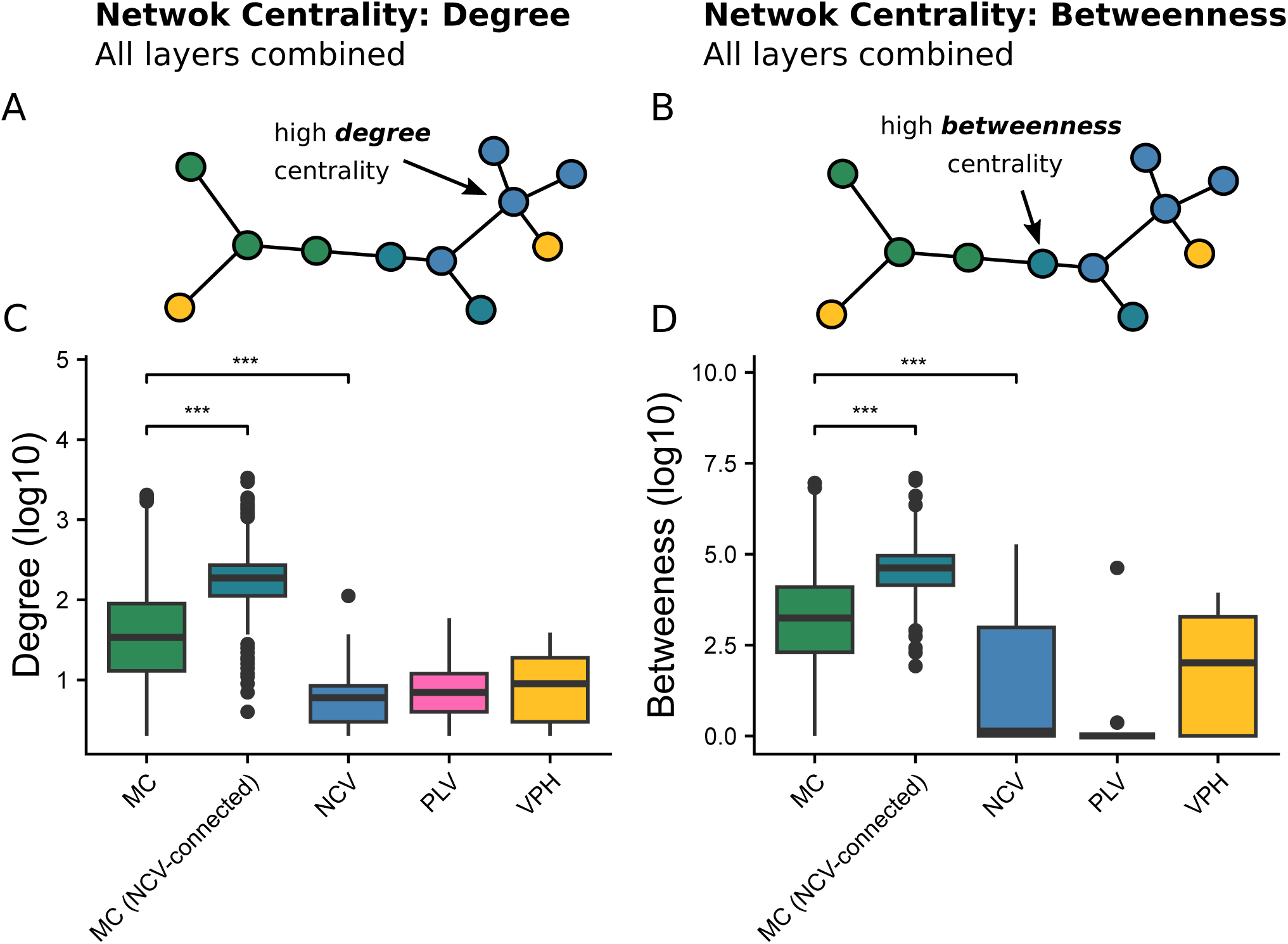
Network centrality measures. This figure illustrates the centrality of nodes within the gene-sharing layer of the multilayered network. Panels A and C are conceptual diagrams showing how degree centrality (Panel A) and betweenness centrality (Panel C) are measured in a network. The distributions of degree centrality (Panel B) and betweenness centrality (Panel D) across different contig types in the analyzed network are shown. The dataset was randomly subsampled to 10% of all gene-sharing connections, due to computational limitations. Pairwise statistical significance was assessed using the Wilcoxon rank-sum test, with stars indicating significance levels (***P < 0.001). MC - microbial contig; NCV - giant virus / nucleocytovirus; PLV - polinton-like virus; VPH - virophage.

Overall, microbial contigs dominated network centrality across both measures (Fig. 3c and 3d): They showed a 10-fold higher degree centrality than NCVs (mean_degree_NCV_ = 8.72, mean_degree_MC_ = 83.2). Virophages and PLVs occupied an intermediate position: While their degree centrality was comparable to that of NCVs (mean_degree_PLV_ = 13, mean_degree_VPH_ = 12.8), their betweenness centrality substantial lower (mean_betweenness_PLV_ = 4,662, mean_betweenness_VPH_ = 1,427 and mean_betweenness_NCV_ = 7,560), suggesting they maintain direct connections to multiple partners but rarely serve as bridges between distinct network components. Interestingly, as soon as a microbial contig is directly connected with at least a single NCV (e.g., co-infecting symbiont-virus relationship), it itself assumes a much more central position within the network, even compared to NCVs (mean_degree_MC_connected_ = 289, Fig. 3c). This dynamic holds also true when comparing the *betweenness* centrality where NCV-connected microbial contigs were again the most central entity type (mean_betweenness_MC_connected_ = 214,554 substantially exceeding both regular MCs (mean_betweenness_MC_ = 24,178) and NCVs (mean_betweenness_NCV_ = 7,560, Fig. 3d). Together, these results indicate that NCVs mediate the centrality of key interaction-hubs, and entities associated with NCVs strongly coincide with more central positions.

Community detection using the Louvain algorithm (resolution = 10) identified densely connected subclusters within the network. When examined in more detail, the network revealed multiple interactions, connecting NCVs, virophages, PLVs and sympatric bacteria. First, virophages and PLVs were often closely associated with NCVs. For example, the cluster around virophage Aved_VPH_2 (Supplementary Figures 2, Fig. 45), contained a group of tightly connected *Imitervirales* linked by gene-sharing and positive (spearman ρ > 0.6, for both long- and short-read data) co-occurrence edges. Aved_VPH_2 was linked to one of these NCVs (OdNE_NCV-Im_1) by a gene-sharing edge and both layers of integration, meaning that read-overhangs mapped to the latter were successfully mapped to the former. Virophage Damh_VPH_2 was connected to Lyne_NCV-Im_1 (Supplementary Figures 2, Fig. 51) by a positive co-occurrence signal. Other virophages and PLVs were also connected to non-viral, microbial contigs: For example, Vibo_VPH_1 and a close cluster of bacteria classified as *Accumulibacter sp.* showed a strong inverse (spearman ρ < -0.6) abundance relationship (Supplementary Figures 2, Fig. 62), while additional positive correlations between PLVs and microbial contigs (Spearman ρ > 0.6) were identified in the network analysis, though individual visualization of these subclusters was limited by size thresholds (<500 nodes) and community detection parameters. Lastly, virophages and PLVs across multiple interaction clusters suggest integration events into microbial contigs, linking them to bacteria based on read-mapping data (Supplementary Figures 2, Fig. 41, 47, 48, 53). A full characterisation and validation of these numerous possible integration events was beyond the scope of the present study and warrants dedicated investigation in future work.

Lastly, NCVs were often found to be connected by negative and positive co-occurrence signals with microbial contigs. For instance, the pandoravirus Fred_NCV-Pa_1 showed positive correlations in abundance with *Saprospiraceae sp.*, while Aved_NCV-As_1 correlated positively with *Sulfuritalea* and Bjer_NCV-As_2 with *Gemmatimonadata* (Supplementary Figures 2, Fig. 15, 2, 3). Conversely, inverse abundance relationships were observed between Bjer_NCV-Pi_2 and *Anaerolina*, as well as between the Hirt_NCV-Im_1 and the same bacterial lineage (Supplementary Figures 2, Fig. 6, 19).

## Discussion

In this study, we recovered high-quality near-complete or complete NCVs, virophages and PLVs from wastewater treatment plants in Denmark. In contrast to similar metagenomic approaches, the number of NCV contigs identified in this study is comparatively small (Rigou et al., 2022; Schulz et al., 2022; Bhattacharjee et al., 2023; Wu et al., 2023; Perini et al., 2024; Minch and Moniruzzaman, 2025). In fact, 9,619 contigs exhibiting NCV signatures were initially identified, but after applying stringent quality criteria, this number was reduced to only 61 high-quality genomes. Focusing on this curated subset of genomes ensured that no false-positive NCV contigs were included in the analyses and provided a reliable basis for exploring NCV interactions with other entities in the dataset.

The identified NCVs displayed remarkable diversity, with genome lengths spanning from a few thousand base pairs to the megabasepair range, characteristic of the phylum. Consistent with the mosaic-like nature of NCV genomes, their gene repertoires reflected functional diversity and complex evolutionary histories. 26 of the identified NCVs carried tRNA genes, which play a potential role in overcoming codon usage mismatches between viruses and hosts (Zhang et al., 2024; Willemsen et al., 2025) and/or sustain translation as host machinery degrades (Yang et al., 2021). Nine NCVs encoded a Thr/Ser kinase. Previously, related kinases were found to be involved in viral protein phosphorylation (Jacob et al., 2011); however, the closest structural homolog was a kinase described as a defense system in *E. coli*, where it triggers an abortive infection in phage-infected cells (Vassallo et al., 2022). While the horizontal acquisition of such a kinase of bacterial origin seems possible, its function remains unclear in this context.

The diversity of the recovered NCVs was further highlighted by seven novel *Mriyaviricetes* genomes, whose major capsid proteins were highly divergent and were detected only through structural prediction. Their coding potential was even more enigmatic, as only for a fraction (as low as ∼5%) of the predicted ORFs homologs were found in databases.

The diversity of virophages and PLVs was equally striking. Their genomes contain multiple indicators of potential environmental interactions, shared with hosts and co-infecting NCVs. For example, the hydrolases found on the newly identified virophages can potentially be used to cleave host or viral macromolecules from co-infecting NCVs to acquire building blocks. At the same time, the His-Me finger endonuclease and ribonucleases might function to modulate gene expression or to degrade nucleic acids to suppress competition and ensure the virophage’s own replication (Filée, 2018). This extensive genomic diversity reflects the dynamic evolutionary history of these viruses, shaped by frequent horizontal gene transfer events fueled by their close association with microbial hosts and co-infecting NCVs and other microbes. Virophages and PLVs have been shown to exist extrachromosomally or in a proviral state integrated into the genomes of the NCVs or their microbial hosts (La Scola et al., 2008; Fischer and Suttle, 2011; Fischer and Hackl, 2016; Koslová et al., 2024). Surprisingly, for only two of the newly identified virophages an integrase domain was detected, and two virophages and one PLV had terminal inverted repeats. Yet no integrated virophage or PLV was found in the dataset. Whether this hints towards an extrachromosomal lifestyle, the reliance on integrases of their host or a co-infecting NCV remains unclear. However, some form of repeated integration and excision mechanisms seems likely as virophages and PLVs consistently have been found to integrate (Bellas et al., 2023; Widen et al., 2023) and read-level evidence from one subcluster suggested a possible integration event between a virophage closely related to Aved_VPH_2 and a co-occurring Imitervirales NCV closely related to OdNE_NCV-Im_1, putatively capturing a transient integration state in a subpopulation of the NCV.

Evidence from the limited number of isolated cell-NCV-virophage systems suggests that the interactions can lead to a wide range of ecological dynamics: While the virophage *Mavirus cafeteriae* significantly reduces replication and virion production of its co-infecting *Rheavirus sinusmexicani* counterpart (Fischer and Suttle, 2011), the virophage *Sputnikvirus zamilonense* appears not to impact the replicative ability of its viral host (Gaia et al., 2014), with other systems falling at intermediate points along this spectrum (La Scola et al., 2008; Yau et al., 2011). On a community level, (hyper)parasitic elements can act as defensive systems (Oliveira et al., 2019; Koonin et al., 2020; Koslová et al., 2024). Yet, virophages and PLVs remain strictly dependent on co-infecting NCVs for replication. Consistent with this, virophages and PLVs were detected across 21 of 23 sampling stations, coinciding with NCV presence, with only two exceptions in either direction and multiple virophages and PLVs forming co-occurrence pairs with NCVs in the network (Supplementary Figures 2, Fig. 22, 32, 48).

Beyond well-described host-NCV and virophage-NCV interactions, NCVs have been shown to associate with sympatric bacteria, adding a further layer of ecological and evolutionary interaction to already intricate and complex systems (Arthofer et al., 2022; Schulz et al., 2024). Our network revealed multiple NCV-bacteria pairs, linked by positive and negative co-occurrence signals. Notably, these associations involve bacterial lineages, particularly *Saprospiraceae* and *Anaerolineaceae*, that are abundant core taxa in activated sludge communities worldwide (Xu et al., 2018; Kondrotaite et al., 2022). Unlike the *Chlamydiae* in the Arthofer et al. (2022) study, neither species are known protist symbionts, suggesting that mechanisms distinct from defensive symbiosis may underlie these NCV-bacteria co-occurrence patterns and warrant further investigation.

Furthermore, the negative association between virophage Vibo_VPH_1 and *Accumulibacter*, another widespread polyphosphate-accumulating species in WWTPs (Singleton et al., 2021; Kondrotaite et al., 2022), raises the intriguing possibility of quadripartite systems involving protist hosts, NCVs, virophages, and bacteria in WWTPs. To our knowledge, no such system has been described, yet it seems plausible that they exist in nature.

Taking these connections together, NCVs emerge as focal points of multi-level ecological interactions: their diverse and highly mobile gene repertoires, their ability to modulate community-scale evolutionary and ecological dynamics through interactions with hyperparasitic elements and co-infecting bacteria, and their physical proximity to multiple community members position them as central interaction partners that draw otherwise unconnected taxa into dense network clusters. Our network analysis captures this dynamic directly: while NCVs themselves do not rank among the most central nodes by either degree or betweenness centrality, NCV-connected microbial contigs assume dramatically elevated centrality compared to both unconnected microbes and the NCVs themselves (3.5 fold and 9.1 fold increase in comparison to unconnected microbes, for degree and betweenness, respectively), positioning NCVs not as hubs, but as key mediators of interaction.

Although this analysis demonstrates the highly central and multifaceted role of NCVs in structuring microbial community interactions within activated sludge, important limitations remain that constrain our conclusions. First, the importance of MGEs such as transpovirons, introns, inteins, and miniature inverted-repeat transposable elements (MITEs) in virus-host systems has been demonstrated (Etten and Meints, 1999; Desnues et al., 2012; Sun et al., 2015; Filée, 2018; Gallot-Lavallée et al., 2023), yet these players were omitted as their detailed characterization was beyond the scope of this study. Furthermore, none of the microbial contigs in our network were classified to be of eukaryotic origin, which leads to a de facto removal of any potential hosts. Most probably, the >100 kbp contig cutoff used for the network, removed shorter fragments of eukaryotic genomes, known to assemble poorly from metagenomes due to large genome size and complexity and lower relative abundance (Delmont et al., 2022). Lastly, the ecological relevance of many putative interaction mechanisms remains uncertain and will require targeted experimental validation, beyond the scope of this study.

Despite these constraints, this study not only provides high-quality NCV, virophage and PLV genomes, but also establishes NCVs as key interaction mediators in wastewater treatment plants, and provides profound guidance for future research: On the one hand, the presented approach provides a practical catalogue of clusters which can be used to identify candidate ecological interactions and isolation targets. On the other hand, it complements established workflows; integrated with temporally and spatially resolved metagenomics, single-cell approaches, or cultivation work, our interaction-centric approach can help trace ecological and evolutionary dynamics across complex microbial communities. Altogether, this study lays out a fundamental framework that advances our understanding of (nucleocyto)viral ecology in wastewater systems and offers a foundation for future investigations into eukaryotic-virus-bacteria-virophage interactions.

## Authors Contributions

Conceptualization: AW, AMM and DL. Data curation: AW, AMM and DL. DL performed the majority of the bioinformatics analyses with assistance from AW and AMM. DL wrote the initial draft, with input and revisions from both AW and AMM. All authors contributed to the interpretation of results, reviewed the manuscript, and approved the final version.

## Acknowledgements

The computational results of this work have been achieved using the Life Science Compute Cluster (LiSC) of the University of Vienna. The authors thank Sean Darcy and Qi Qi for discussions and expertise. This project has received funding from the European Union’s Horizon 2020 research and innovation programme under the Marie Sklodowska-Curie grant agreement No. 891572 and the European Union (ERC, CHIMERA, 101039843). Views and opinions expressed are however those of the author(s) only and do not necessarily reflect those of the European Union or the European Research Council Executive Agency. Neither the European Union nor the granting authority can be held responsible for them. AW acknowledges funding from the Austrian Science Fund (FWF) [doi.org/10.55776/COE7].

## Data Availability

An interactive network visualization and associated supplementary data are hosted on a GitHub Pages homepage accompanying this publication: https://dluecking.github.io/wwtp_linking_homepage/. Genomes will be deposited in the ENA under project accession PRJEB105214. Assembled genomes, proteins, annotation files, phylogenetic data and network data are provided through Zenodo: 10.5281/zenodo.17735769.

## Additional Data

**Supplementary Data 1:** Tabular overview of NCVs (Table S1), custom hmm bitscore cutoffs (Table S2), *Mriyaviricetes* major capsid protein alignment scores (Table S3), virophages (Table S4), PLVs (Table S5), genome accession of publicly available virophages and PLVs used in this study (Table S6).

**Supplementary Figures 1:** ANI and MASH distances of NCVs recovered in this study (Figure S1), maximum-likelihood trees of NCV core genes (Figure S2-S10), maximum-likelihood trees of *Mriyaviricetes* major capsid protein (Figure S11) and log-log degree distribution of the established interaction network (Figure S12).

**Supplementary Figures 2:** Network cluster visualization showing connections among *Nucleocytoviricota, Preplasmiviricota* (Virophages, Polinton-like viruses) and microbial (MC) contigs.

