## Supplementary Figures 1 for "A multilayered network reveals the centrality of newly discovered *Nucleocytoviricota* in wastewater treatment plant communities"

**List of Figures:**

**Supplementary Fig. 1, page 3.** MASH output of average nucleotide identity (ANI) and clustering of the identified *Nucleocytoviricota.* Vertical dashed line indicates >95 % identity.

**Supplementary Fig. 2, page 4.** Maximum-likelihood phylogeny of NCVs based on MCP (GVOGm0003). ML tree constructed using IQ-TREE (v2.4.0) and visualized with iTOL (v7.5.1), midpoint rooted. Black circles indicate branch-support (Bootstrap values > 90). Color stripes indicate order assignment, sequences from this study are not assigned. Note that alternative IDs for sequences from this study were used (Supplementary Data, Table S1).

**Supplementary Fig. 3, page 5.** Maximum-likelihood phylogeny of NCVs based on SFII (GVOGm0013). ML tree constructed using IQ-TREE (v2.4.0) and visualized with iTOL (v7.5.1), midpoint rooted. Black circles indicate branch-support (Bootstrap values > 90). Color stripes indicate order assignment, sequences from this study are not assigned. Note that alternative IDs for sequences from this study were used (Supplementary Data, Table S1).

**Supplementary Fig. 4, page 6.** Maximum-likelihood phylogeny of NCVs based on RNAPS (GVOGm0022). ML tree constructed using IQ-TREE (v2.4.0) and visualized with iTOL (v7.5.1), midpoint rooted. Black circles indicate branch-support (Bootstrap values > 90). Color stripes indicate order assignment, sequences from this study are not assigned. Note that alternative IDs for sequences from this study were used (Supplementary Data, Table S1).

**Supplementary Fig. 5, page 7.** Maximum-likelihood phylogeny of NCVs based on RNAPL (GVOGm0023). ML tree constructed using IQ-TREE (v2.4.0) and visualized with iTOL (v7.5.1), midpoint rooted. Black circles indicate branch-support (Bootstrap values > 90). Color stripes indicate order assignment, sequences from this study are not assigned. Note that alternative IDs for sequences from this study were used (Supplementary Data, Table S1).

**Supplementary Fig. 6, page 8.** Maximum-likelihood phylogeny of NCVs based on PolB (GVOGm0054). ML tree constructed using IQ-TREE (v2.4.0) and visualized with iTOL (v7.5.1), midpoint rooted. Black circles indicate branch-support (Bootstrap values > 90). Color stripes indicate order assignment, sequences from this study are not assigned. Note that alternative IDs for sequences from this study were used (Supplementary Data, Table S1).

**Supplementary Fig. 7, page 9.** Maximum-likelihood phylogeny of NCVs based on TFIIB (GVOGm0172). ML tree constructed using IQ-TREE (v2.4.0) and visualized with iTOL (v7.5.1), midpoint rooted. Black circles indicate branch-support (Bootstrap values > 90). Color stripes indicate order assignment, sequences from this study are not assigned. Note that alternative IDs for sequences from this study were used (Supplementary Data, Table S1).

**Supplementary Fig. 8, page 10.** Maximum-likelihood phylogeny of NCVs based on TopoII (GVOGm0461). ML tree constructed using IQ-TREE (v2.4.0) and visualized with iTOL (v7.5.1), midpoint rooted. Black circles indicate branch-support (Bootstrap values > 90). Color stripes indicate order assignment, sequences from this study are not assigned. Note that alternative IDs for sequences from this study were used (Supplementary Data, Table S1).

**Supplementary Fig. 9, page 11.** Maximum-likelihood phylogeny of NCVs based on A32 (GVOGm0760). ML tree constructed using IQ-TREE (v2.4.0) and visualized with iTOL (v7.5.1), midpoint rooted. Black circles indicate branch-support (Bootstrap values > 90). Color stripes indicate order assignment, sequences from this study are not assigned. Note that alternative IDs for sequences from this study were used (Supplementary Data, Table S1).

**Supplementary Fig. 10, page 12.** Maximum-likelihood phylogeny of NCVs based on VLTF3 (GVOGm0890). ML tree constructed using IQ-TREE (v2.4.0) and visualized with iTOL (v7.5.1), midpoint rooted. Black circles indicate branch-support (Bootstrap values > 90). Color stripes indicate order assignment, sequences from this study are not assigned. Note that alternative IDs for sequences from this study were used (Supplementary Data, Table S1).

**Supplementary Fig. 11, page 13.** Unrooted maximum-likelihood phylogeny of detected *Mriyaviricetes* and sequences from Yutin et al., 2024, based on the major capsid protein.

**Supplementary Fig. 12, page 14.** Log-log degree distribution of the established interaction network.

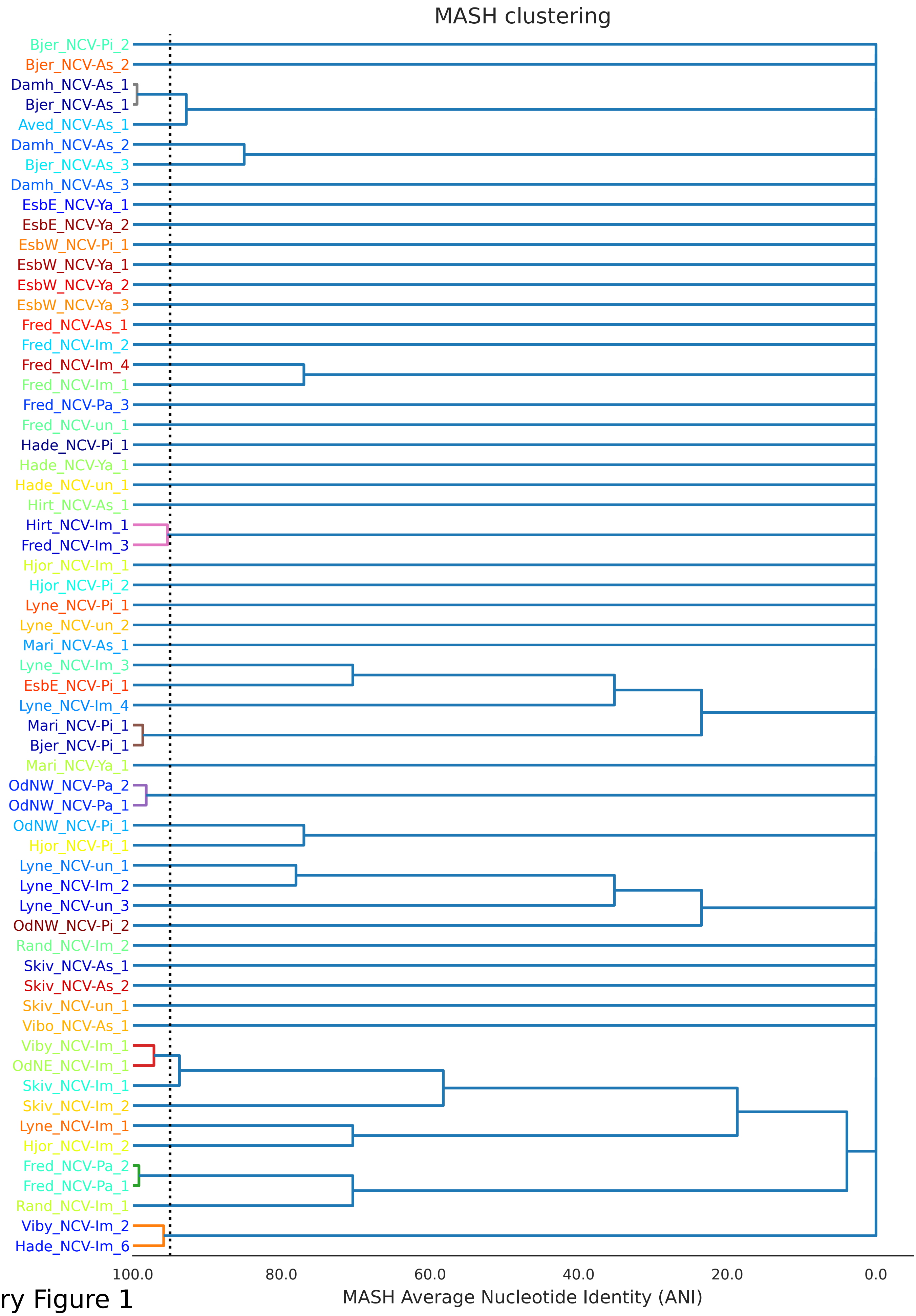


Supplementary Figure 1

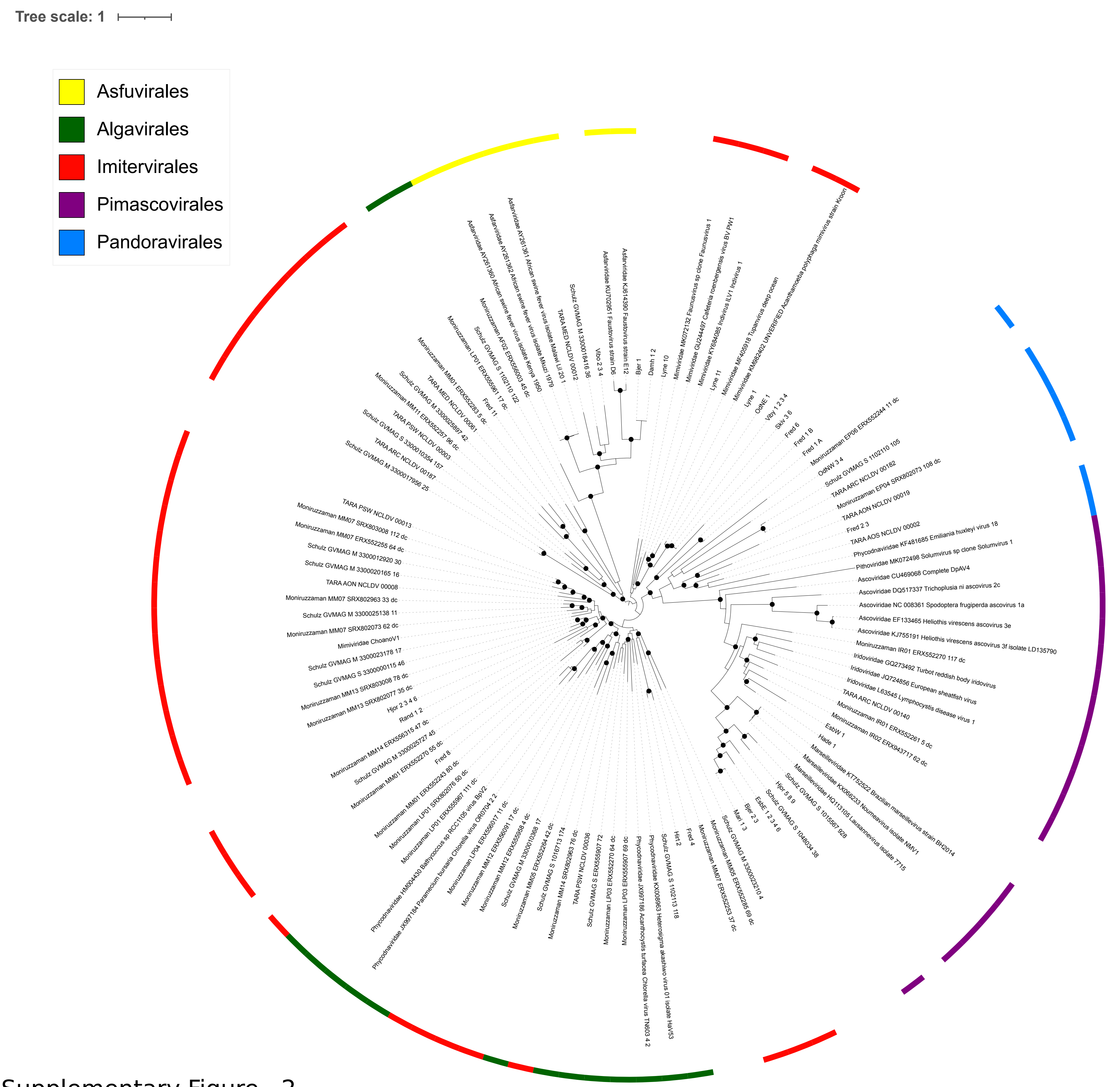


Supplementary Figure 2

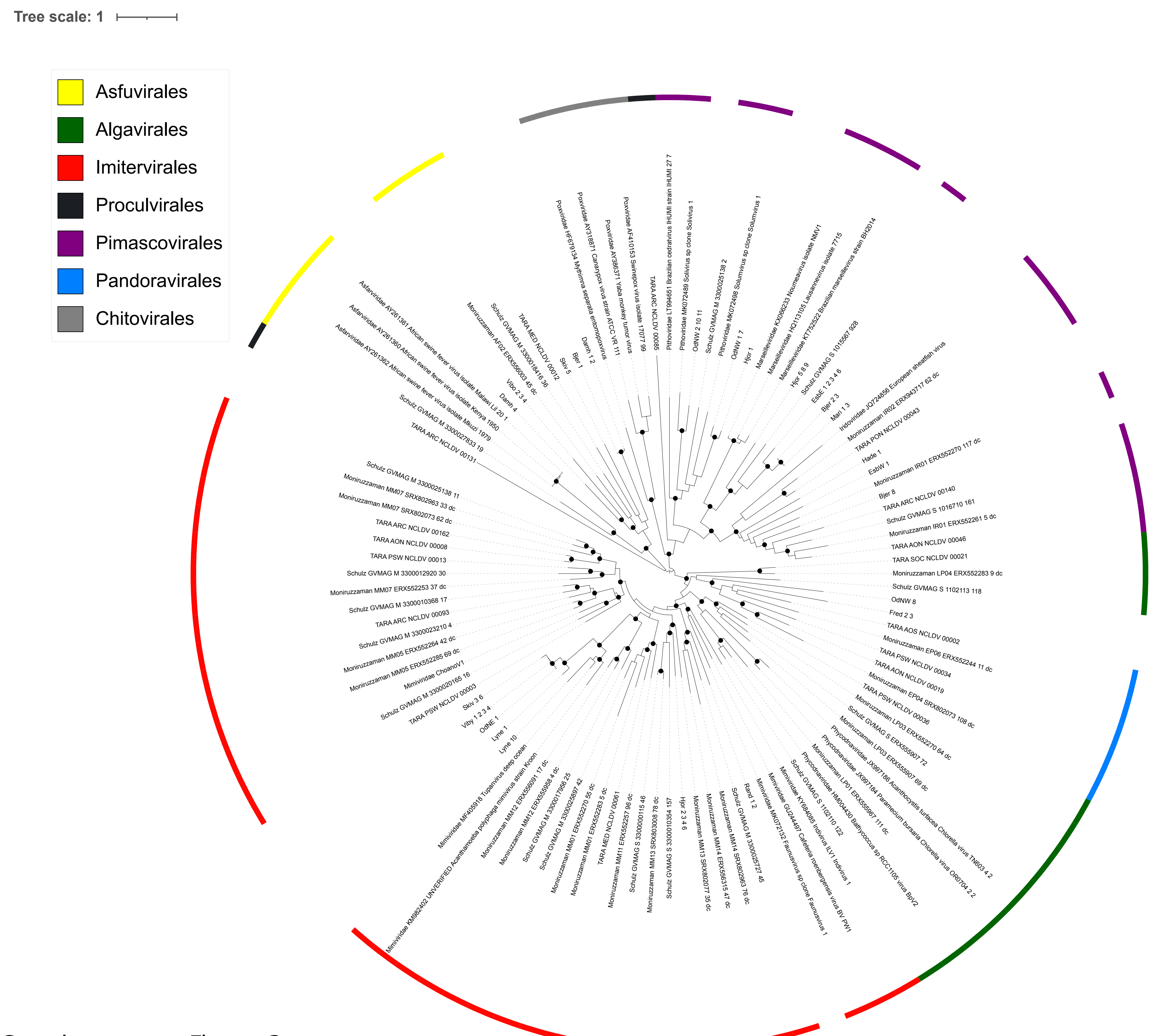


Supplementary Figure 3

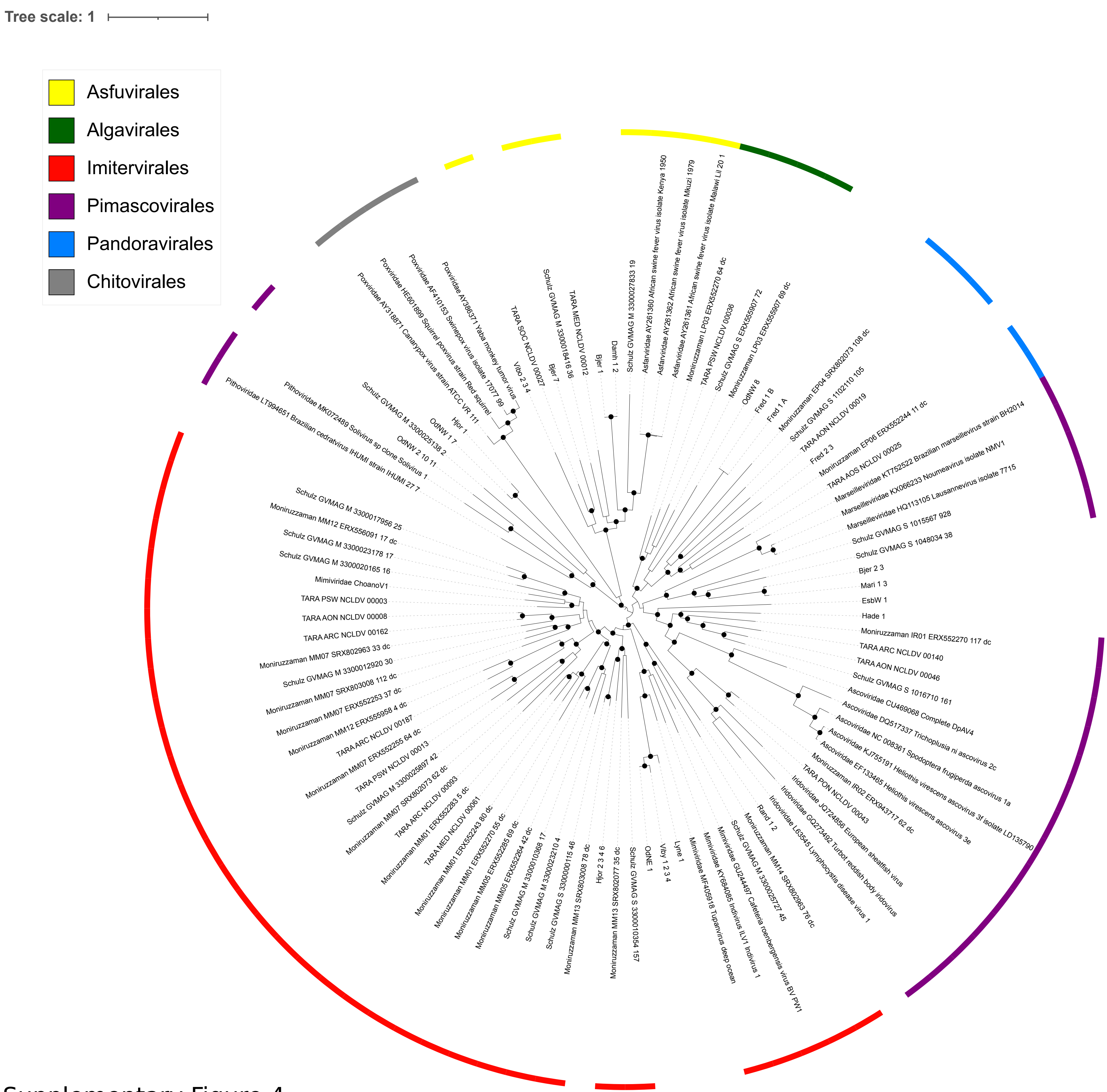


Supplementary Figure 4

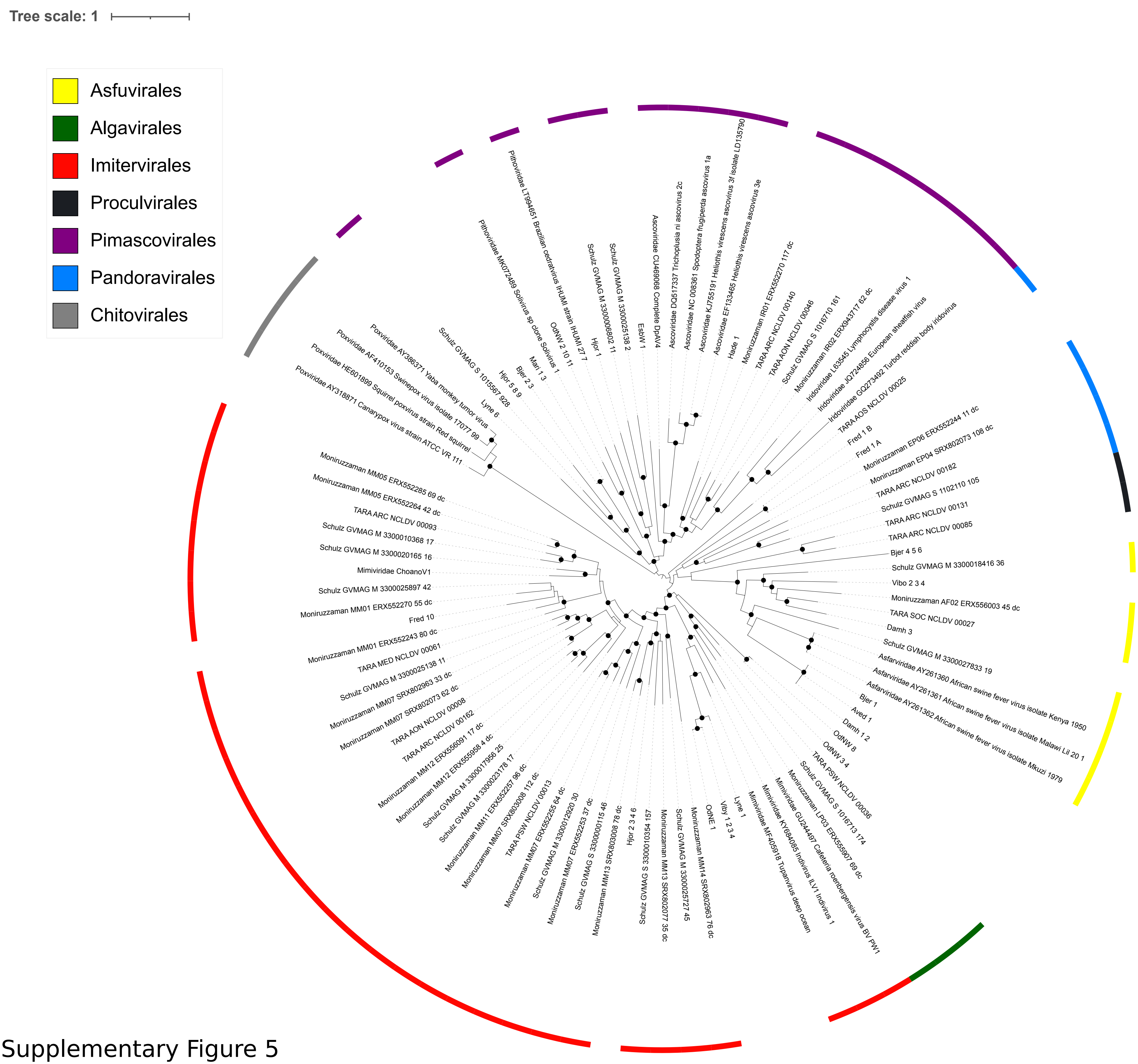


Supplementary Figure 5

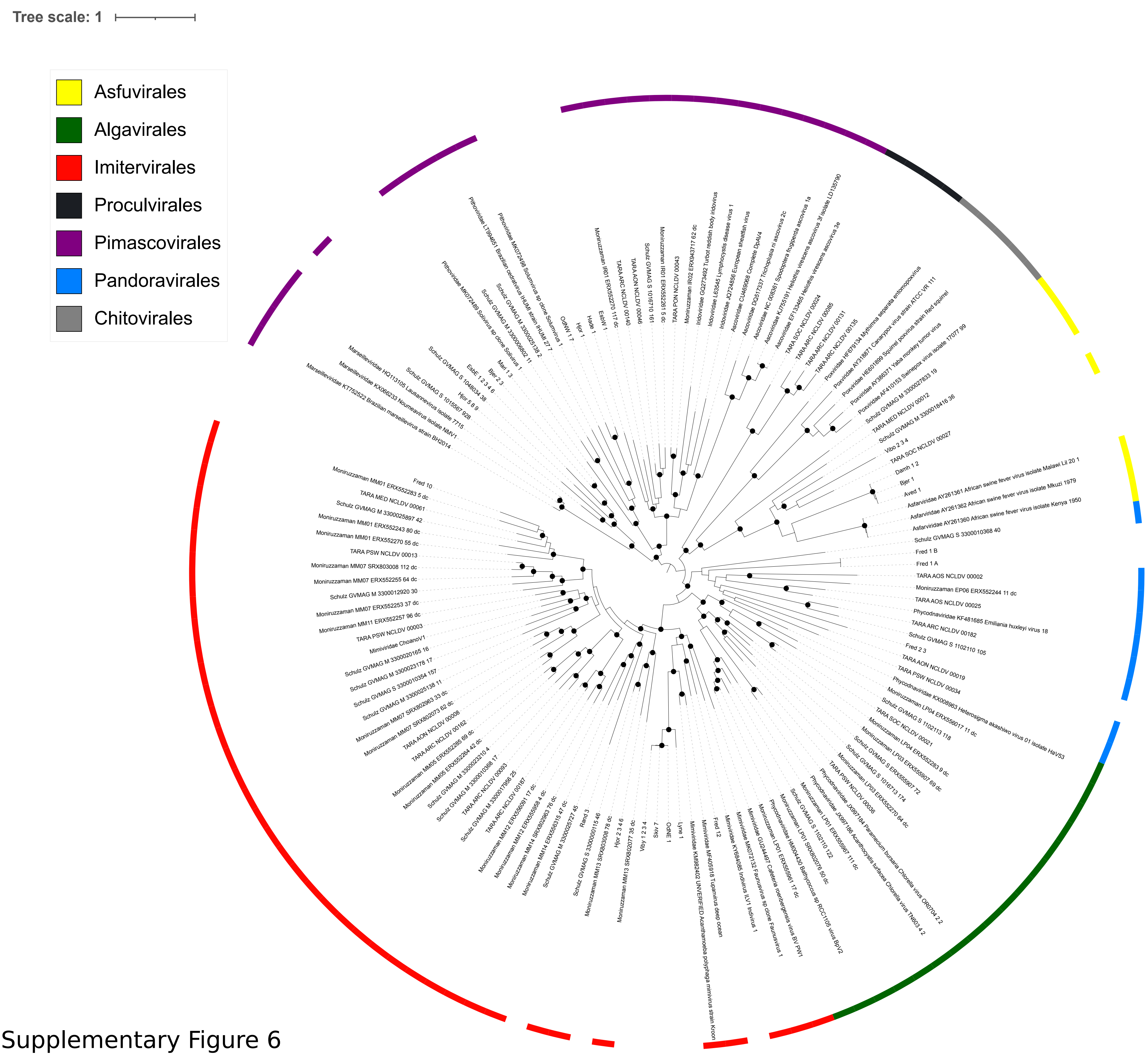


Supplementary Figure 6

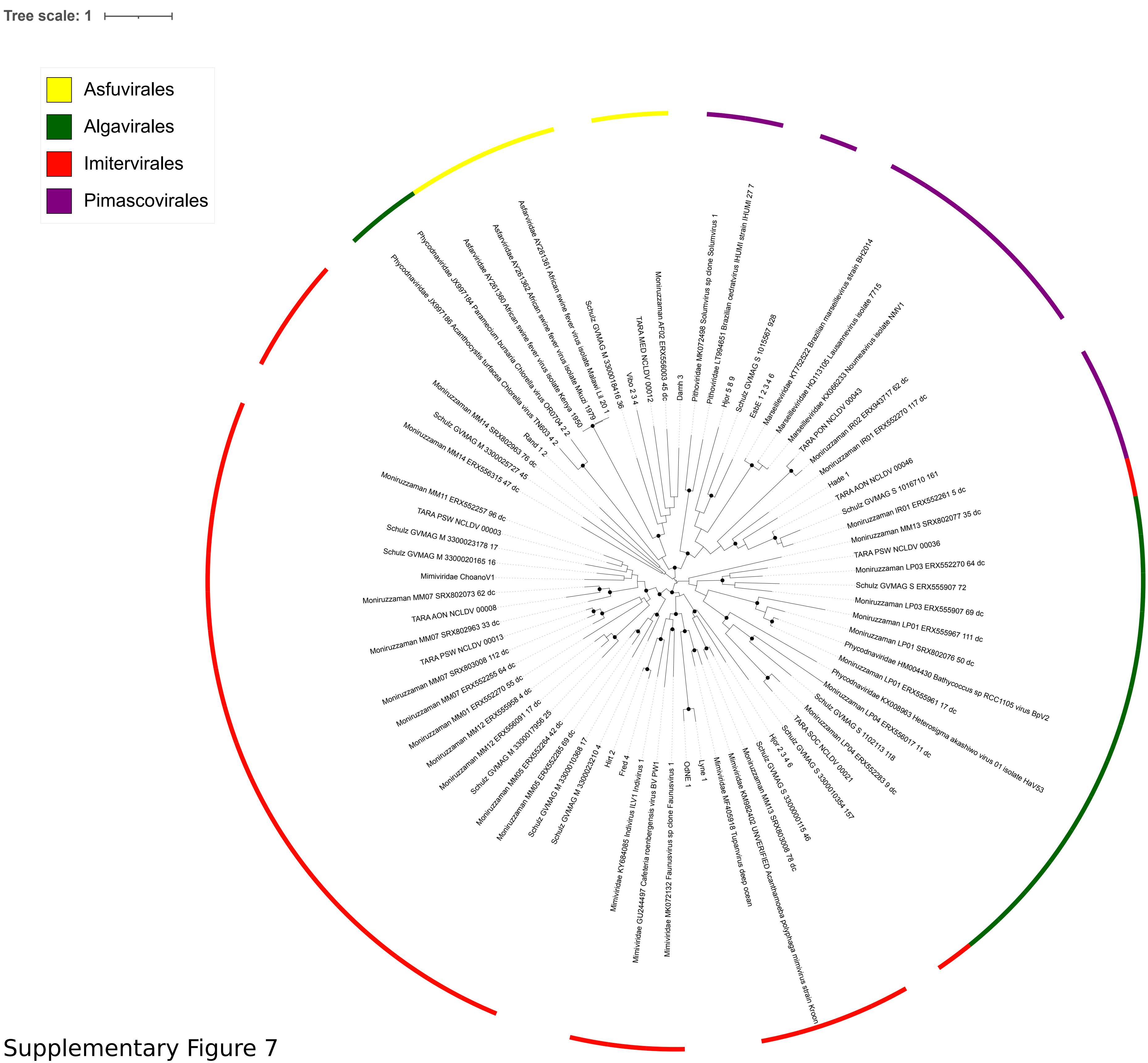


Supplementary Figure 7

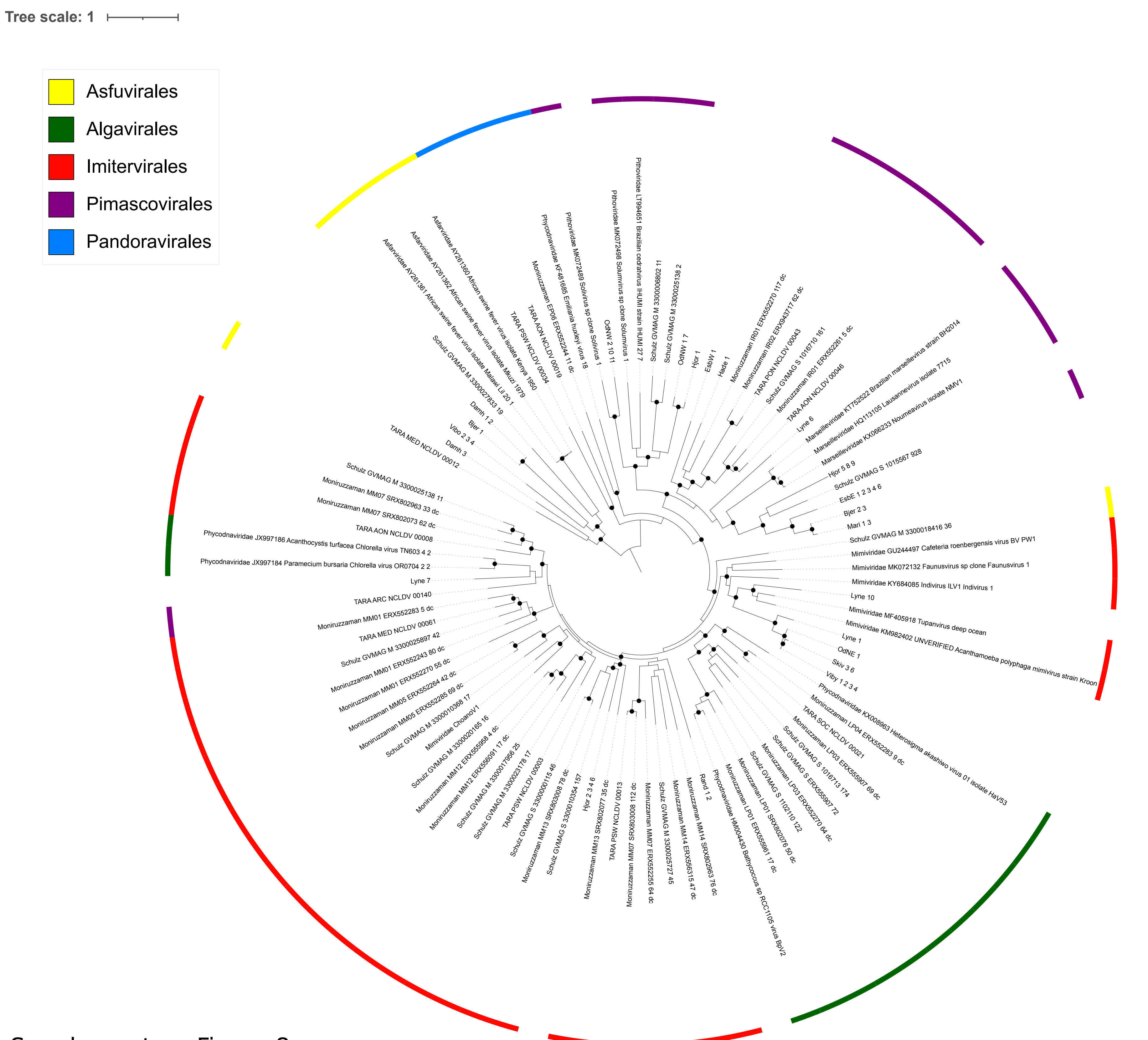


Supplementary Figure 8

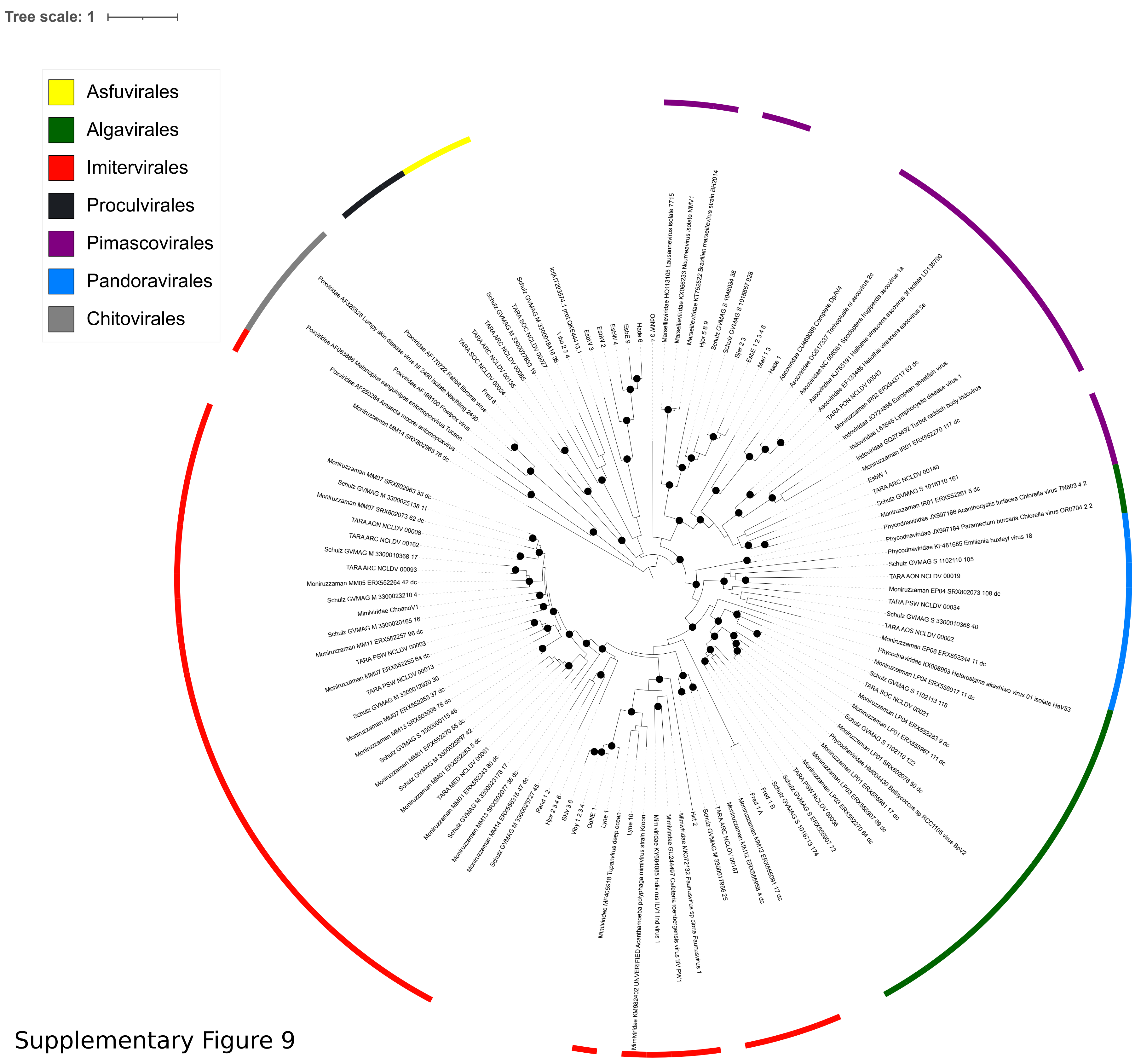


Supplementary Figure 9

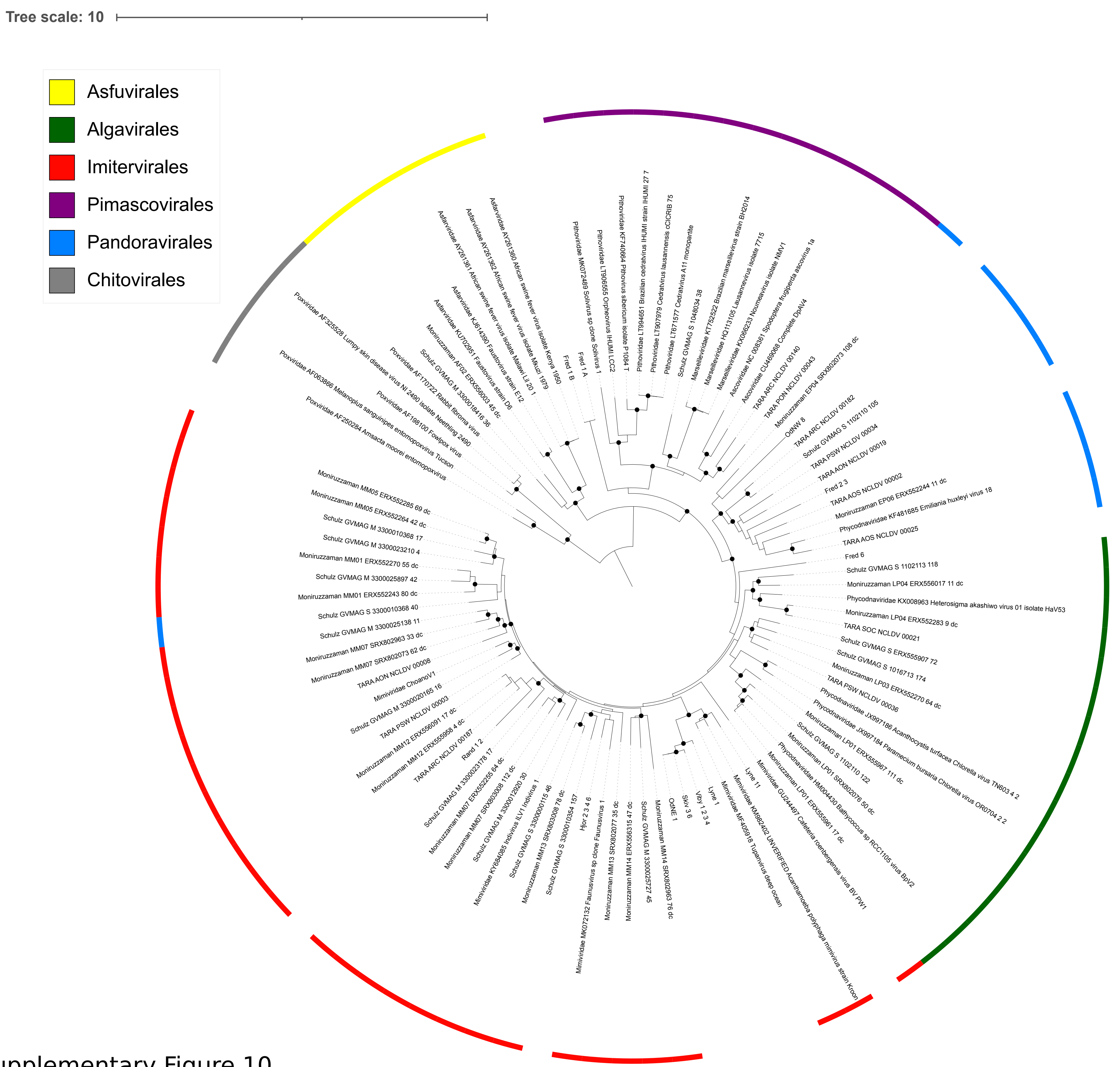


Supplementary Figure 10

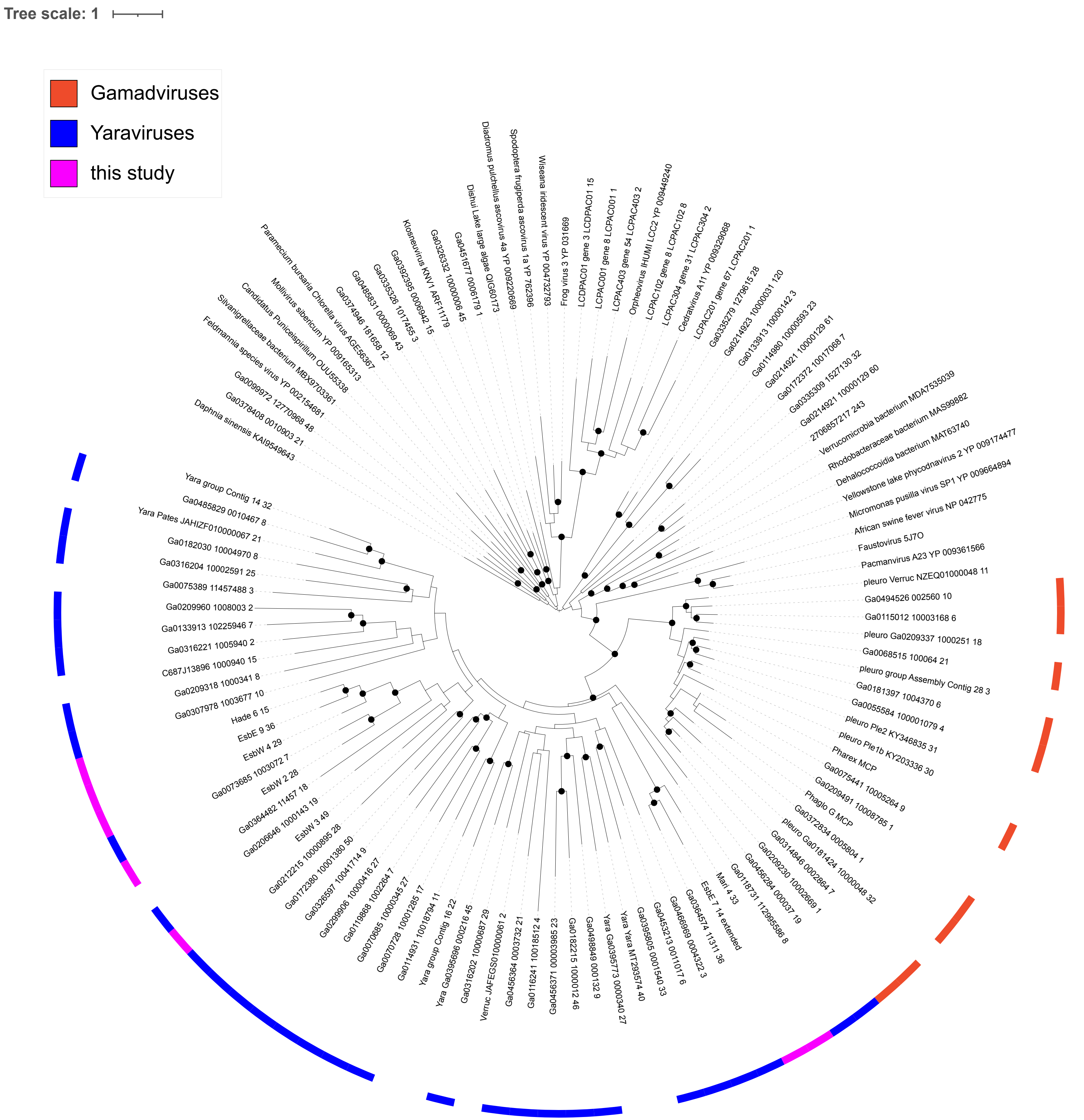


Supplementary Figure 11

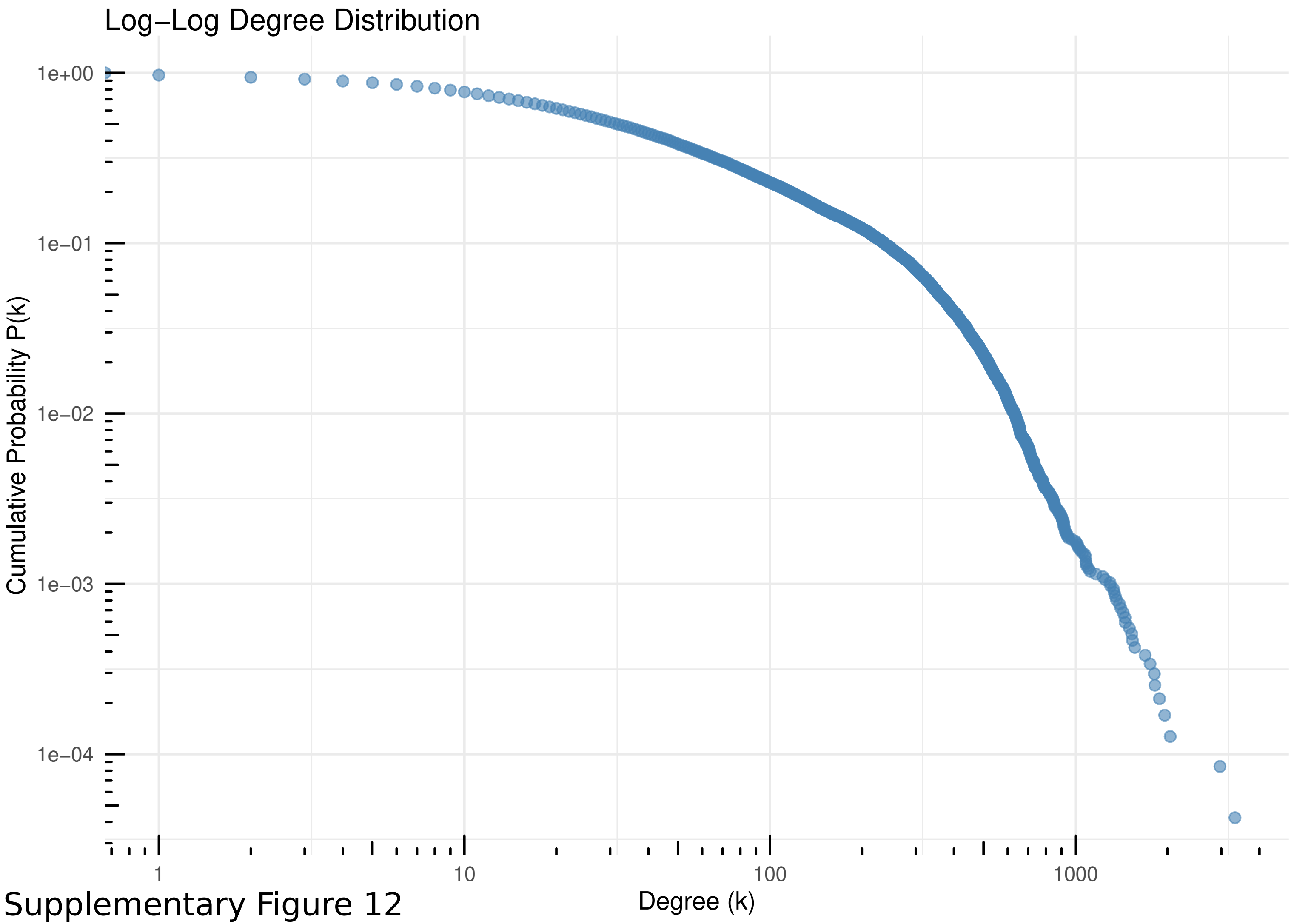


Supplementary Figure 12
