## Supplementary Figures 2 for "A multilayered network reveals the centrality of newly discovered *Nucleocytoviricota* in wastewater treatment plant communities"

**Figure Description:**

Network cluster visualization showing connections among *Nucleocytoviricota, Preplasmiviricota* (Virophages, Polinton-like viruses) and microbial (MC) contigs. Each figure is centered on a specific node and shows all nodes within the same Louvain-group. Mind that this can result in cases where two figures show the same plot, but are centered around two different nodes if these two nodes share the same Louvain-group. Only clusters with <500 edges are included for visualization reasons. Each panel depicts a distinct layer of the multi-layered network, revealing different mechanisms of genetic and ecological association: Gene sharing, co-occurance (positive / negative, Illumina read-mapping / ONT read-mapping) and integration (boundary, middle). For details please refer to the method section of the publication (section Network Analysis).

**List of Figures:**

**NCVs**


Fig. 1, page 4. Cluster plot surrounding the node "Aved_NCV-As_1"
Fig. 2, page 5. Cluster plot surrounding the node "Bjer_NCV-As_1"
Fig. 3, page 6. Cluster plot surrounding the node "Bjer_NCV-As_2"
Fig. 4, page 7. Cluster plot surrounding the node "Bjer_NCV-As_3"
Fig. 5, page 8. Cluster plot surrounding the node "Bjer_NCV-Pi_1"
Fig. 6, page 9. Cluster plot surrounding the node "Bjer_NCV-Pi_2"
Fig. 7, page 10. Cluster plot surrounding the node "Damh_NCV-As_1"
Fig. 8, page 11. Cluster plot surrounding the node "Damh_NCV-As_2"
Fig. 9, page 12. Cluster plot surrounding the node "Damh_NCV-As_3"
Fig. 10, page 13. Cluster plot surrounding the node "EsbE_NCV-Pi_1"
Fig. 11, page 14. Cluster plot surrounding the node "EsbW_NCV-Pi_1"
Fig. 12, page 15. Cluster plot surrounding the node "EsbW_NCV-Ya_1"
Fig. 13, page 16. Cluster plot surrounding the node "EsbW_NCV-Ya_2"
Fig. 14, page 17. Cluster plot surrounding the node "Fred_NCV-Im_3"
Fig. 15, page 18. Cluster plot surrounding the node "Fred_NCV-Pa_1"
Fig. 16, page 19. Cluster plot surrounding the node "Fred_NCV-Pa_2"
Fig. 17, page 20. Cluster plot surrounding the node "Hade_NCV-Im_6"

Fig. 18, page 21. Cluster plot surrounding the node "Hirt_NCV-As_1"
Fig. 19, page 22. Cluster plot surrounding the node "Hirt_NCV-Im_1"
Fig. 20, page 23. Cluster plot surrounding the node "Hjor_NCV-Im_1"
Fig. 21, page 24. Cluster plot surrounding the node "Hjor_NCV-Pi_1"
Fig. 22, page 25. Cluster plot surrounding the node "Lyne_NCV-Im_1"
Fig. 23, page 26. Cluster plot surrounding the node "Lyne_NCV-Im_2"
Fig. 24, page 27. Cluster plot surrounding the node "Lyne_NCV-Im_3"
Fig. 25, page 28. Cluster plot surrounding the node "Lyne_NCV-Im_4"
Fig. 26, page 29. Cluster plot surrounding the node "Lyne_NCV-un_1"
Fig. 27, page 30. Cluster plot surrounding the node "Lyne_NCV-un_2"
Fig. 28, page 31. Cluster plot surrounding the node "Lyne_NCV-un_3"
Fig. 29, page 32. Cluster plot surrounding the node "Mari_NCV-As_1"
Fig. 30, page 33. Cluster plot surrounding the node "Mari_NCV-Pi_1"
Fig. 31, page 34. Cluster plot surrounding the node "OdNE_NCV-Im_1"
Fig. 32, page 35. Cluster plot surrounding the node "OdNW_NCV-Pa_1"
Fig. 33, page 36. Cluster plot surrounding the node "OdNW_NCV-Pa_2"
Fig. 34, page 37. Cluster plot surrounding the node "OdNW_NCV-Pi_1"
Fig. 35, page 38. Cluster plot surrounding the node "Skiv_NCV-As_2"
Fig. 36, page 39. Cluster plot surrounding the node "Skiv_NCV-Im_1"
Fig. 37, page 40. Cluster plot surrounding the node "Skiv_NCV-Im_2"
Fig. 38, page 41. Cluster plot surrounding the node "Skiv_NCV-un_1"
Fig. 39, page 42. Cluster plot surrounding the node "Vibo_NCV-As_1"
Fig. 40, page 43. Cluster plot surrounding the node "Viby_NCV-Im_1"

**Polinton-like viruses / virophages**

Fig. 41, page 44. Cluster plot surrounding the node "AalE_VPH_1"
Fig. 42, page 45. Cluster plot surrounding the node "AalW_PLV_1"
Fig. 43, page 46. Cluster plot surrounding the node "Aved_PLV_1"
Fig. 44, page 47. Cluster plot surrounding the node "Aved_VPH_1"
Fig. 45, page 48. Cluster plot surrounding the node "Aved_VPH_2"
Fig. 46, page 49. Cluster plot surrounding the node "Bjer_PLV_1"
Fig. 47, page 50. Cluster plot surrounding the node "Damh_VPH_1"
Fig. 48, page 51. Cluster plot surrounding the node "Damh_VPH_2"
Fig. 49, page 52. Cluster plot surrounding the node "Damh_VPH_3"

Fig. 50, page 53. Cluster plot surrounding the node "Fred_VPH_1"
Fig. 51, page 54. Cluster plot surrounding the node "Fred_VPH_2"
Fig. 52, page 55. Cluster plot surrounding the node "Hade_VPH_1"
Fig. 53, page 56. Cluster plot surrounding the node "Lyne_PLV_1"
Fig. 54, page 57. Cluster plot surrounding the node "Lyne_PLV_2"
Fig. 55, page 58. Cluster plot surrounding the node "Lyne_VPH_1"
Fig. 56, page 59. Cluster plot surrounding the node "Lyne_VPH_2"
Fig. 57, page 60. Cluster plot surrounding the node "Lyne_VPH_3"
Fig. 58, page 61. Cluster plot surrounding the node "Mari_PLV_1"
Fig. 59, page 62. Cluster plot surrounding the node "Rand_PLV_1"
Fig. 60, page 63. Cluster plot surrounding the node "Ribe_PLV_1"
Fig. 61, page 64. Cluster plot surrounding the node "Vibo_PLV_1"
Fig. 62, page 65. Cluster plot surrounding the node "Vibo_VPH_1"

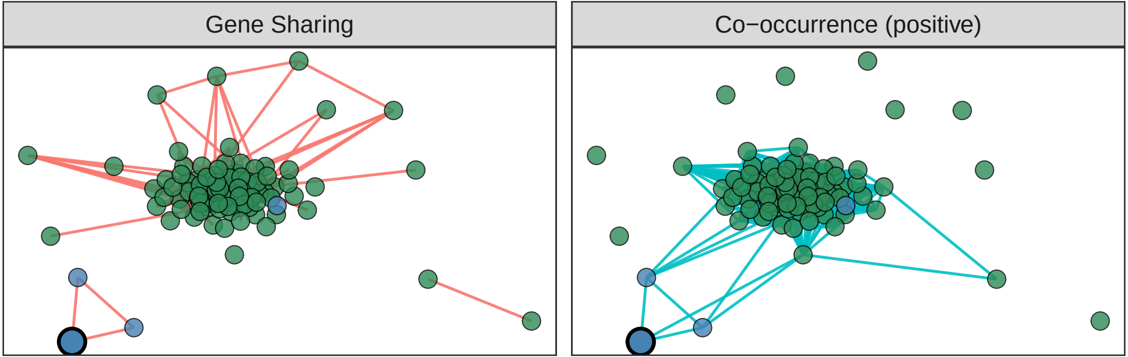
Gene Sharing
Co−occurrence (positive)

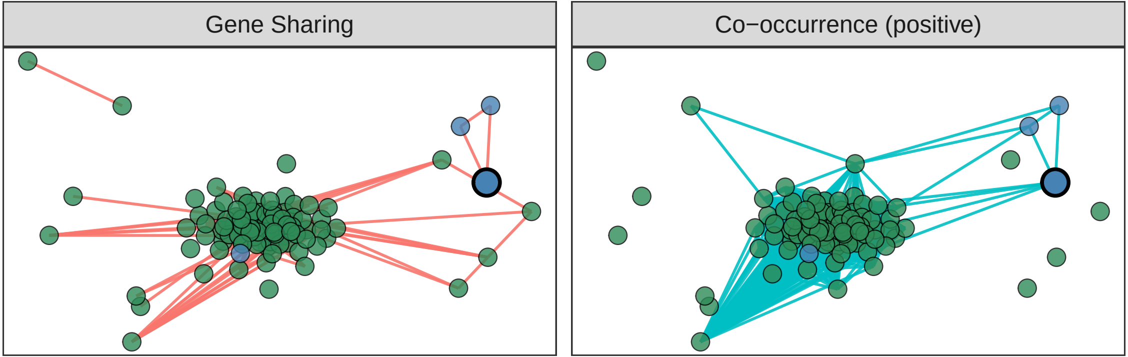
Gene Sharing
Co−occurrence (positive)

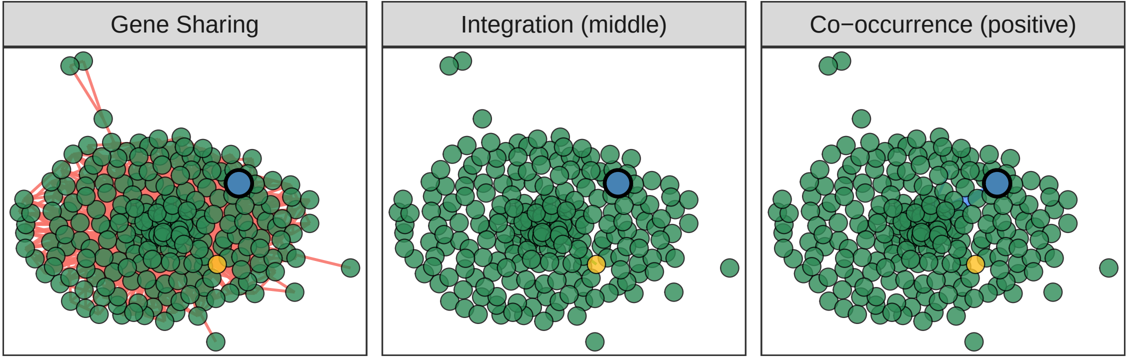
Gene Sharing
Integration (middle)
Co−occurrence (positive)

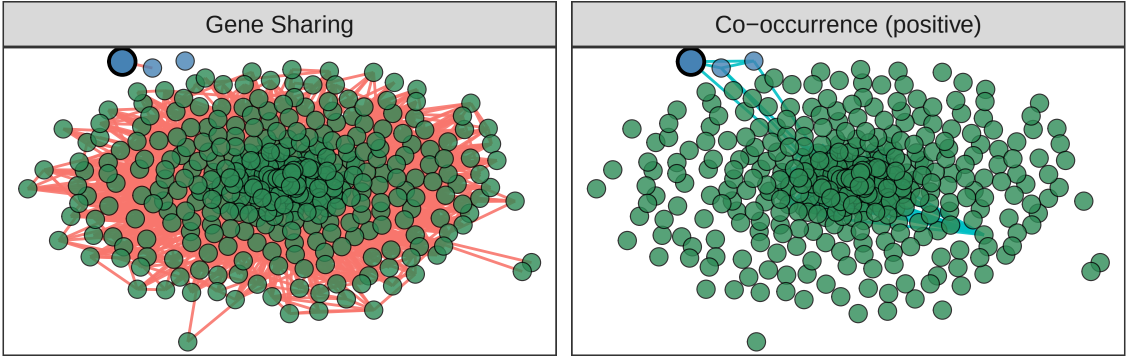

Gene Sharing
Co−occurrence (positive)

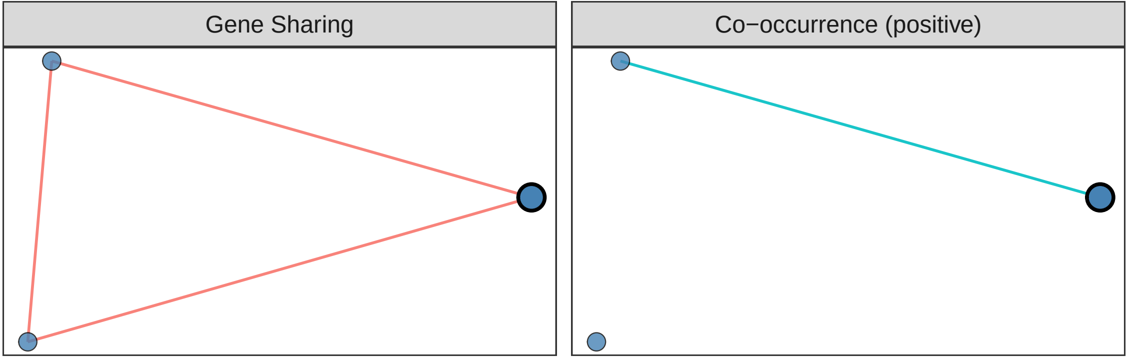
Gene Sharing
Co−occurrence (positive)

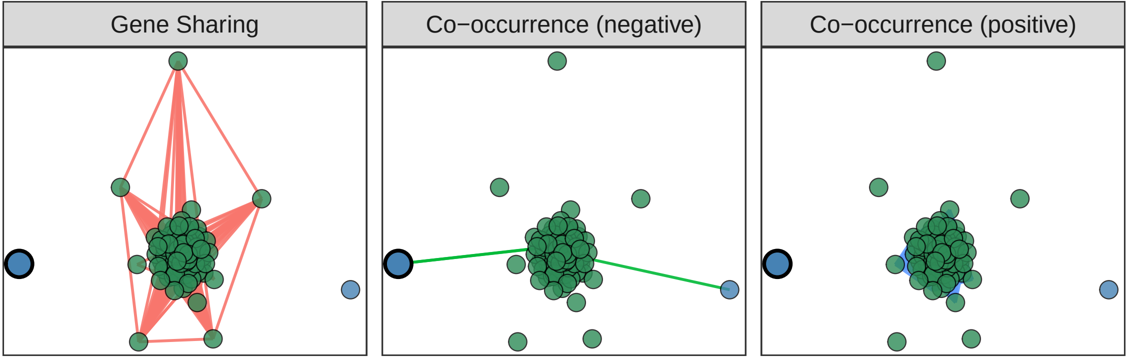
Gene Sharing
Co−occurrence (negative)
Co−occurrence (positive)

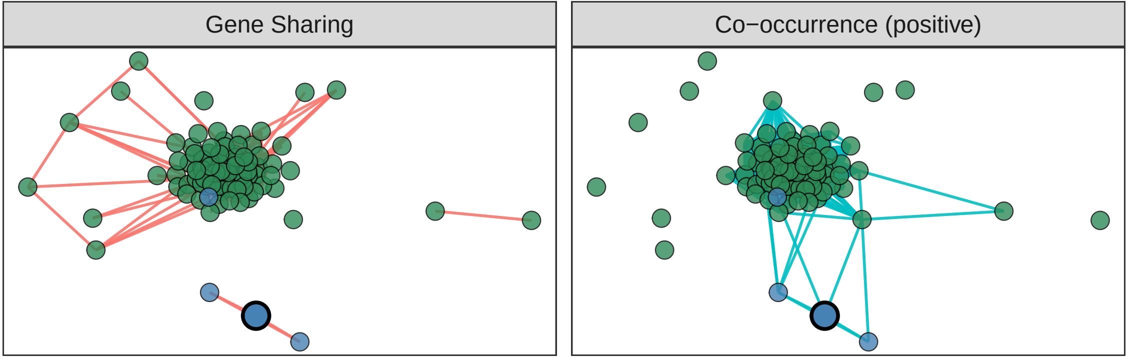
Gene Sharing
Co−occurrence (positive)

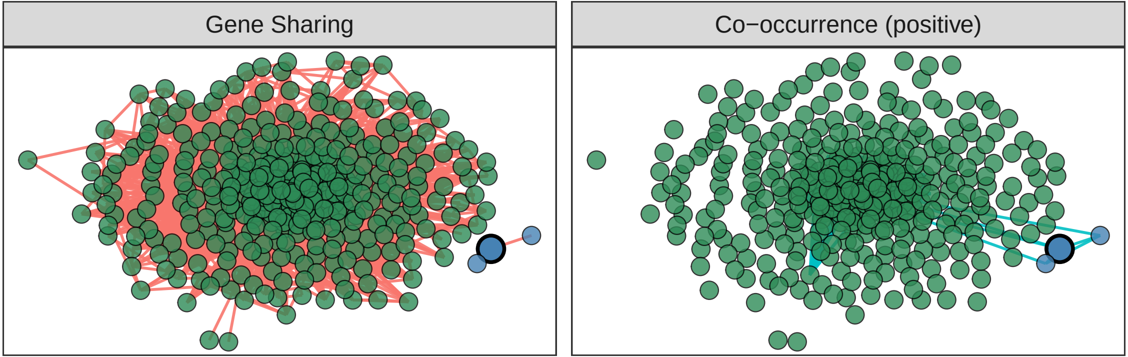
Gene Sharing
Co−occurrence (positive)

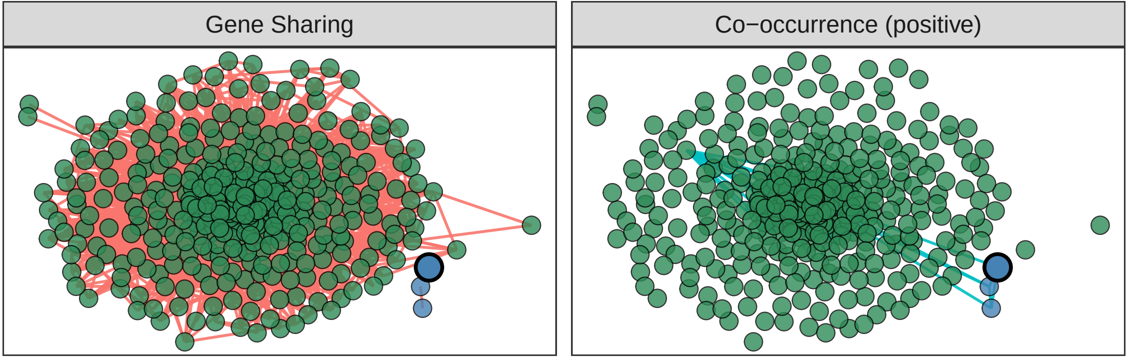

Gene Sharing
Co−occurrence (positive)

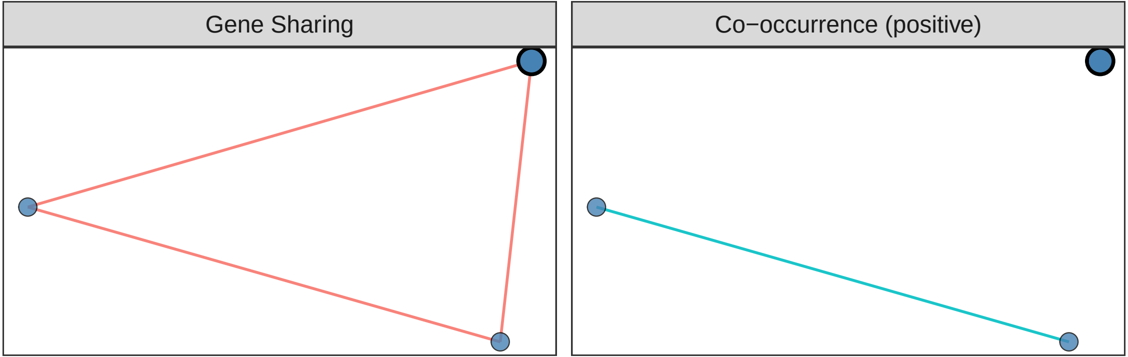
Gene Sharing
Co−occurrence (positive)

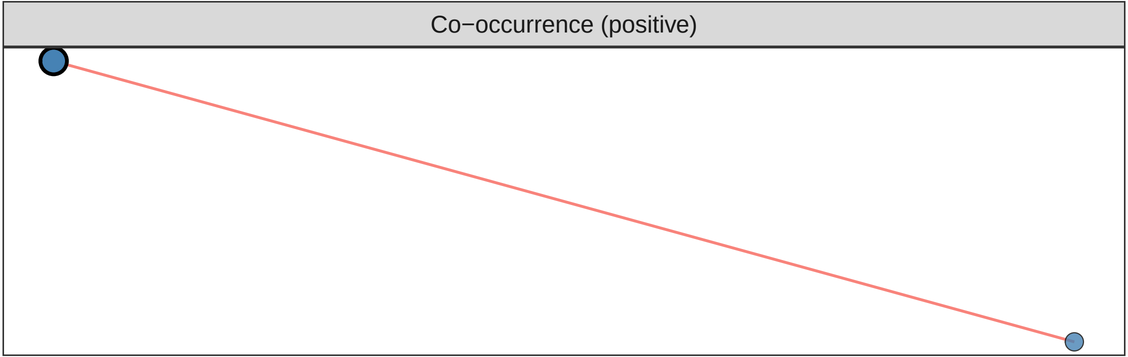
Co−occurrence (positive)

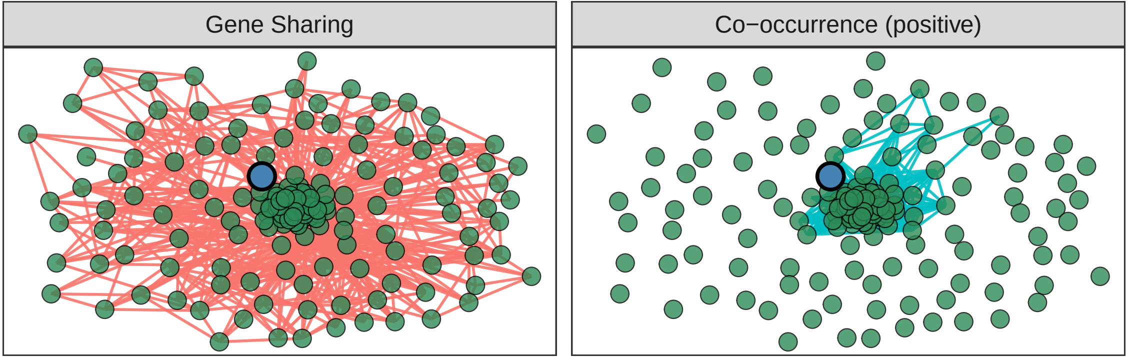
Gene Sharing
Co−occurrence (positive)

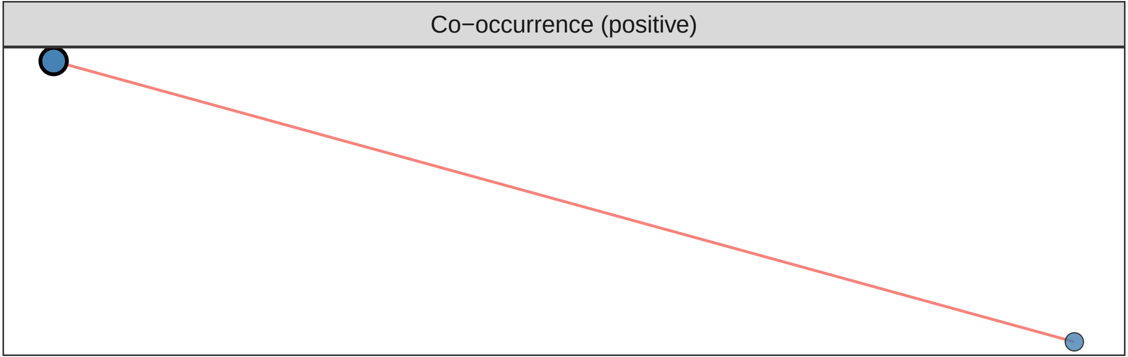
Co−occurrence (positive)

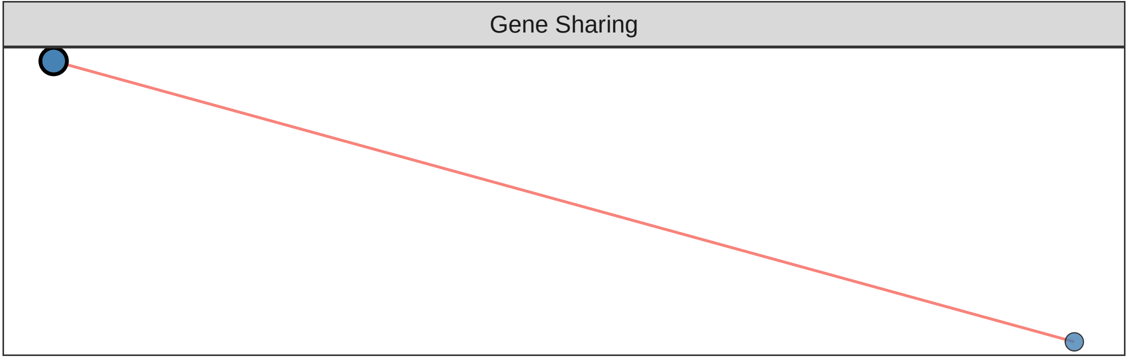
Gene Sharing

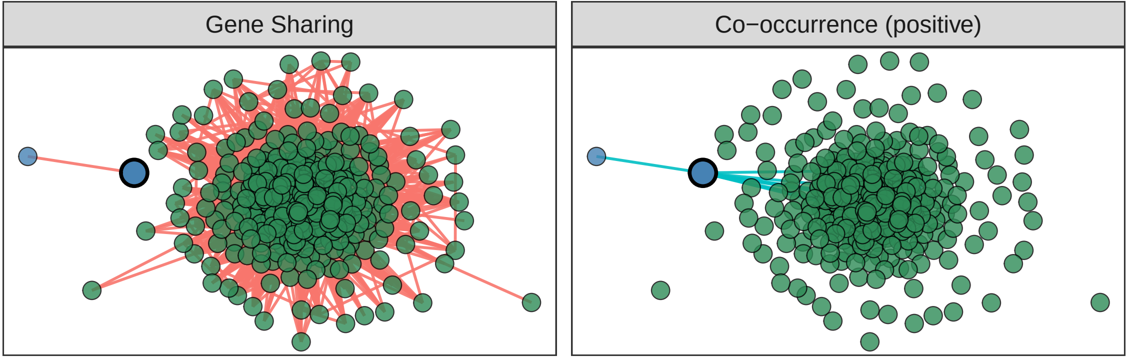
Gene Sharing
Co−occurrence (positive)

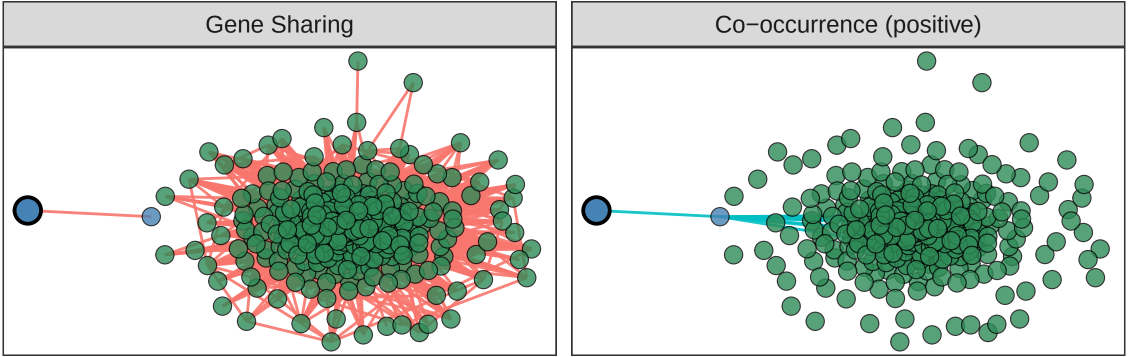
Gene Sharing
Co−occurrence (positive)

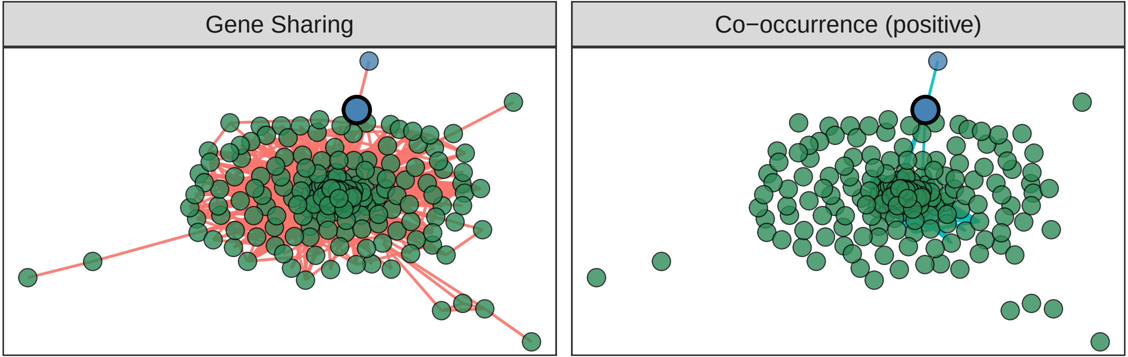
Gene Sharing
Co−occurrence (positive)

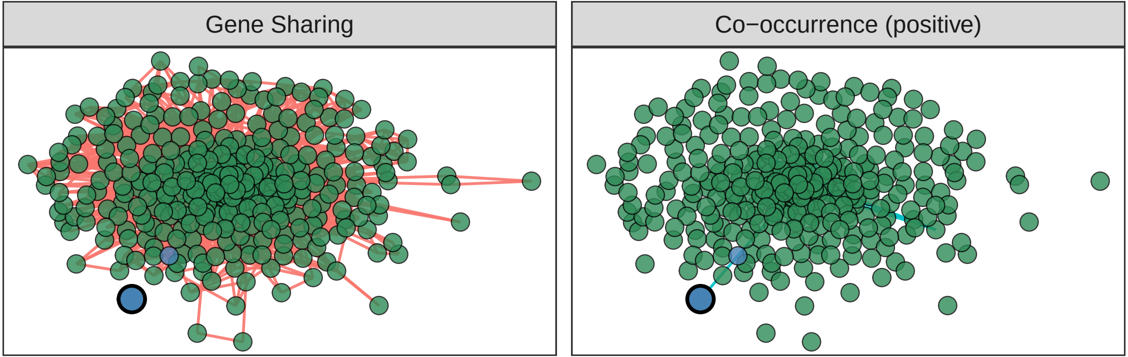
Gene Sharing
Co−occurrence (positive)

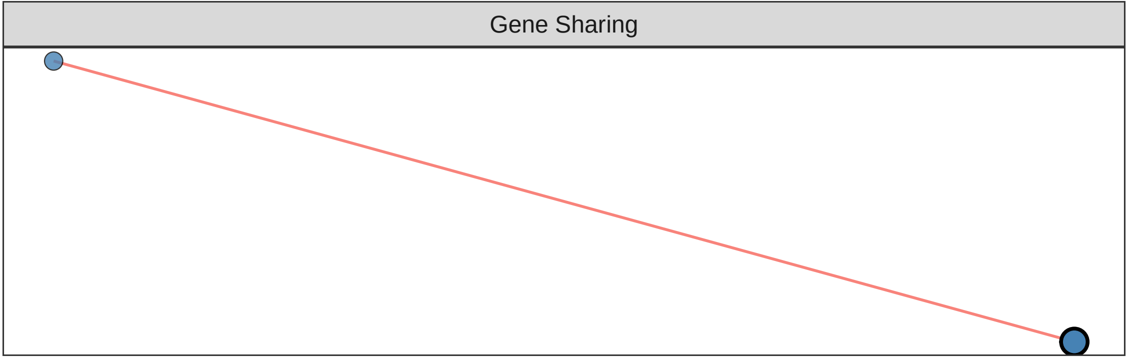
Gene Sharing

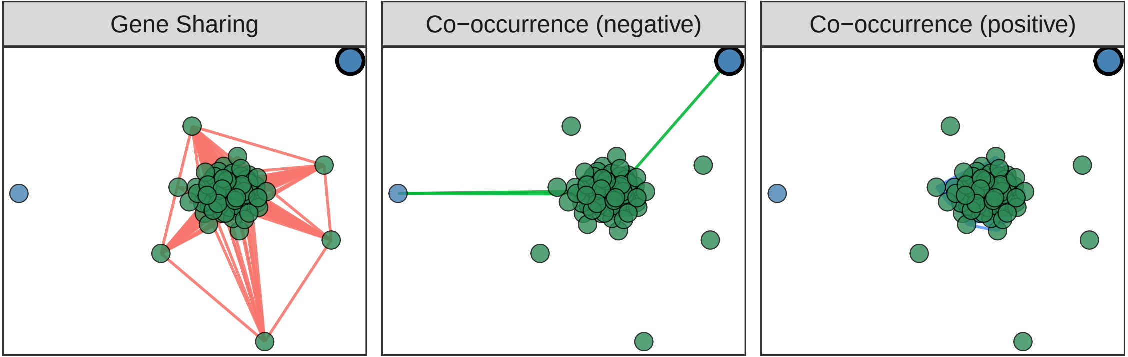
Gene Sharing
Co−occurrence (negative)
Co−occurrence (positive)

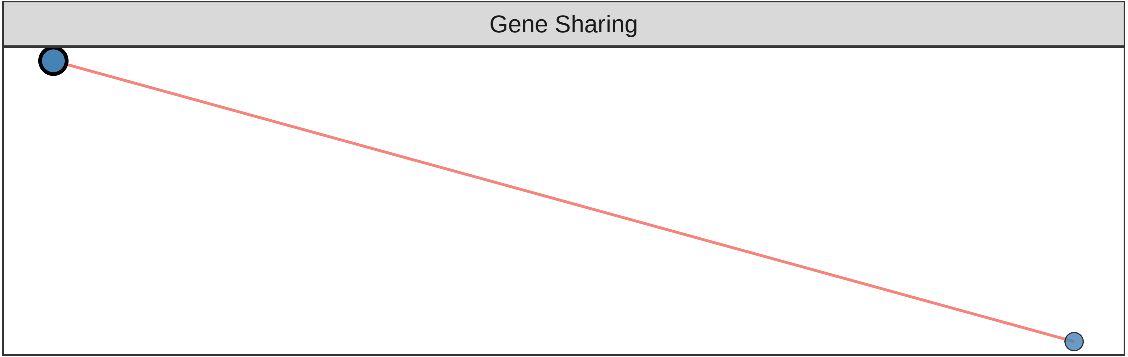
Gene Sharing

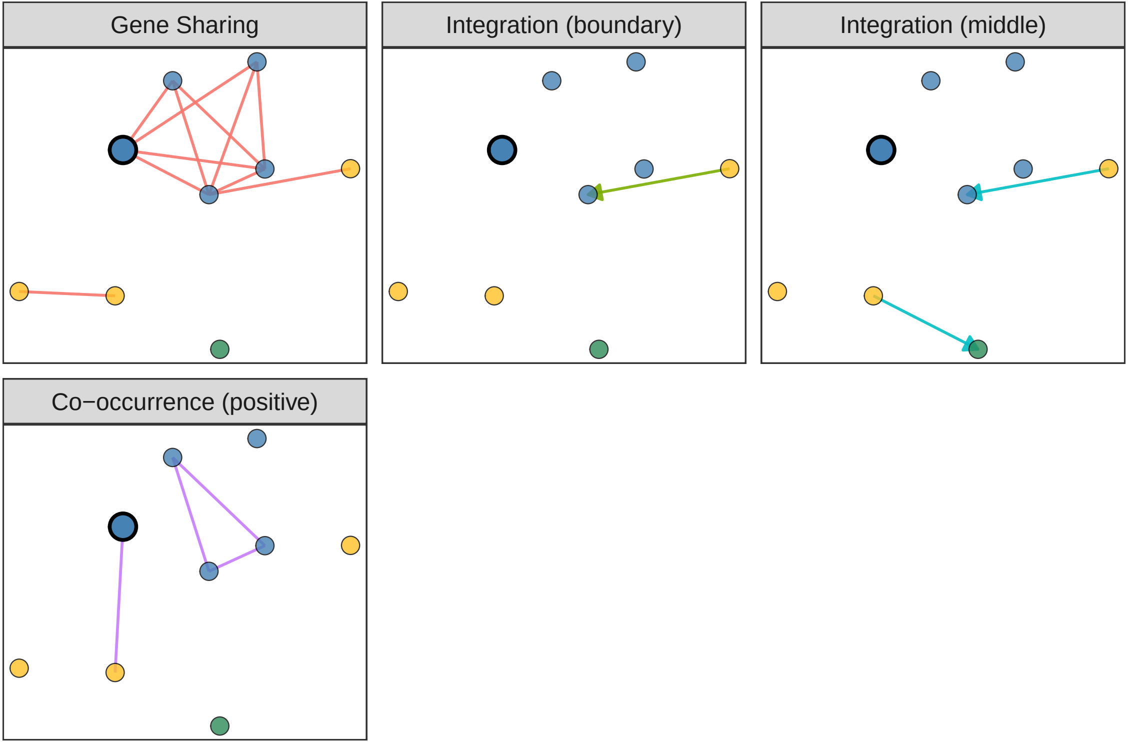
Gene Sharing
Integration (boundary)
Integration (middle)
Co−occurrence (positive)

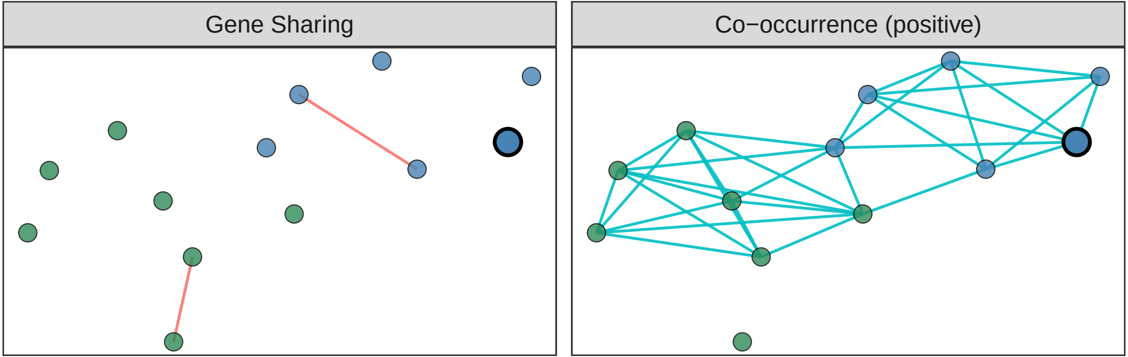
Gene Sharing
Co−occurrence (positive)

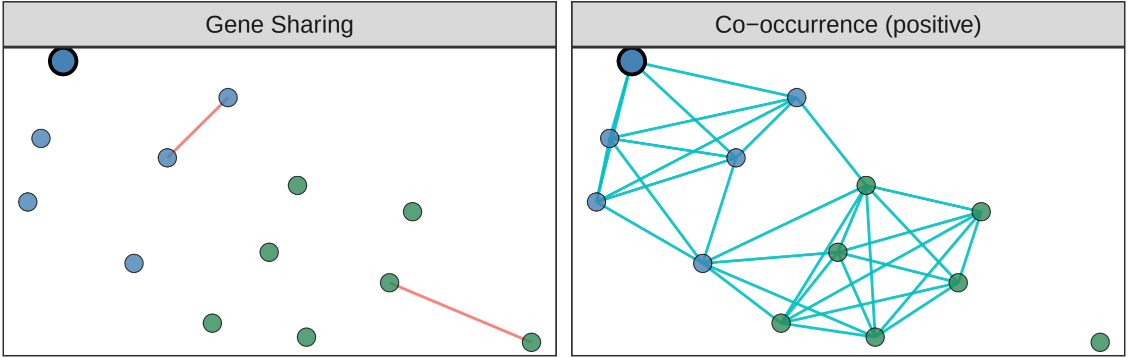

Gene Sharing
Co−occurrence (positive)

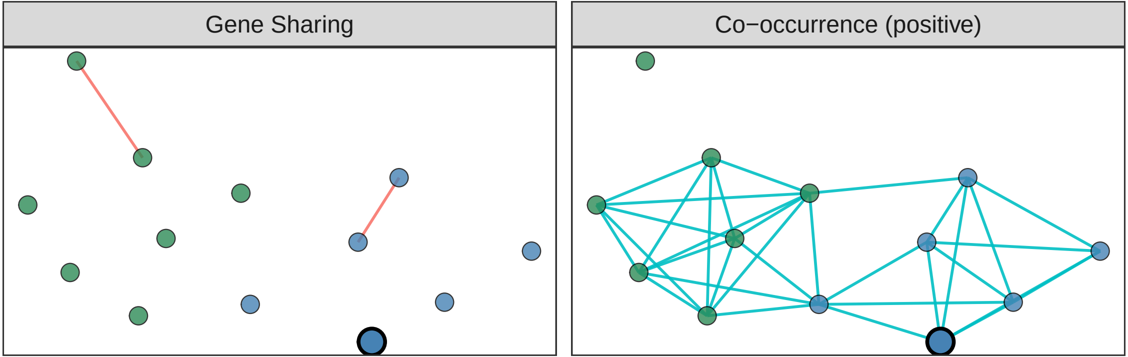
Gene Sharing
Co−occurrence (positive)

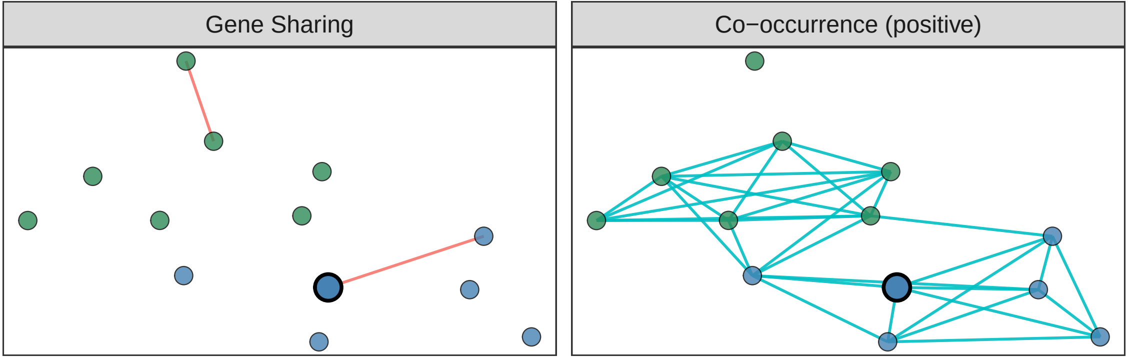
Gene Sharing
Co−occurrence (positive)

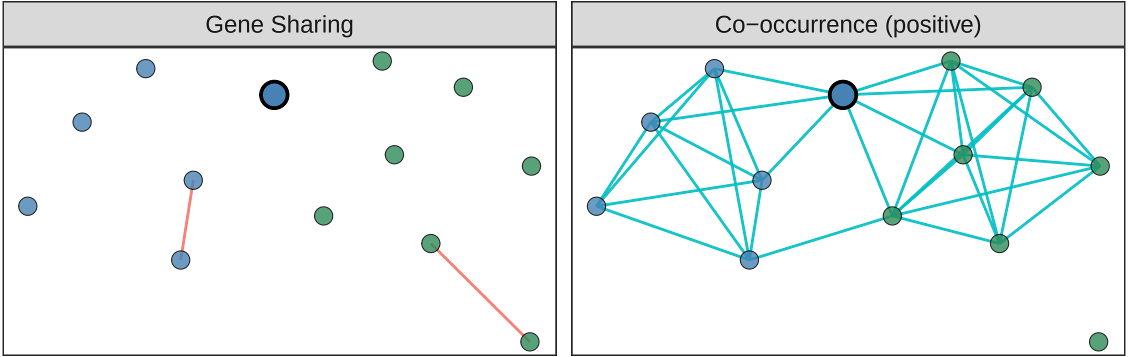
Gene Sharing
Co−occurrence (positive)

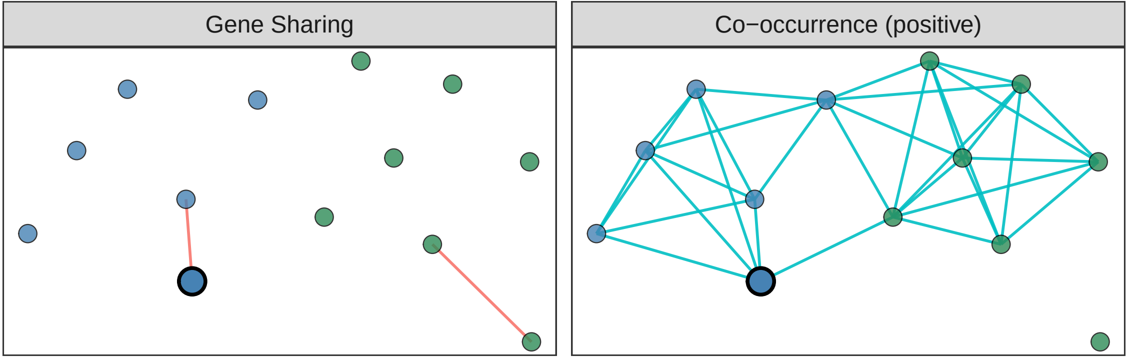

Gene Sharing
Co−occurrence (positive)

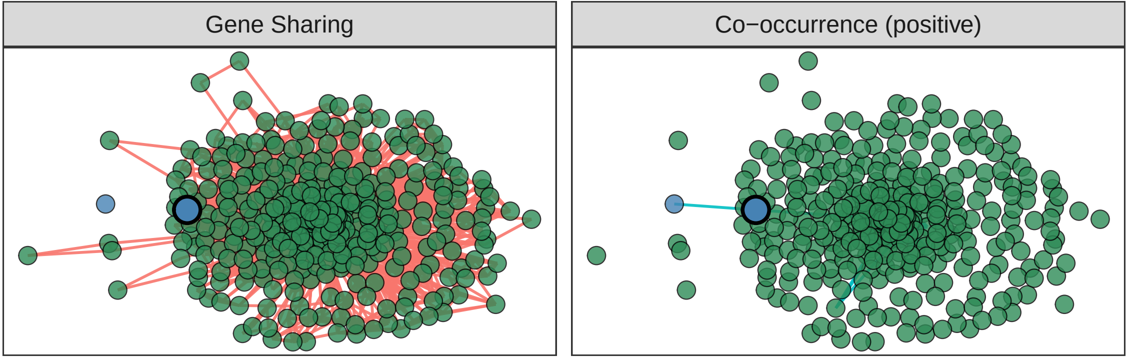
Gene Sharing
Co−occurrence (positive)

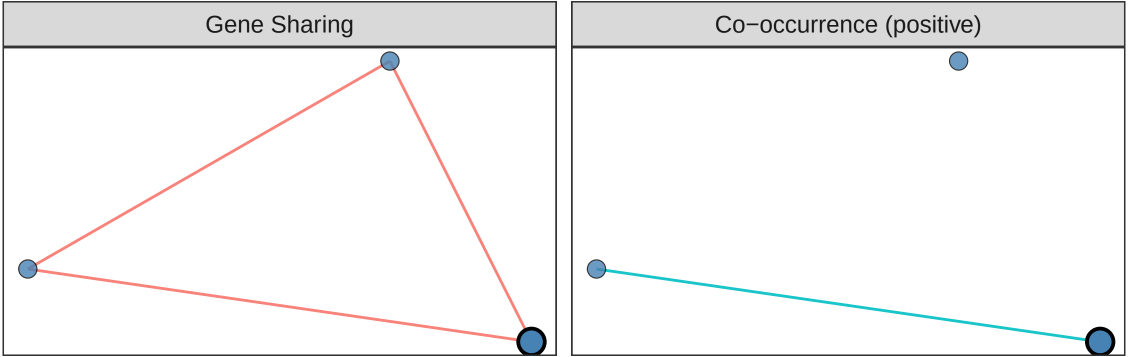
Gene Sharing
Co−occurrence (positive)

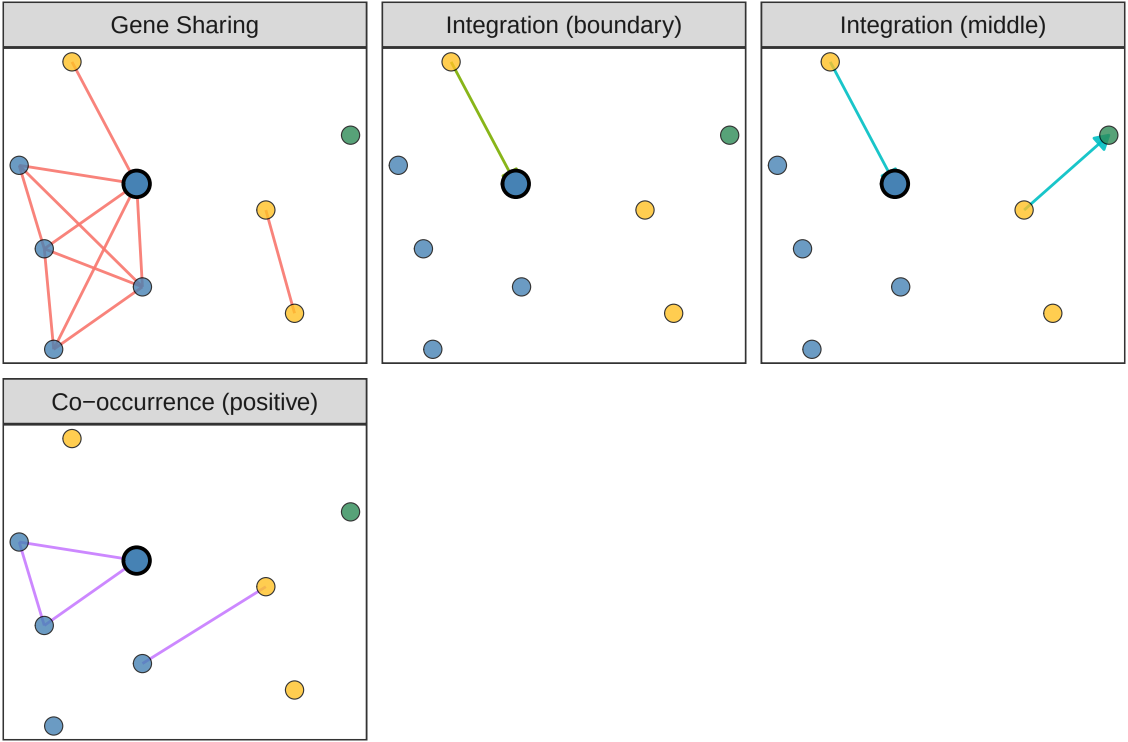
Gene Sharing
Integration (boundary)
Integration (middle)
Co−occurrence (positive)

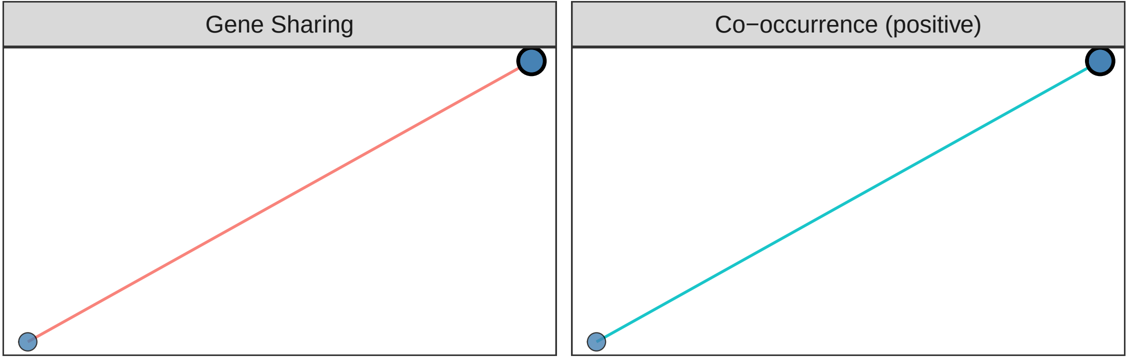
Gene Sharing
Co−occurrence (positive)

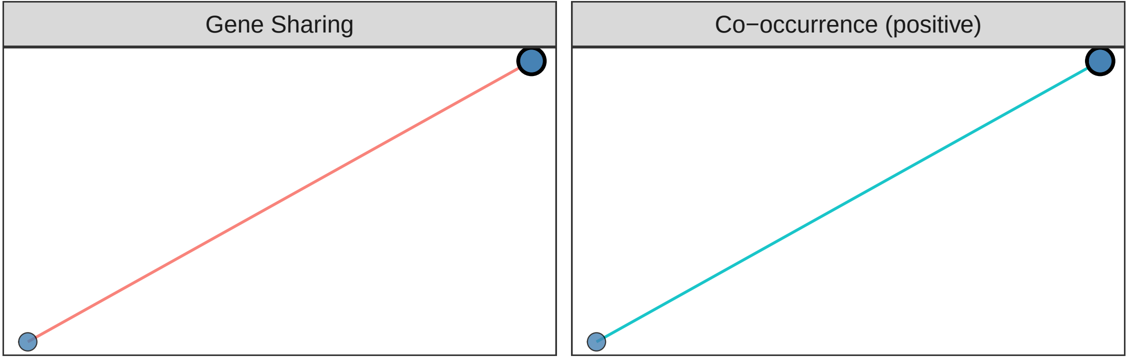
Gene Sharing
Co−occurrence (positive)

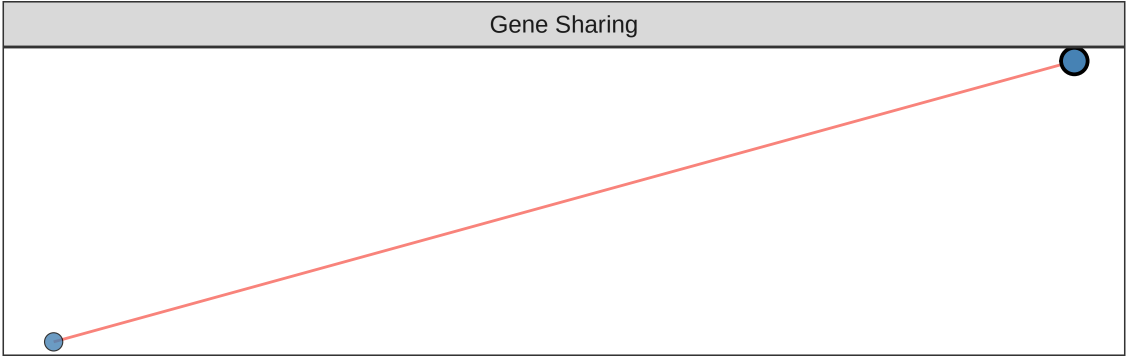
Gene Sharing

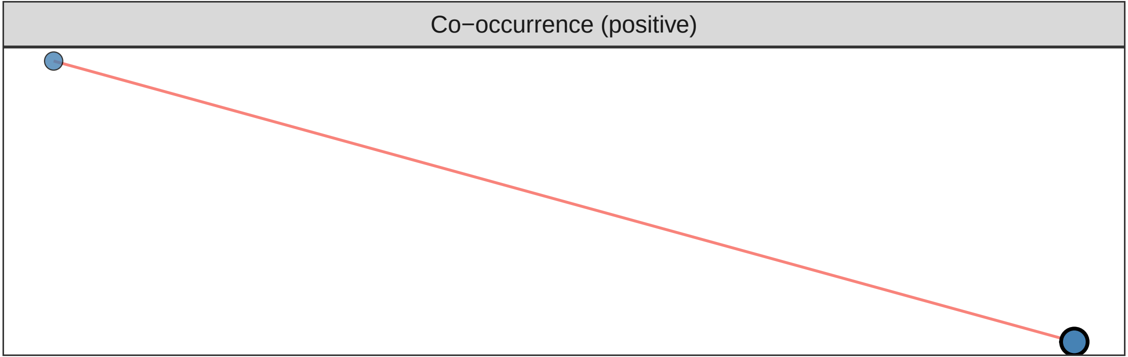
Co−occurrence (positive)

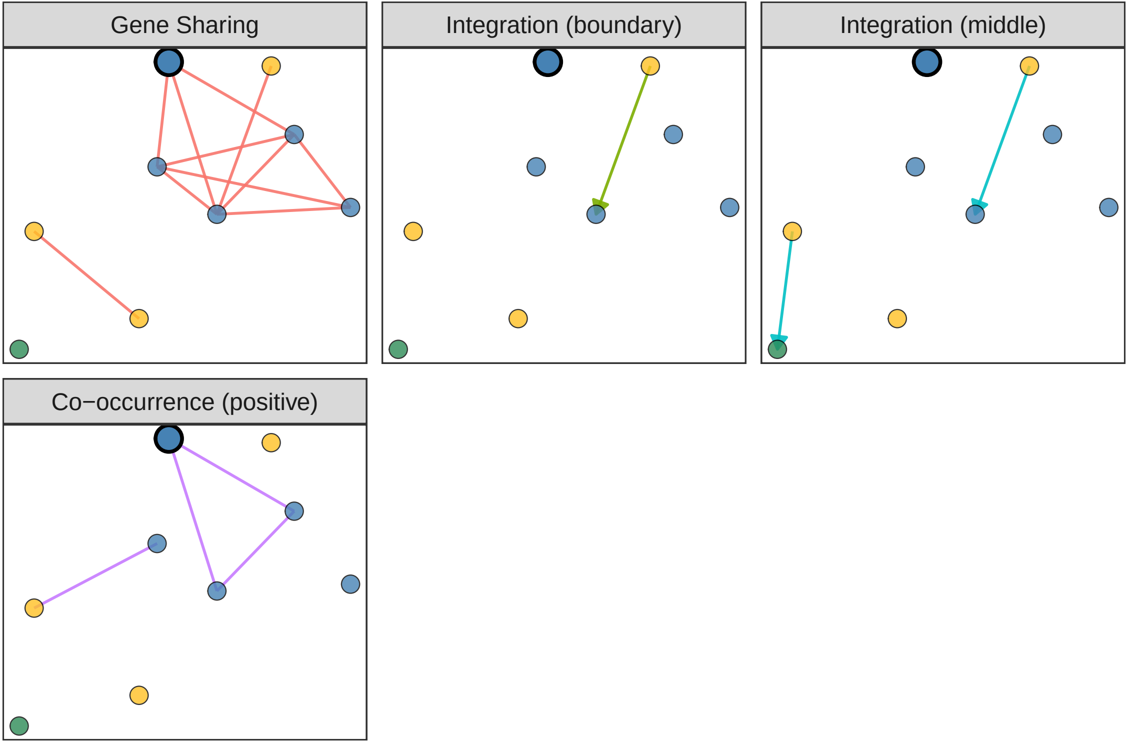
Gene Sharing
Integration (boundary)
Integration (middle)
Co−occurrence (positive)

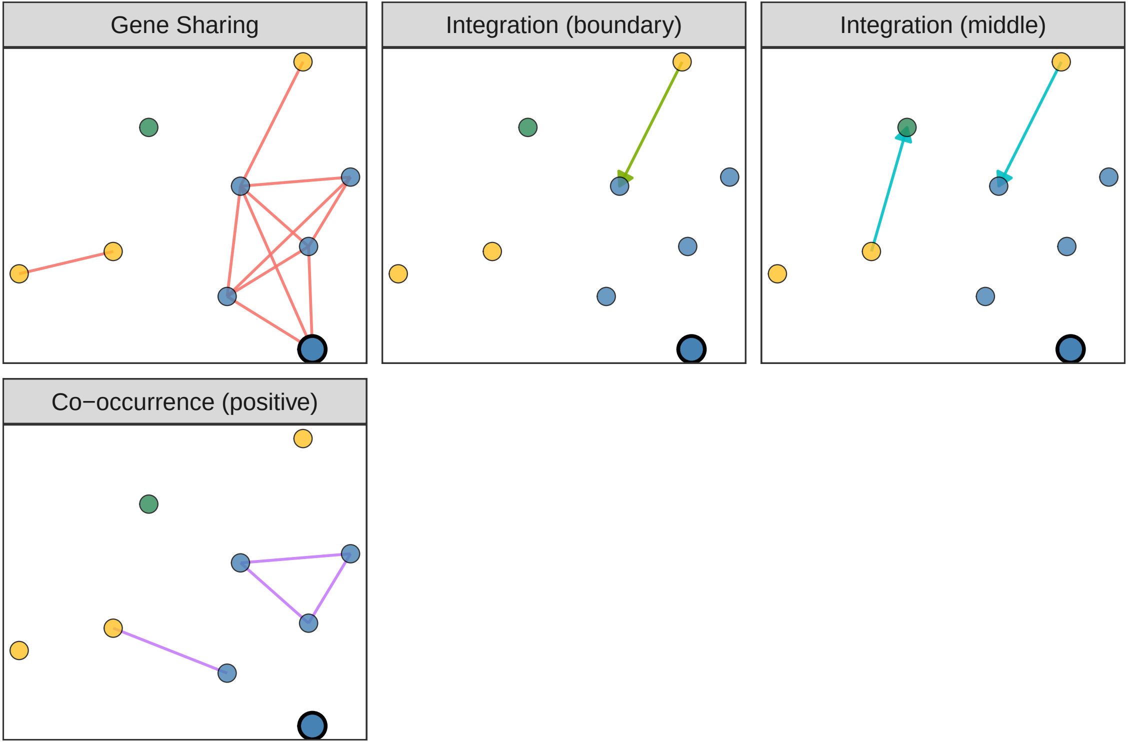
Gene Sharing
Integration (boundary)
Integration (middle)
Co−occurrence (positive)

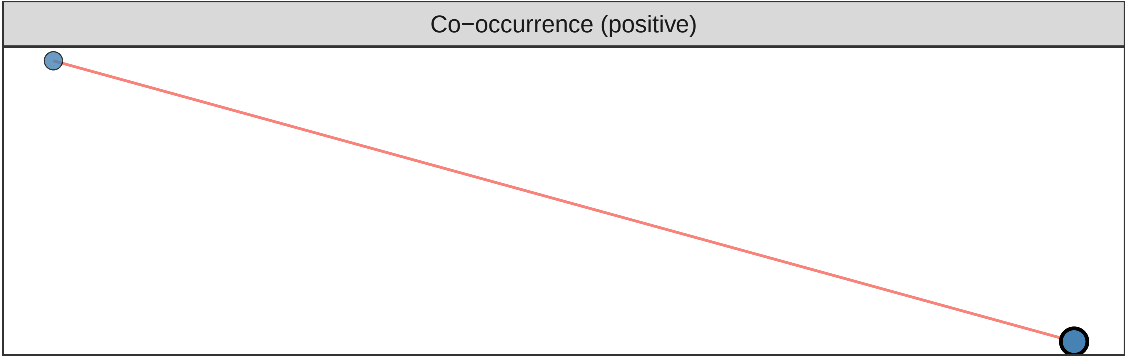
Co−occurrence (positive)

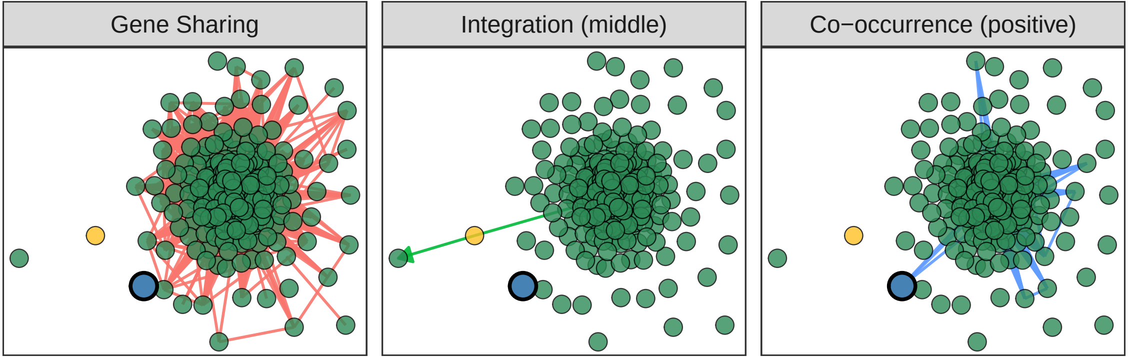
Gene Sharing
Integration (middle)
Co−occurrence (positive)

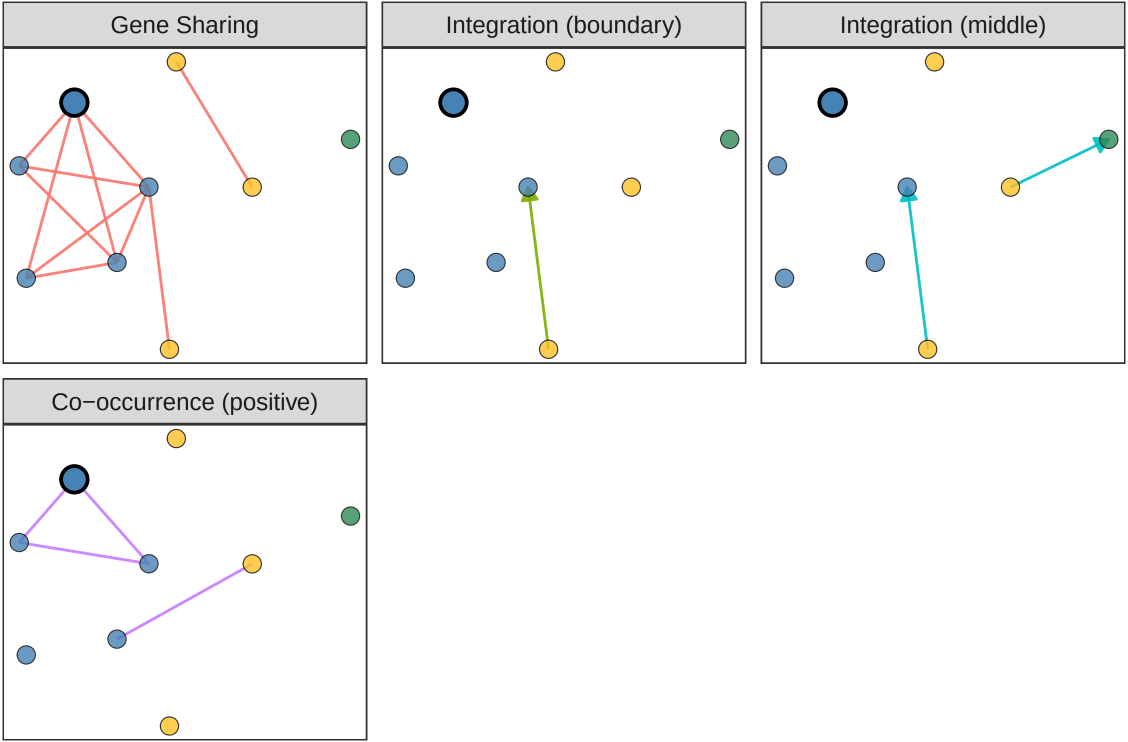
Gene Sharing
Integration (boundary)
Integration (middle)
Co−occurrence (positive)

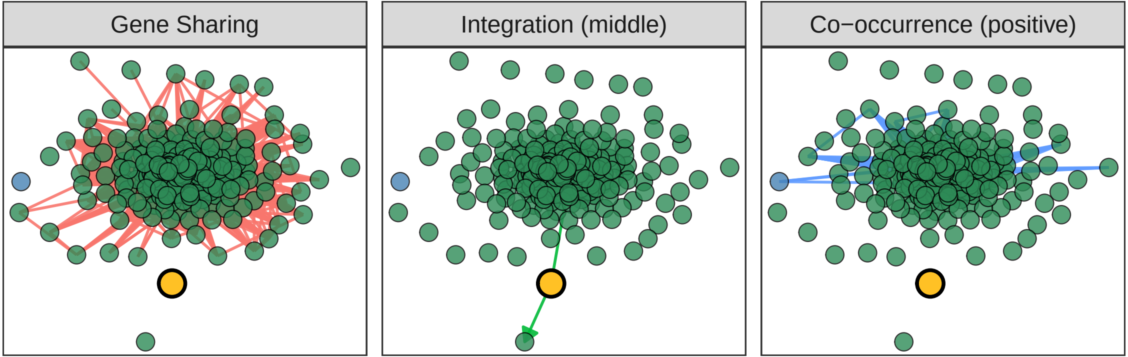
Gene Sharing
Integration (middle)
Co−occurrence (positive)

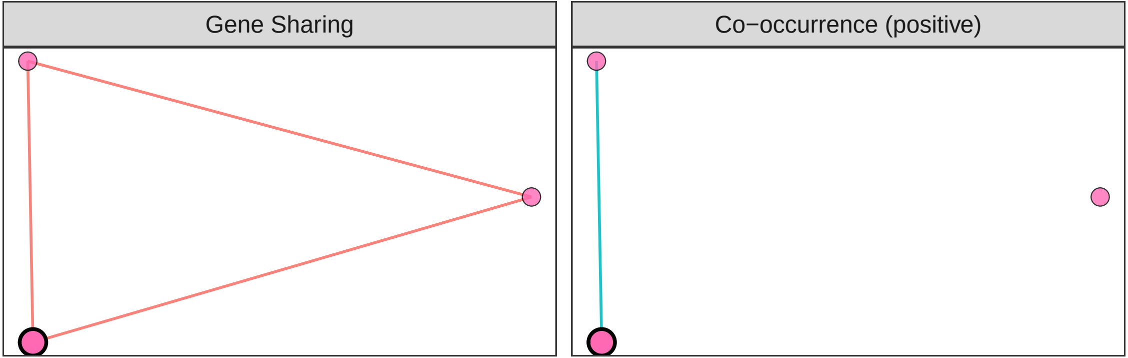
Gene Sharing
Co−occurrence (positive)

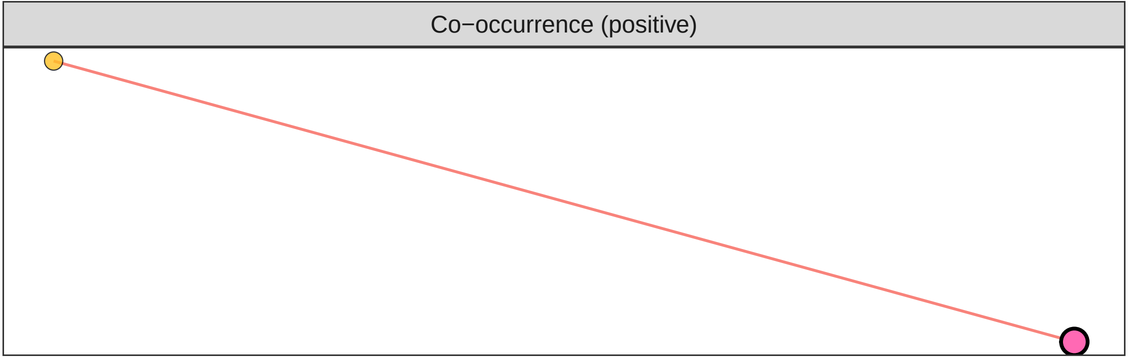
Co−occurrence (positive)

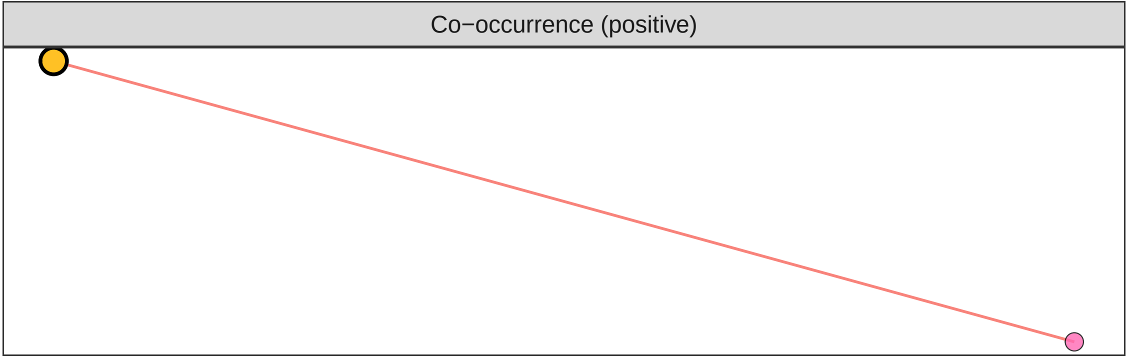
Co−occurrence (positive)

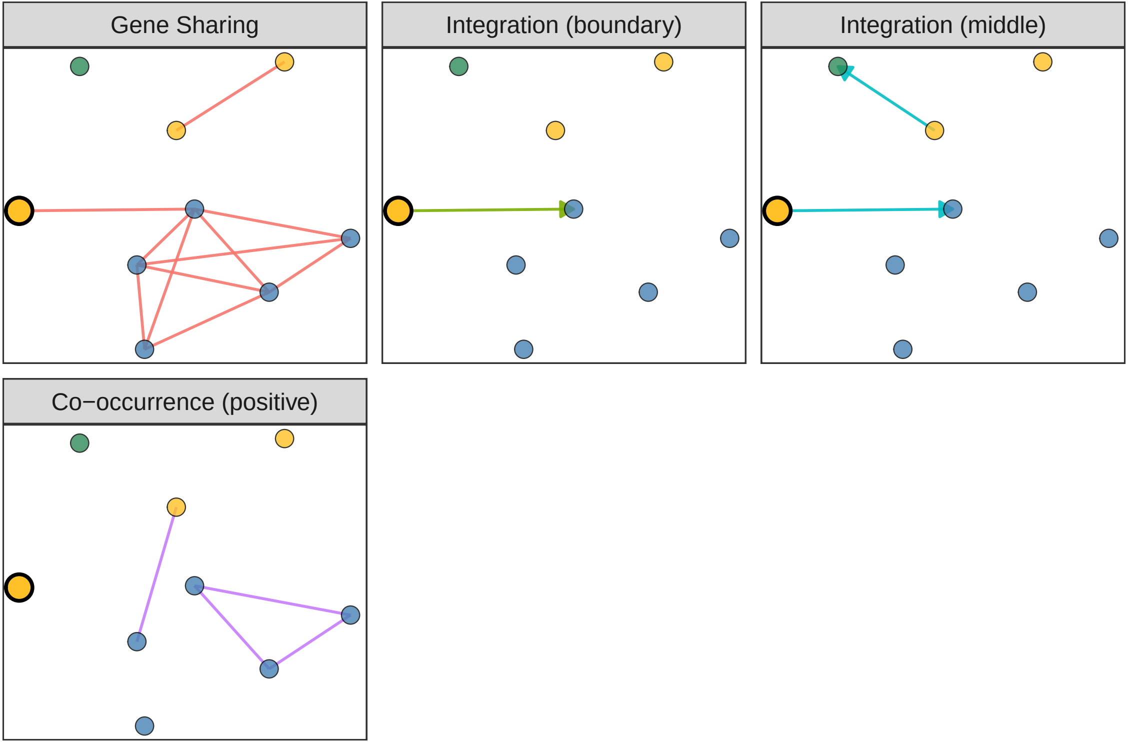
Gene Sharing
Integration (boundary)
Integration (middle)
Co−occurrence (positive)

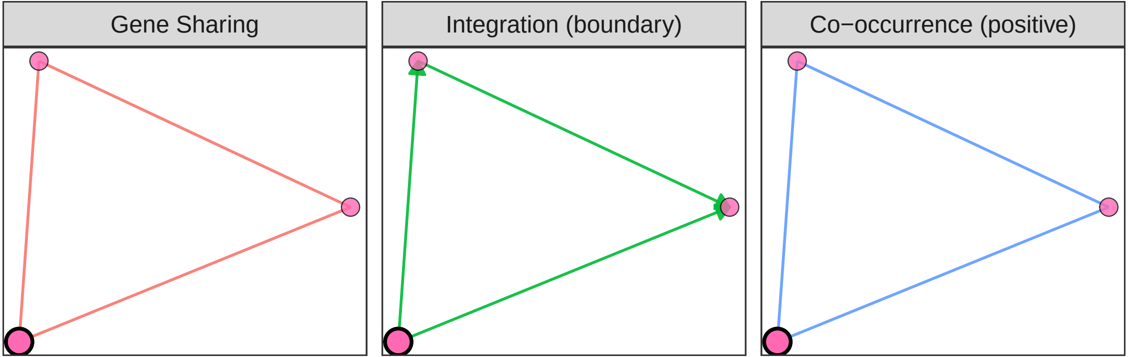
Gene Sharing
Integration (boundary)
Co−occurrence (positive)

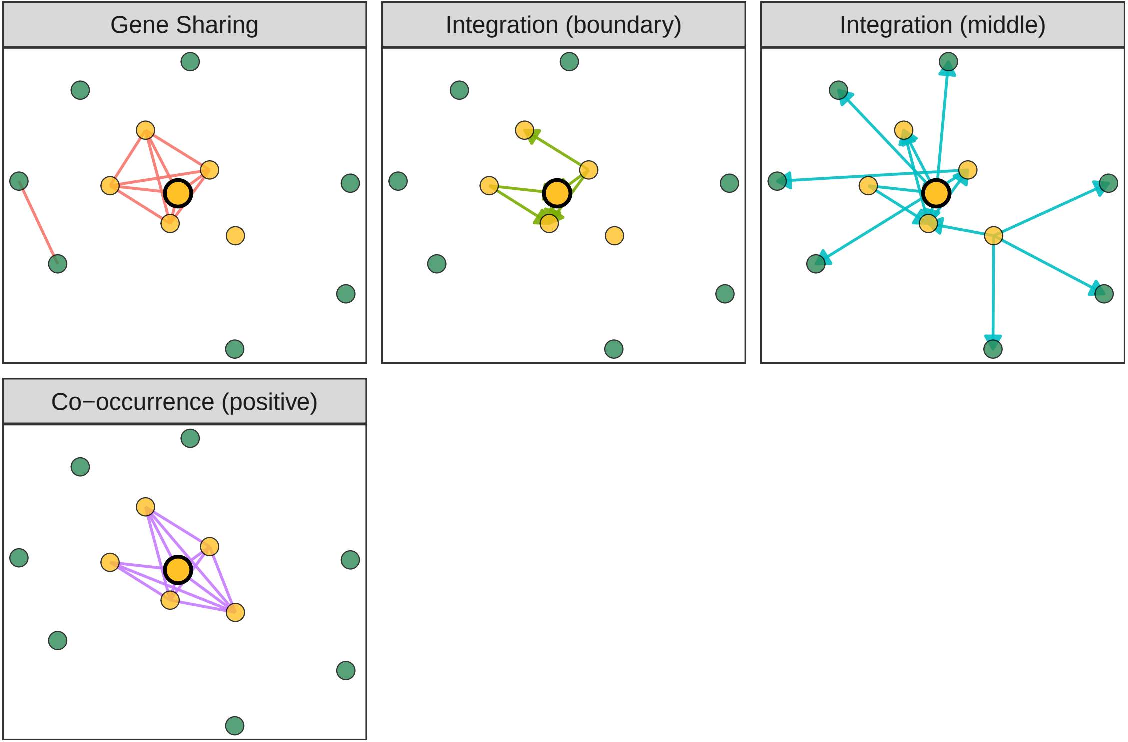
Gene Sharing
Integration (boundary)
Integration (middle)
Co−occurrence (positive)

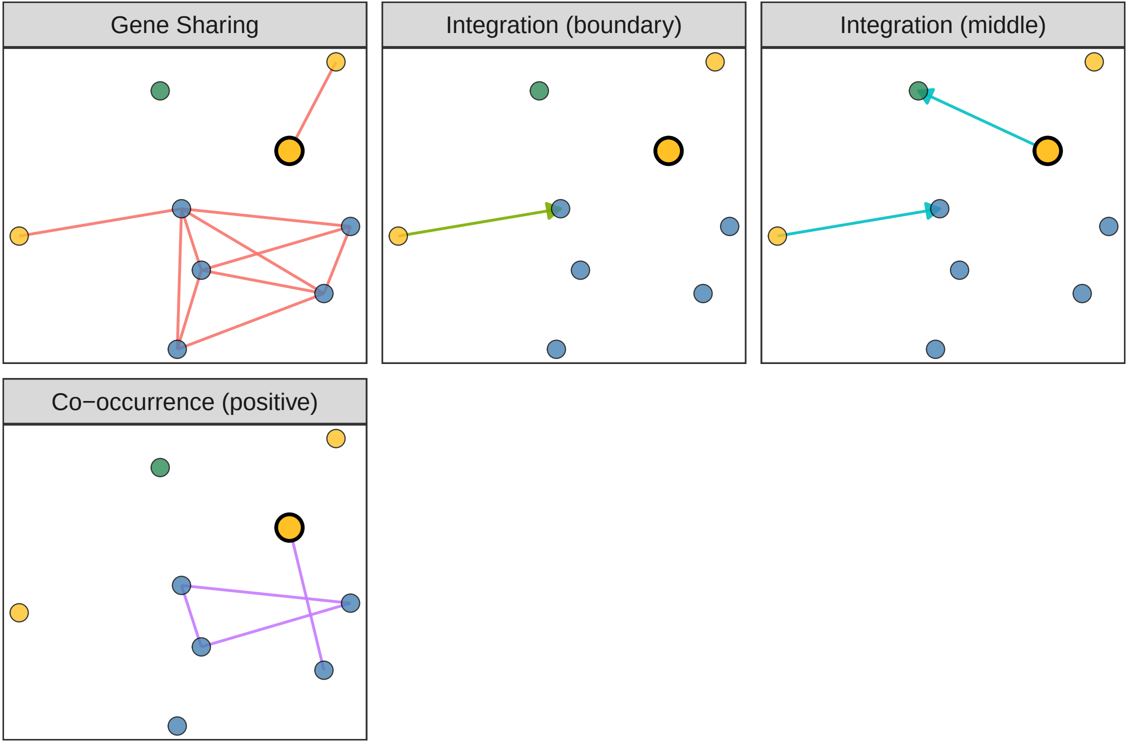
Gene Sharing
Integration (boundary)
Integration (middle)
Co−occurrence (positive)

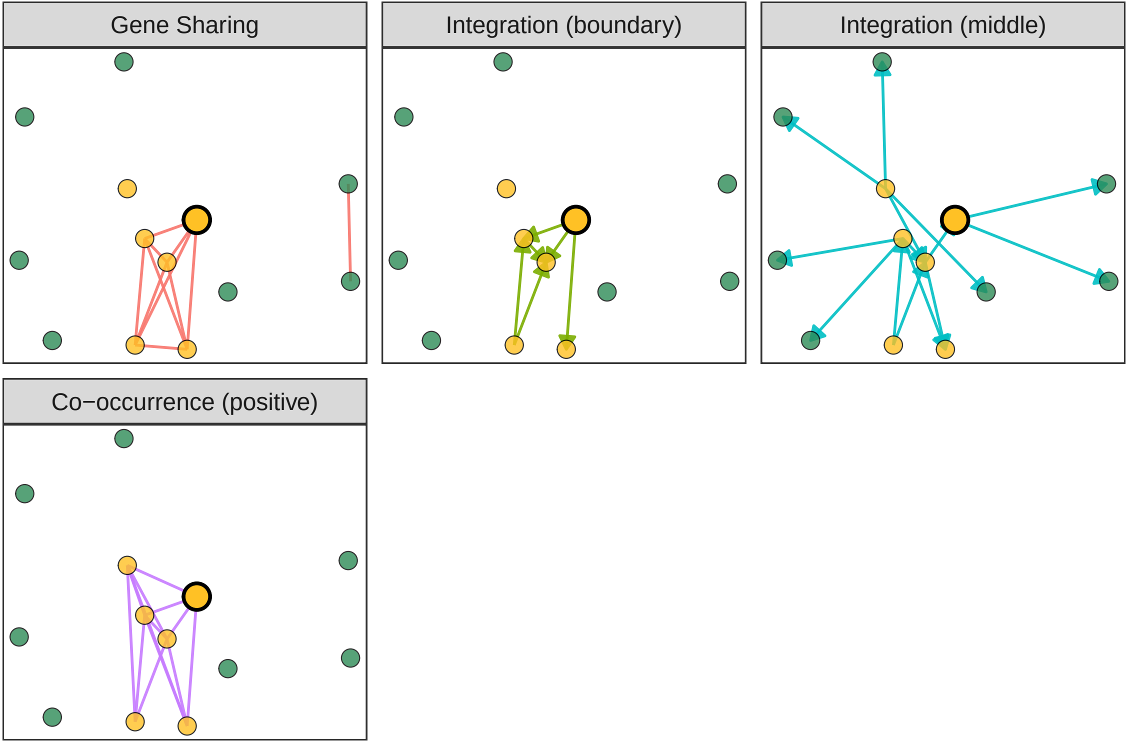
Gene Sharing
Integration (boundary)
Integration (middle)
Co−occurrence (positive)

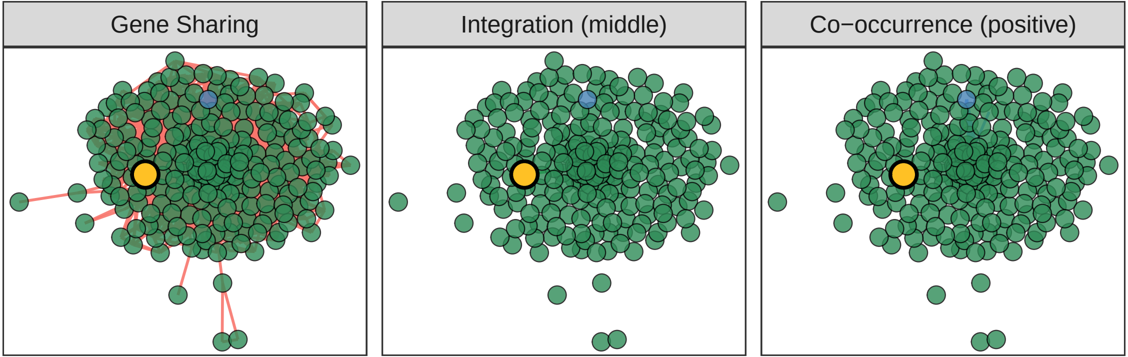
Gene Sharing
Integration (middle)
Co−occurrence (positive)

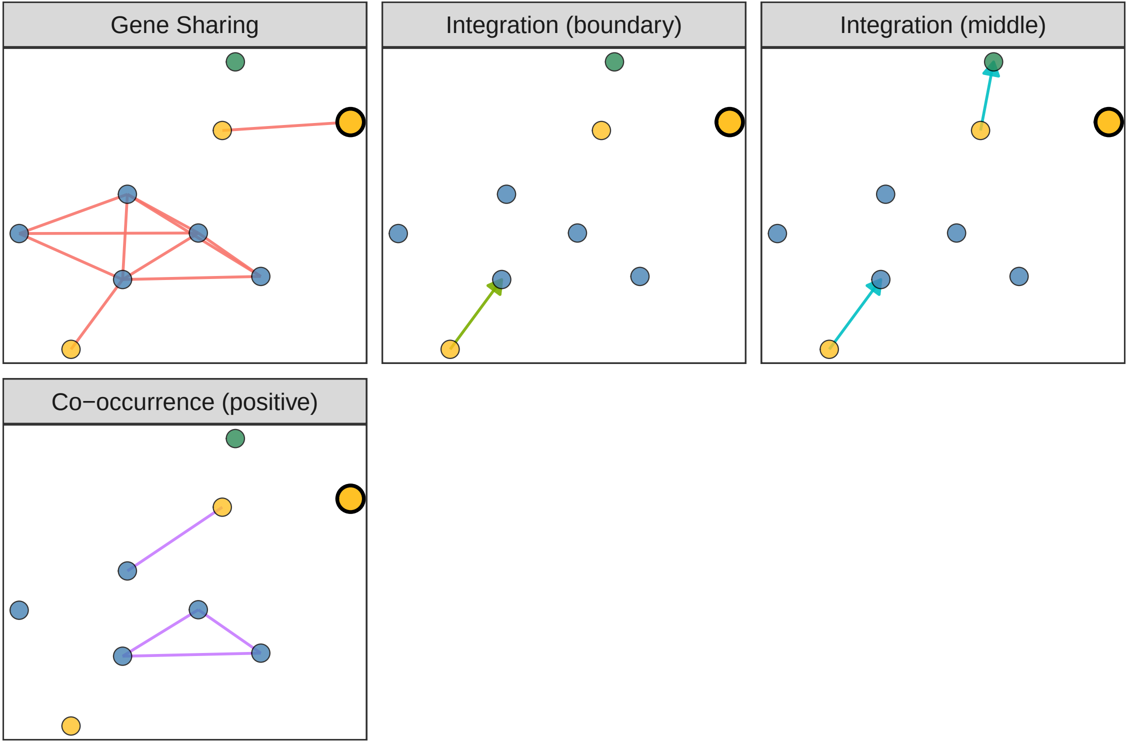
Gene Sharing
Integration (boundary)
Integration (middle)
Co−occurrence (positive)

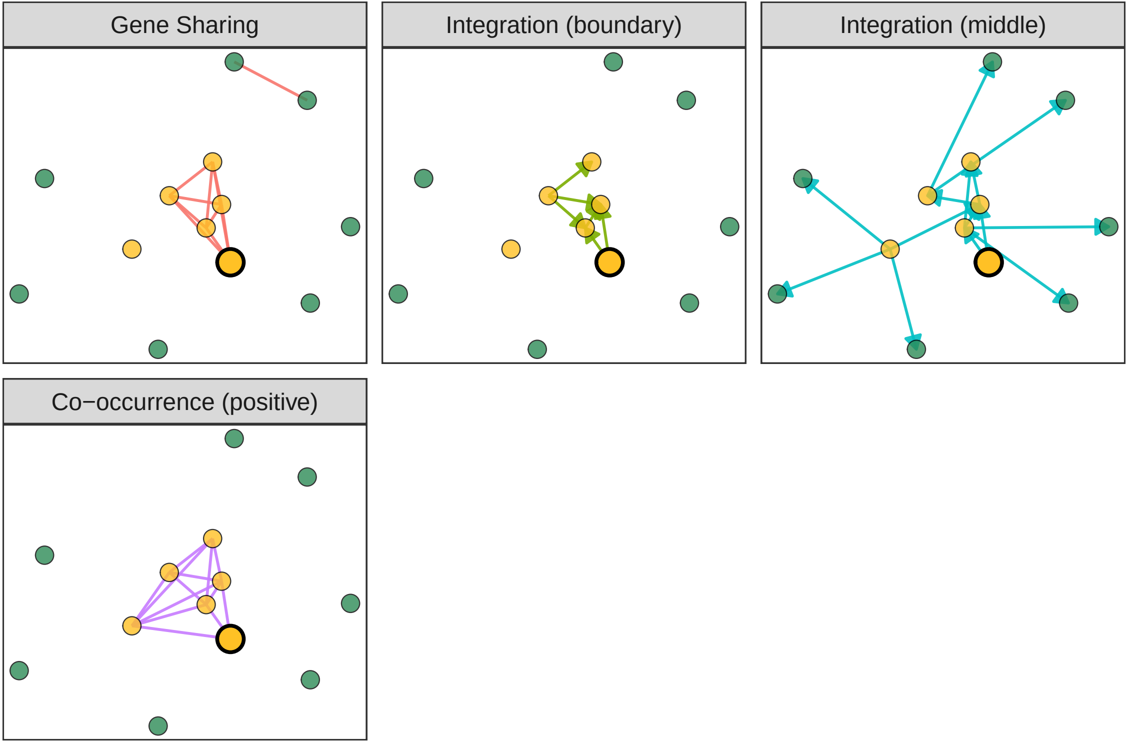
Gene Sharing
Integration (boundary)
Integration (middle)
Co−occurrence (positive)

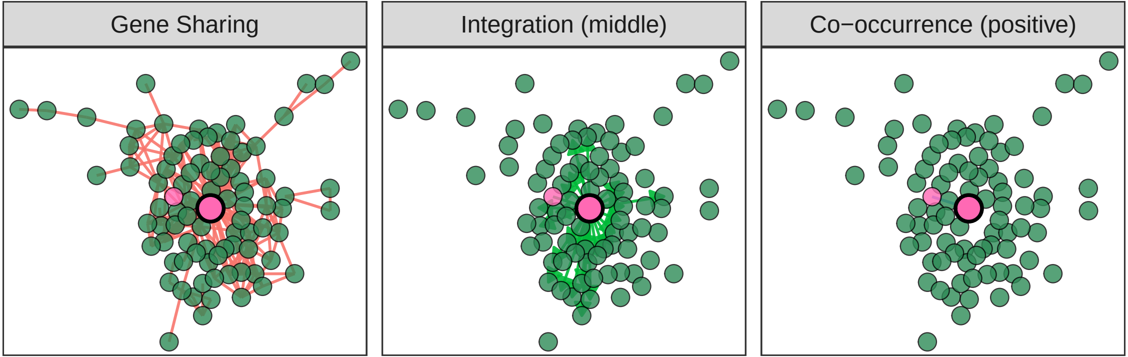
Gene Sharing
Integration (middle)
Co−occurrence (positive)

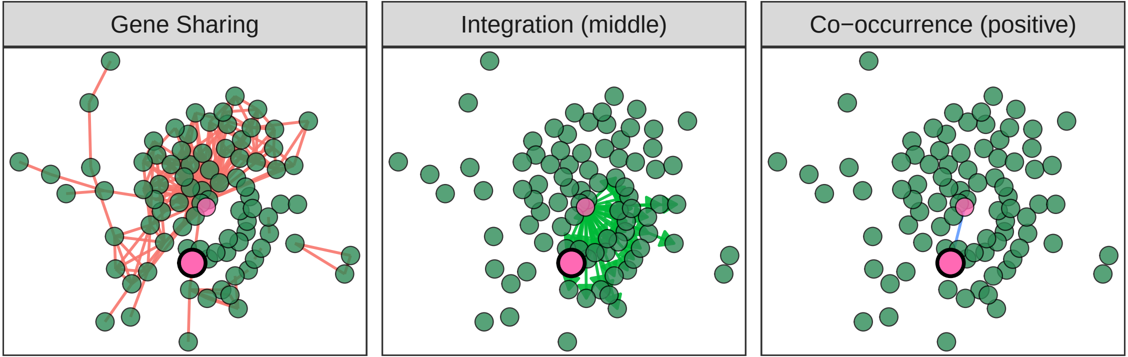
Gene Sharing
Integration (middle)
Co−occurrence (positive)

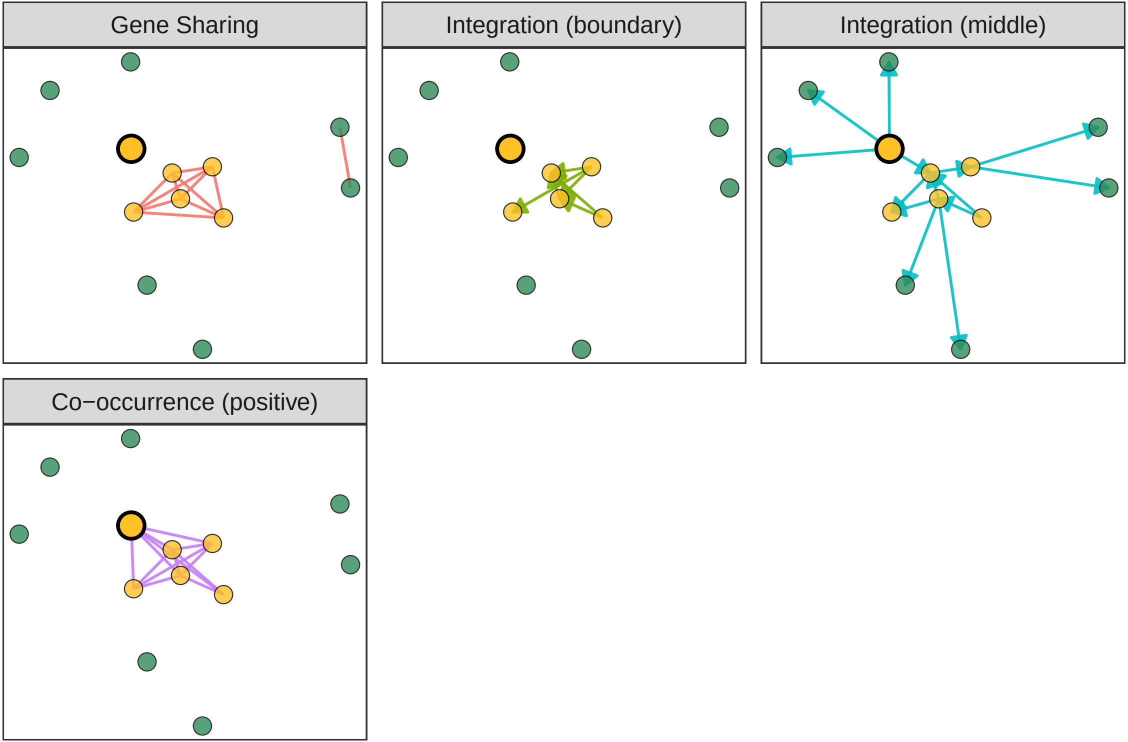
Gene Sharing
Integration (boundary)
Integration (middle)
Co−occurrence (positive)

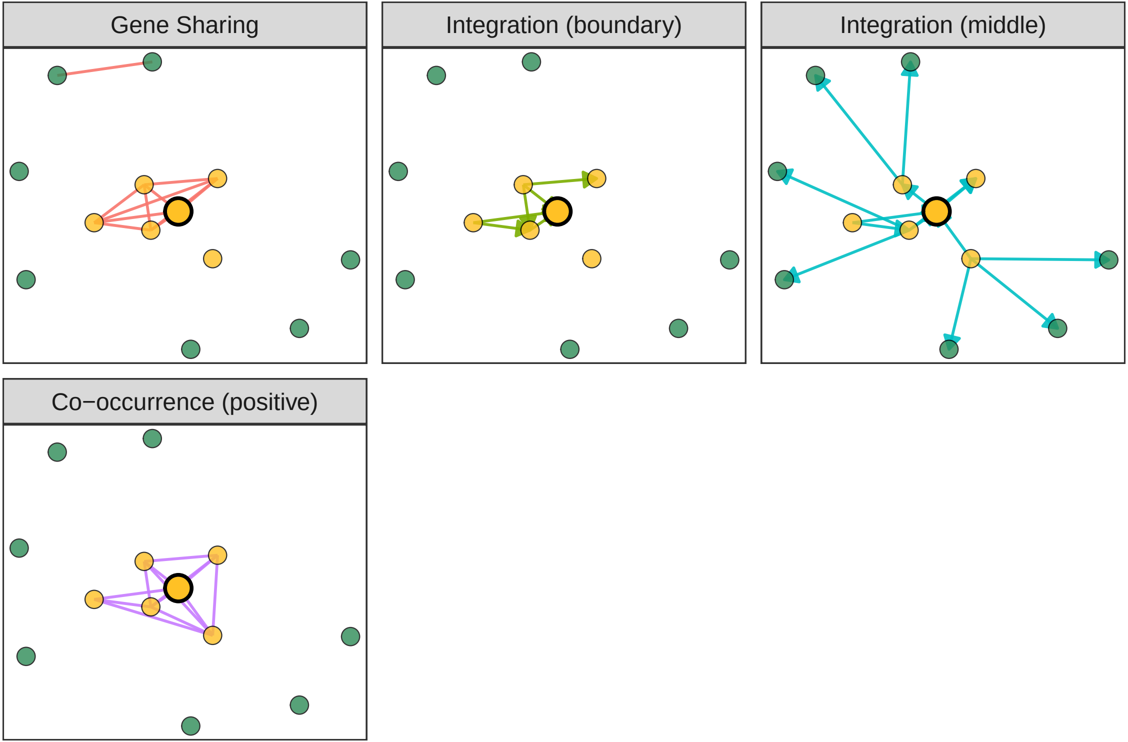
Gene Sharing
Integration (boundary)
Integration (middle)
Co−occurrence (positive)

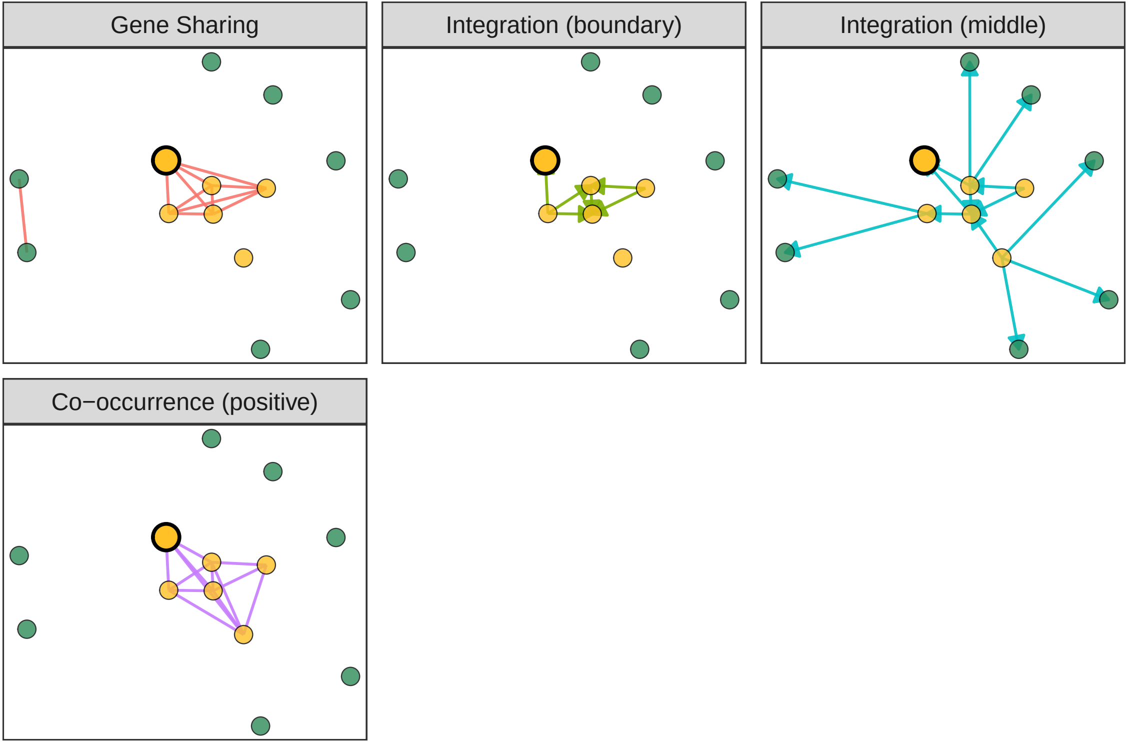
Gene Sharing
Integration (boundary)
Integration (middle)
Co−occurrence (positive)

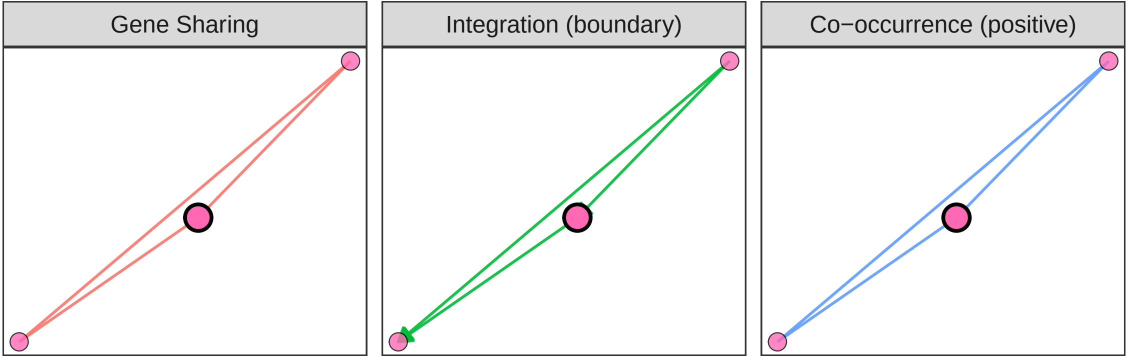
Gene Sharing
Integration (boundary)
Co−occurrence (positive)

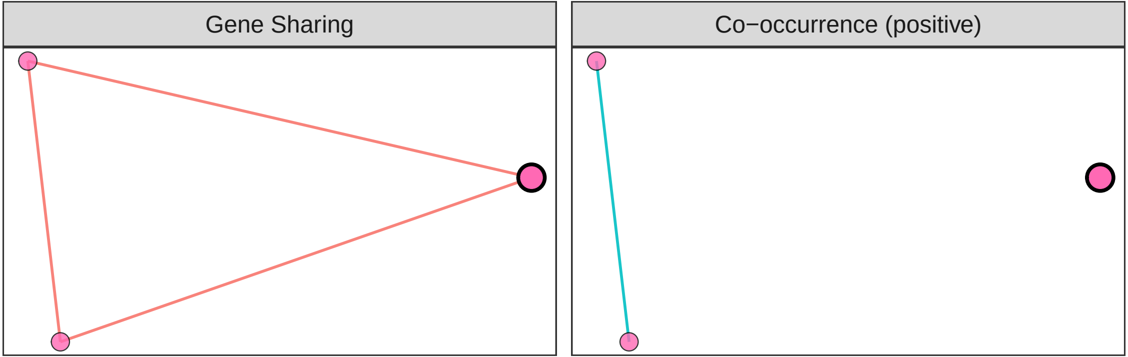
Gene Sharing
Co−occurrence (positive)

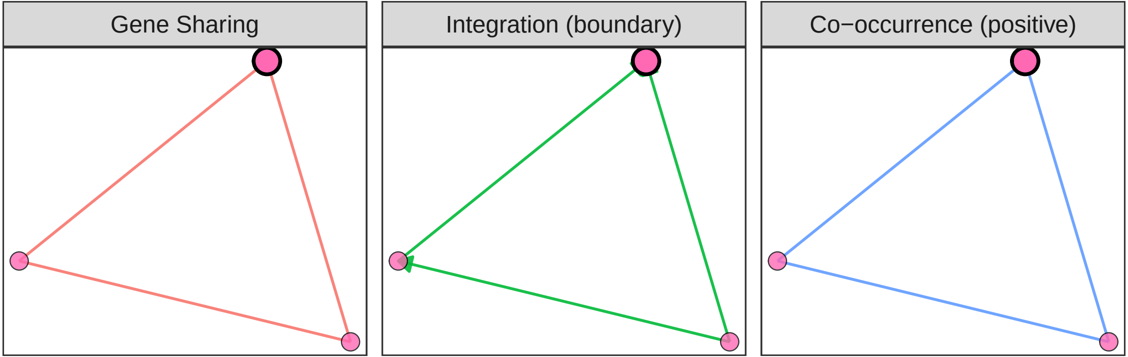
Gene Sharing
Integration (boundary)
Co−occurrence (positive)

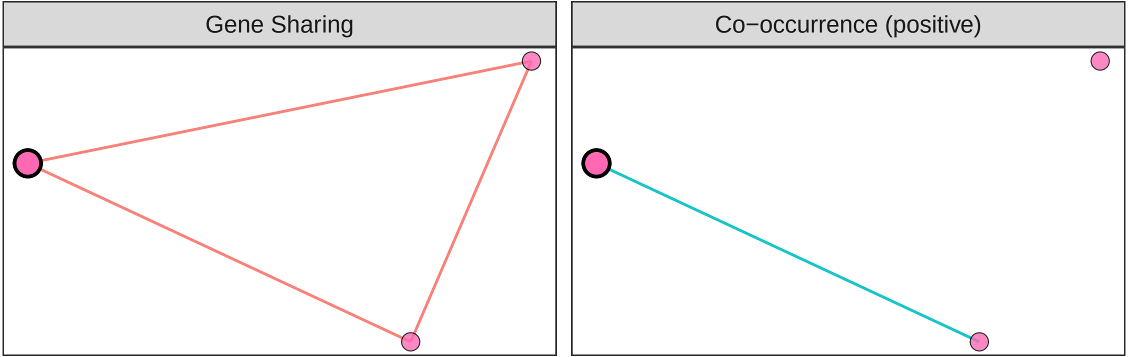
Gene Sharing
Co−occurrence (positive)

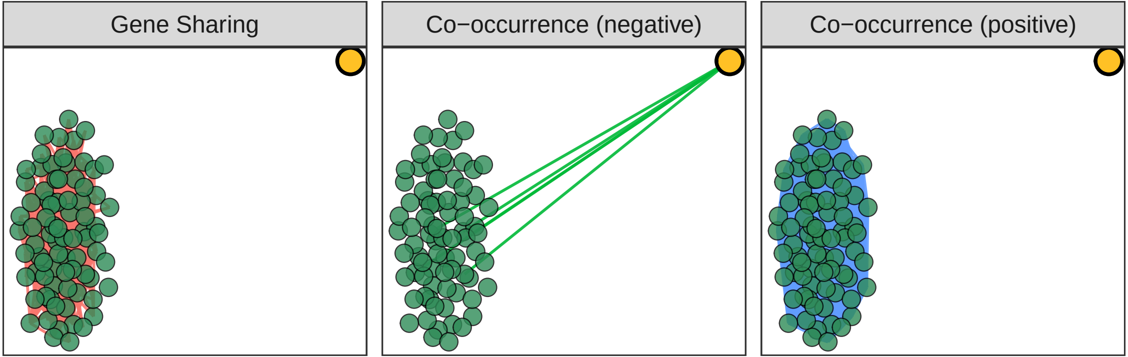
Gene Sharing
Co−occurrence (negative)
Co−occurrence (positive)
